# Multiome Profiling Reveals Astrocyte and Neuroendocrine Targets of Prenatal Acoustic Programming in Zebra Finch Embryos

**DOI:** 10.64898/2026.06.01.729058

**Authors:** Prakrit Subba, Vijay Shankar, Kaitlyn Williams, Trudy F. C. Mackay, Patrick S. Freymuth, Natalie A. Shay, Allison M. Rees, Shannon R. Liedl, Jonathan Roberts, David F. Clayton, Julia M. George

## Abstract

Climate change is driving more frequent and severe heat events, yet the developmental mechanisms by which organisms anticipate and adapt to thermal stress remain poorly understood. In zebra finches (*Taeniopygia guttata*), parents exposed to extreme heat emit distinctive “heat call” vocalizations during late incubation that trigger adaptive phenotypic responses in offspring, including altered growth, thermoregulation, and reproductive success. Recent transcriptomic analysis revealed that prenatal heat call exposure induces hypothalamic gene expression changes enriched in non-neuronal populations, but cellular heterogeneity has obscured which cell types are primary targets and the regulatory mechanisms underlying acoustic programming. Here, we performed single-nucleus multiome sequencing to generate a cell-type-resolved atlas of gene expression and chromatin accessibility in the late-stage embryonic zebra finch hypothalamus. We find that heat call exposure selectively remodels astrocyte regulatory landscapes, with most playback-responsive chromatin changes localized to glial populations like astrocytes, challenging neuron-centric models of developmental programming. Motif enrichment and gene regulatory network analysis showed Nuclear Factor I-C, AP-1, and Notch pathway components - canonical drivers of the gliogenic switch - coordinate this response. Chromatin co-accessibility analysis revealed extensive astrocyte-specific enhancer-promoter rewiring at developmental signaling hubs, and pseudotemporal analysis identified condition-associated shifts in developmental trajectories and astrocyte gene-expression dynamics consistent with altered gliogenic programs. In parallel, we identified sex-dimorphic regulation of transthyretin in glutamatergic neurons of the paraventricular nucleus, linking acoustic programming to thyroid-dependent neuroendocrine pathways. These findings suggest that prenatal acoustic experience engages astrocyte-selective epigenetic priming and neuroendocrine transcriptional responses that may reshape hypothalamic developmental programs under predicted thermal environments.

## 1 Introduction

Early-life sensory experiences during critical developmental windows shape neural circuits and function with lasting consequences for physiology and behavior (Molnár et al., 2020). Recent evidence reveals that prenatal sensory input can exert instructive effects on developing circuits before the onset of conventional postnatal critical periods (Roy et al., 2020), raising the possibility that embryonic experience could adaptively program offspring for predicted environmental conditions. The hypothalamus, which coordinates physiological and behavioral responses to environmental challenges, undergoes developmental trajectories that are strongly shaped by early-life experience (Gali Ramamoorthy et al., 2015; Kaplan et al., 2025; Korosi and Baram, 2010; Spencer, 2013). In zebra finches (*Taeniopygia guttata*), parents produce characteristic “heat call” vocalizations under hot weather conditions, and late-stage embryonic exposure to these heat calls programs offspring toward heat-adapted phenotypes, including altered growth trajectories, begging behavior, and mitochondrial function (Mariette and Buchanan, 2016; Pessato et al., 2022; Udino and Mariette, 2022; Udino et al., 2021). Recent bulk-tissue transcriptomic analysis revealed that prenatal heat call exposure induces anticipatory gene expression changes in the embryonic hypothalamus, with robust enrichment in neurovascular and glial cell populations, suggesting adaptive reorganization in preparation for thermal stress (Subba et al., 2026). The hypothalamus integrates such environmental signals through neuroendocrine circuits, particularly the paraventricular nucleus (PVN), which coordinates metabolic and stress-adaptive responses to thermal challenge (Herman and Tasker, 2016). However, the cellular heterogeneity of the hypothalamus precludes identification of which specific cell types (e.g., glial, neuronal, or neuroendocrine) are the primary targets of acoustic developmental programming. Moreover, the regulatory mechanisms through which early sensory experience establishes lasting developmental changes in hypothalamic circuits remain unknown.

While neurons have traditionally dominated models of developmental programming, astrocytes – glial cells that constitute nearly half of all brain cells – are emerging as critical mediators of adaptive responses to environmental challenges (Bhatia et al., 2019; Bylicky et al., 2018; Perea et al., 2014). In the developing hypothalamus, astrocytes provide structural scaffolding, metabolic and nutritive support, and neuroprotective factors that buffer neural circuits against stress (Beard et al., 2021; Bélanger and Magistretti, 2009; Sloan and Barres, 2014; Verkhratsky and Nedergaard, 2018). The timing of astrocyte differentiation is tightly controlled by coordinated transcriptional programs: Nuclear Factor I (NFI) transcription factors drive astrocyte specification (Deneen et al., 2006; Wilczynska et al., 2009), while Notch signaling orchestrates the developmental transition from neurogenesis to gliogenesis (Taylor et al., 2007; Zhou et al., 2010), and the proneural transcription factor ASCL1 (Achaete-scute homolog 1) controls expression of Delta-like Notch ligands to balance neuronal versus glial fate decisions in a context-dependent manner (Nelson et al., 2009; Tran et al., 2023; Vue et al., 2014). These developmental transitions are further modulated by local thyroid hormone availability, with astrocytic deiodinases and thyroid hormone transport proteins regulating the pace of glial maturation and the timing of developmental plasticity windows (Ge et al., 2020). Critically, astrocyte maturation influences critical period plasticity through the production of extracellular matrix components and the regulation of synaptic metabolism (Ameroso and Rios, 2024; Bhatia et al., 2019; Brandt and Ackerman, 2025; Bylicky et al., 2018; Hashimoto et al., 2025; Lin et al., 2025; Ribot et al., 2021; Starkey et al., 2023). The astrocyte and neurovascular enrichment observed in heat call-responsive genes at bulk-tissue resolution (Subba et al., 2026) suggests that non-neuronal cells may represent primary targets of prenatal acoustic programming, yet whether, and through what regulatory mechanisms, early sensory experience selectively remodels non-neuronal developmental programs remains unexplored.

We hypothesized that prenatal acoustic experience triggers cell-type-selective molecular reorganization in the developing hypothalamus, targeting both glial populations (based on bulk-tissue transcriptomic enrichment) and neuroendocrine circuits (based on the role of the PVN in coordinating thermal adaptation). To identify the specific cell types and regulatory mechanisms underlying heat call-induced programming, we performed single-nucleus multiome sequencing, pairing single-nucleus ATAC-seq (snATAC-seq) and single-nucleus RNA-seq (snRNA-seq), on the medial hypothalamus of embryonic day 13 zebra finch embryos exposed to either a prenatal heat call or a control call playback. This approach enables simultaneous profiling of chromatin accessibility and gene expression at single-cell resolution, allowing us to address three key questions: (1) Which hypothalamic cell types exhibit chromatin and transcriptional responses to prenatal heat calls, and do responses differ by sex? (2) What transcriptional regulatory networks are activated by early acoustic experience? (3) Does prenatal acoustic signaling alter developmental trajectories of specific cell lineages? Our findings reveal that prenatal acoustic experience triggers astrocyte-selective chromatin reorganization at gliogenic regulatory elements and sex-dimorphic regulation of thyroid hormone signaling in neuroendocrine neurons, suggesting that early-life sensory programming reshapes glial and neuronal developmental programs in the hypothalamus, thereby preparing circuits for anticipated environmental challenges.

## 2 Results

### 2.1 Single-Nucleus Multiomic Profiles Revealed Diverse Clusters in the Hypothalamus of Zebra Finch Embryos

We used the 10X Chromium Single Cell Multiome ATAC + Gene Expression kit to profile the transcriptome and chromatin accessibility via snRNA-seq and snATAC-seq in 8 pooled samples of embryonic day 13 zebra finch embryos (4 heat call and 4 control call exposed) (Figure 1A). We processed medial hypothalamic punches from E13 zebra finch embryos to obtain joint profiles of chromatin accessibility and gene expression from 57,388 nuclei in the snRNA-seq analysis (Supplemental Table 1), and 70,720 nuclei in the snATAC-seq analysis (Supplemental Table 2). Each data set was analyzed independently to assess differences in clustering between RNA and ATAC modalities (Figure 1B). Overall, the snATAC-seq dataset (57 clusters) had more clusters than the snRNA-seq dataset (47 clusters). An unsupervised joint clustering using the Weighted Nearest Neighbor (WNN) method was performed on the paired modalities of the same single nuclei. The WNN method analyzed 55,950 nuclei across both modalities and resulted in 49 clusters (Figure 1C, Supplemental Table 3). The identified cell clusters included 27 neuronal clusters, 14 glial clusters, and 8 non-neuronal clusters (endothelial, vascular, and epithelial cells) based on marker gene expression (Figure 1C). These WNN clusters were grouped into 13 cell types: glutamatergic neurons (18,530 nuclei), astrocytes (14,949 nuclei), GABAergic neurons (11,444 nuclei), glutamatergic-GABAergic neurons (4,850 nuclei), oligodendrocytes (2,459 nuclei), microglia (1,156 nuclei), ependymal cells (633 nuclei), endothelial cells (479 nuclei), fibroblasts (431 nuclei), macrophages (341 nuclei), capillary endothelial cells (328 nuclei), vascular smooth muscle cells (247 nuclei), and choroid plexus epithelial cells (103 nuclei) (Figure 1D, Supplemental Table 3).

**Figure 1:**
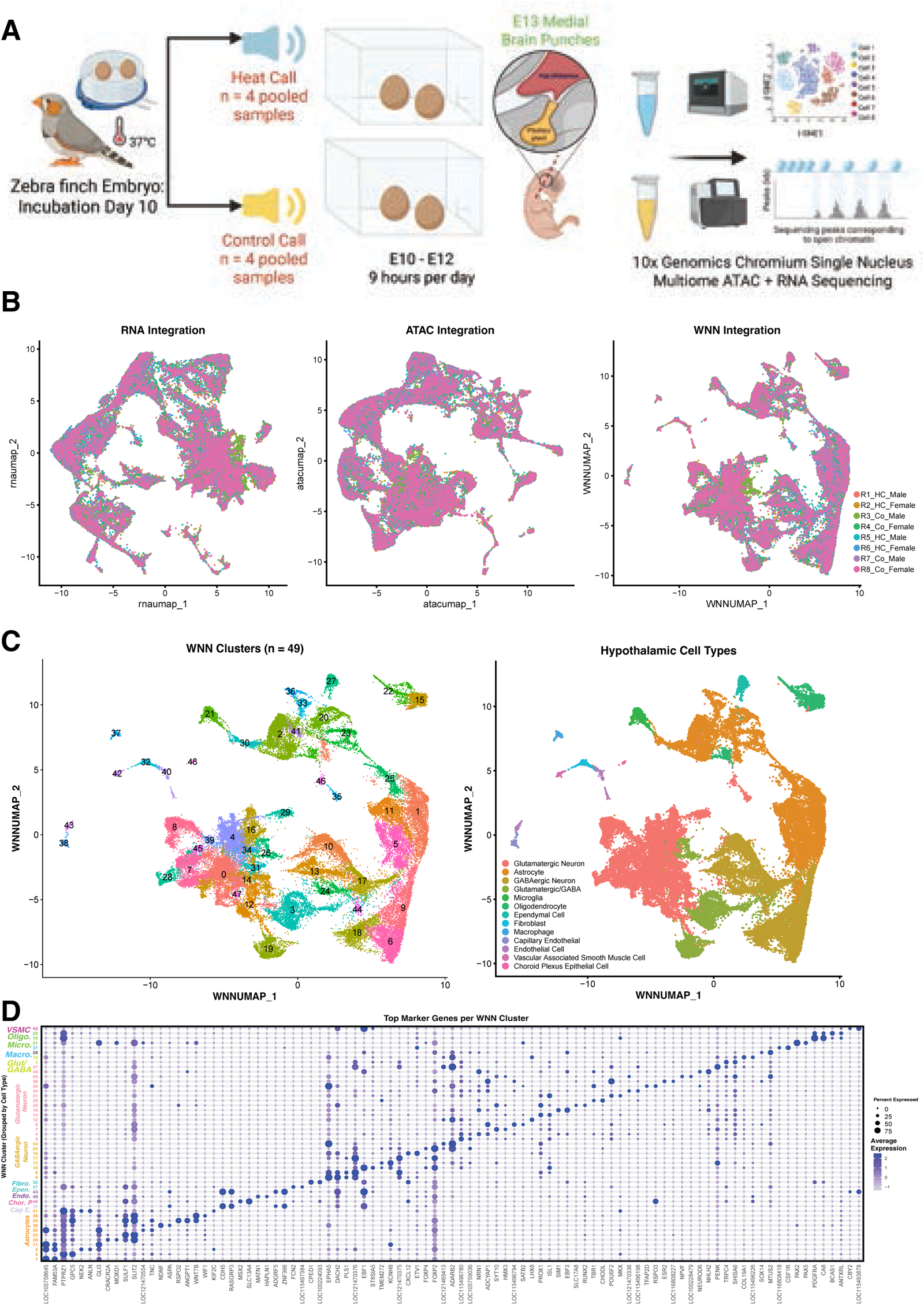
(A) Schematic of the experimental design for RNA-sequencing in late-stage zebra finch embryos exposed to chronic heat call (n = 4) or control call (n = 4) playback. (B) UMAP visualization of single-nucleus RNA-seq (left), ATAC-seq (middle), and weighted nearest neighbor (WNN) integration (right) colored by sample identity. RNA integration used SCT-normalized expression data (top 35 principal components). ATAC integration used latent semantic indexing (LSI, dimensions 2-50). WNN integration combined both modalities using weighted nearest neighbor analysis. n = 55,950 nuclei across 8 samples (4 control, 4 heat call treatment; 4 male, 4 female). (C) UMAP of WNN-integrated data colored by 49 unsupervised clusters identified at resolution 0.6 using Louvain clustering on the weighted shared nearest neighbor graph (left). UMAP of WNN-integrated data colored by 13 hypothalamic cell type annotations assigned based on marker gene expression and differential expression analysis (right). (D) Dot plot of the top two differentially expressed marker genes for each of 49 WNN clusters, ordered by cell type annotation. Marker genes were identified by differential expression testing (adjusted P < 0.05, pct.1 ≥ 0.25, pct.1 − pct.2 ≥ 0.25) and ranked by log2 fold-change. Dot size indicates percentage of expressing cells; color shows scaled average expression (gray to blue). Heat Call (HC); R (Replicate); Control (Co). Panel A was created using BioRender.com

### 2.2 Heat Call Exposure is Associated with Transthyretin Gene Expression Changes in a Sexual Dimorphic Manner

To profile gene expression differences between heat-call-exposed and control-call-exposed embryos, we used a sex-stratified and non-sex-stratified method. We noticed that transthyretin (*TTR*) was downregulated in heat-call-exposed embryos in a glutamatergic cell cluster (cluster #4) in the non-sex stratified analysis (Figure 2A). *TTR* had the highest expression in choroid plexus epithelial cells, for which it also served as a marker gene (cluster #48; Supplemental Table 4). Further, *TTR* was downregulated in multiple cell clusters (20 out of 49 WNN clusters) in female heat-call-exposed embryos when compared to female control-call-exposed embryos (Supplemental Table 5). Further, one specific glutamatergic cell cluster (cluster #16) with high expression of SIM BHLH Transcription Factor 1 (*SIM1*), a paraventricular nucleus marker, displayed sexually dimorphic expression of *TTR*, with upregulation in males and downregulation in females (Figure 2B). Taken together, the sexually dimorphic expression of *TTR* in the paraventricular nucleus and its downregulation in multiple cell clusters are associated with developmental programming in heat-call-exposed embryos.

**Figure 2:**
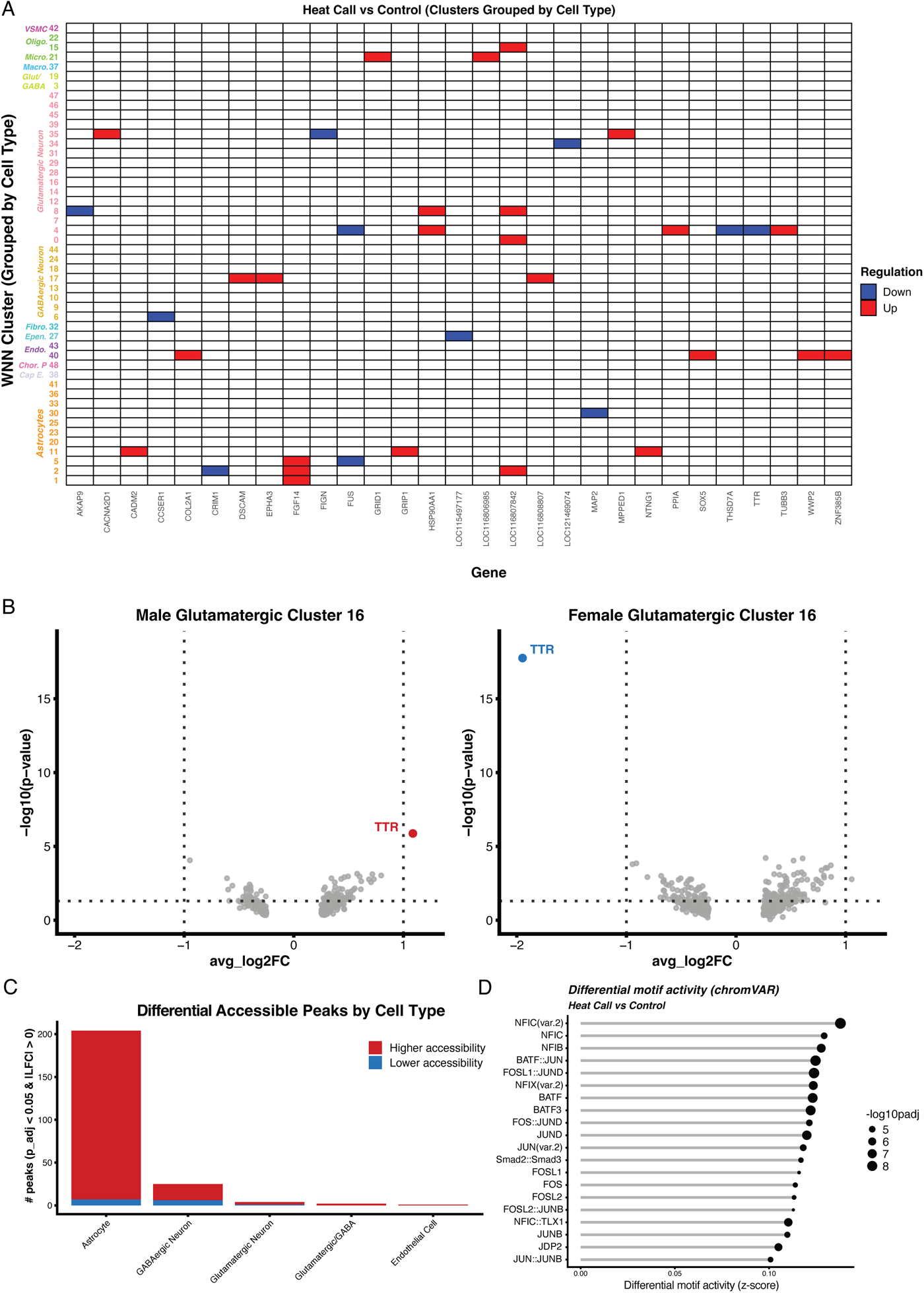
Heat call playback induces cell type-specific transcriptional and chromatin accessibility changes involved in stress response in hypothalamic cells. (A) Cell type-specific differential gene expression under heat call playback. Heatmap showing differentially expressed genes (DEGs) across 49 WNN clusters comparing heat call versus control conditions. Red: upregulated; blue: downregulated (adjusted P < 0.05). Clusters are grouped by cell type annotation. (B) Sexually dimorphic expression of *TTR* in glutamatergic neurons (cluster 16). Volcano plots for male (left, upregulated in red) and female (right, downregulated in blue) embryos in cluster 16 (glutamatergic neurons). X-axis: Average log2 fold-change; y-axis: −log10(P-value). Dotted lines indicate significance thresholds (|log2FC| > 1, P < 0.05). (C) Differential accessibility across hypothalamic cell types. Stacked bar plot showing the number of differentially accessible (DA) peaks per cell type (adjusted P < 0.05, |log2FC| > 0). Red: increased accessibility in heat call; blue: decreased accessibility. DA peaks were identified per WNN cluster and aggregated by hypothalamic cell type annotation. Astrocytes and GABAergic neurons show the most extensive chromatin remodeling. (D) Transcription factor motifs with elevated activity in heat call exposed embryos. Ranked lollipop plot showing the top 20 motifs with significantly increased chromVAR deviation scores in heat call versus control embryos. The x-axis represents the mean difference in per-cell chromVAR z-scores; point size indicates statistical significance (−log₁₀ adjusted P-value, Wilcoxon rank-sum test). Nuclear Factor I-C (*NFIC*) exhibits the highest differential activity (mean deviation = 0.138, adjusted P = 1.47 × 10⁻⁹), followed by four AP-1 family members (FOSL1::JUND, BATF::JUN, BATF3, BATF), indicating coordinated activation of astrocyte differentiation and stress-responsive transcriptional programs.

### 2.3 Developmental Programming is Linked to Higher Chromatin Accessibility in Astrocyte Clusters

To profile epigenetic changes associated with prenatal heat call playback, we performed differential chromatin accessibility (DA) analysis between heat-call and control-call-exposed embryos across all 49 WNN clusters. We identified 236 differentially accessible (DA) chromatin regions (adjusted P < 0.05, |log₂ Fold Change (FC)| > 0, MAST test) with a predominant increase in chromatin accessibility (222 peaks vs. 14 peaks showing decreased accessibility) in heat-call-exposed embryos (Figure 2C, Supplemental Table 6). Differentially accessible peaks were annotated to capture both direct (overlapping) and nearest-gene (proximal or distal) relationships between chromatin accessibility sites and the gene. Strikingly, there was a cell-type-specific distribution of chromatin accessibility changes, with astrocytes showing the highest chromatin remodeling in 204 DA peaks (86.4% of all DA regions; Supplemental Table 6). Within astrocytes, 197 out of 204 DA peaks displayed an increase in chromatin accessibility in heat-call-exposed embryos (Supplemental Table 6). GABAergic neurons (25 DA peaks; 19 increased, 6 decreased) followed by glutamatergic neurons (4 DA peaks) showed fewer changes in chromatin accessibility (Supplemental Table 6). Non-neuronal cell types including microglia, endothelial cells, and vascular cells showed minimal chromatin accessibility changes (0-1 DA peaks each; Figure 2C, Supplemental Table 6). These results show an enrichment of astrocyte-specific chromatin changes and position astrocytes as key mediators of developmental programming by prenatal heat call playback in the embryonic hypothalamus.

### 2.4 Altered Activity of Nuclear Factor One Transcription Factors in Heat Call Exposed Embryos

To identify transcription factors (TFs) driving chromatin remodeling in heat-call-exposed embryos, we quantified per-nucleus TF activity using the chromVAR algorithm (reference), which computes deviation scores reflecting accessibility of peaks containing TF motifs after correcting for GC content and technical biases. We scanned all accessible chromatin regions against the JASPAR 2020 vertebrate motif database (reference) and identified 27 TF motifs with significantly elevated activity in heat-call-exposed versus control-call-exposed embryos (adjusted P < 0.05, Wilcoxon rank-sum test; Figure 2D, Supplemental Table 7). Nuclear Factor I-C (NFIC) emerged as the single most enriched transcription factor motif, showing the strongest differential activity (mean chromVAR deviation = 0.138, adjusted P = 1.47 × 10⁻⁹; Figure 2D, Supplemental Table 7). The NFI family is particularly notable given its established role in astrocyte differentiation and chromatin remodeling (Cobo et al., 2023; Singh et al., 2011). The four next highest-ranked motifs belonged to the AP-1 superfamily of transcription factors involved in cellular stress response: FOSL1::JUND (mean deviation = 0.124, adjusted P = 3.92 × 10⁻⁹), BATF::JUN (mean deviation = 0.125, adjusted P = 4.39 × 10⁻⁹), BATF3 (mean deviation = 0.122, adjusted P = 1.51 × 10⁻⁸), and BATF (mean deviation = 0.123, adjusted P = 1.92 × 10⁻⁸; Figure 2D, Supplemental Table 7) (Alaiz-Noya et al., 2025). The elevated NFIC activity we observed directly links our chromVAR findings to the astrocyte-dominant chromatin accessibility changes described above. Together, the co-enrichment of NFIC and multiple AP-1 family members among the top enriched motifs suggest that prenatal heat call exposure activates coordinated regulatory programs linking glial maturation and stress-adaptive transcriptional networks.

### 2.5 Transcriptional Co-Expression Network Analysis Identifies Heat Call-Responsive Astrocyte Module Astro-M7

To identify coordinated changes in gene expression associated with heat call playback in astrocytes, we conducted a weighted gene co-expression network analysis using 10,901 genes expressed in ≥ 5% of nuclei. We identified 12 co-expression astrocytic modules representing distinct transcriptional programs (Figure 3A). Module Astro-M7 exhibited the strongest positive association with heat call exposure (r = 0.042, adjusted P = 1.49 × 10⁻²²; Fig. 3A), indicating coordinated upregulation of this 50-gene module in heat-call-exposed embryos (Supplemental Table 9). Astro-M7 module activity, quantified by module eigengene scores, showed an astrocyte subtype-specific response with higher expression in subcluster 0, *FOXG1*/*KCNQ5*-positive progenitor-like astrocyte-associated cells, and in subcluster 2 (*AURKA*-positive proliferative astrocyte-associated precursor-like cells) (mean scores: 4.37 and 4.02, respectively), while subcluster 1 (*THRB1*-positive developing astrocyte-associated cells) and subcluster 3 (*SHH*/*NKX2-4*/*SLIT2*-positive ventral gliogenic-like astrocyte-associated cells) showed negative module scores (mean: -1.11 and -0.72; Figure 3C). Finally, the Astro-M7 module consisted of genes enriched in GO terms associated with positive regulation of Notch receptor target (Fold Enrichment = 146.66 (zebra finch annotation), FDR < 0.05; Figure 3E, Supplemental Tables 10-11). These results suggest that heat call-associated transcriptional programs are enriched in specific astrocyte-associated subpopulations, rather than uniformly distributed across all astrocytes.

**Figure 3:**
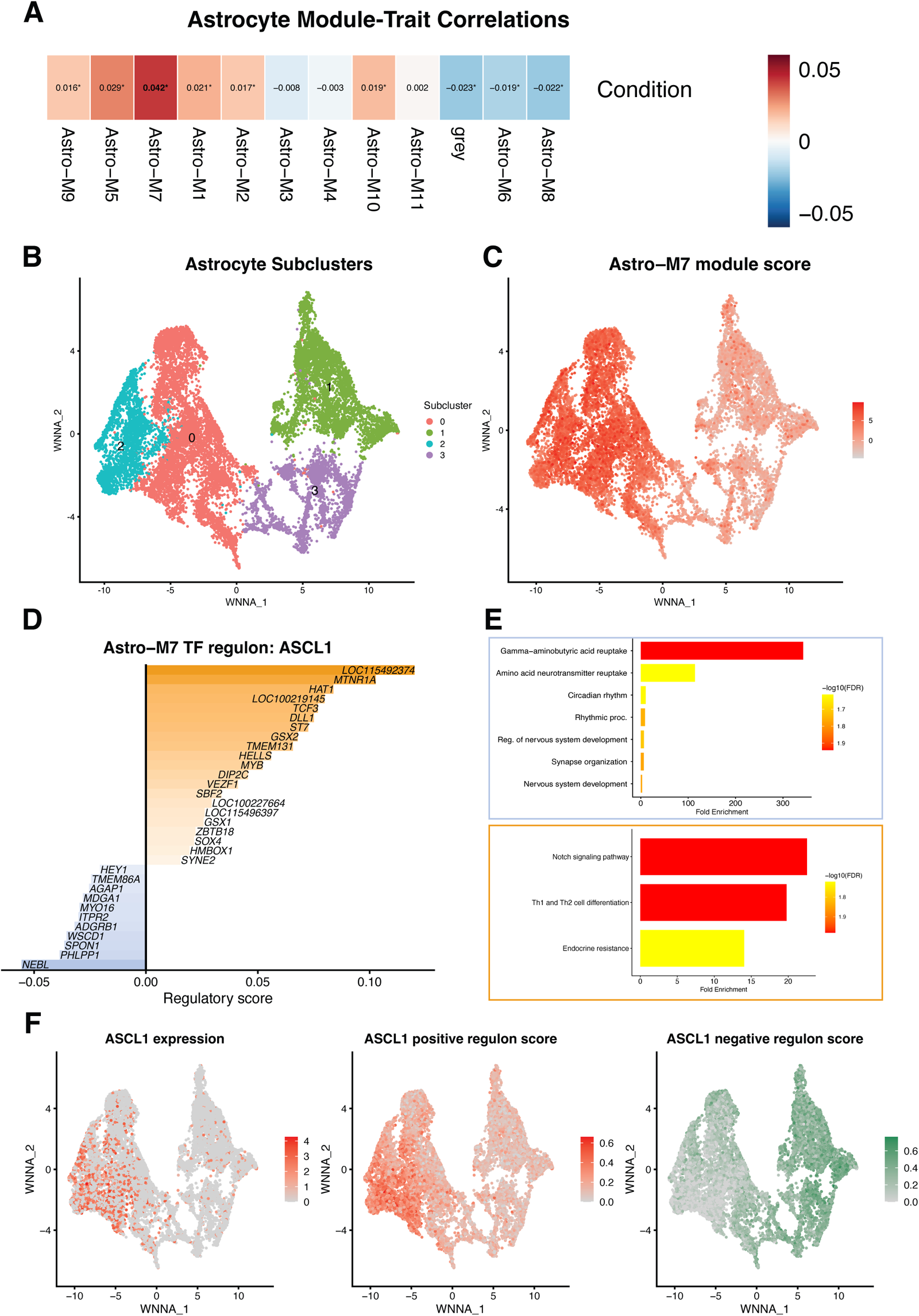
Heat call-responsive astrocyte module Astro-M7 is regulated by transcription factor ASCL1. (A) Module-trait correlation heatmap showing Pearson correlation coefficients between astrocyte co-expression module eigengenes (n = 12 modules) and heat call playback exposure (binary treatment: Control = 0, Heat = 1; n = 14,949 nuclei). Astro-M7 exhibits the strongest positive correlation with heat exposure (r = 0.042, FDR = 1.49 × 10⁻²²). Color intensity indicates correlation strength; asterisk denotes statistical significance (Student’s asymptotic test with Benjamini-Hochberg FDR correction). (B) WNN UMAP embedding of zebra finch embryonic astrocytes (n = 14,949 nuclei) reveals four transcriptionally distinct astrocyte-associated subpopulations identified by Leiden clustering (resolution 0.06, algorithm 3). Each point represents a single nucleus colored by subcluster identity. (C) Astro-M7 module eigengene scores across subclusters. Color intensity (grey to red) indicates module activity, with elevated scores in subclusters 0 (*FOXG1/KCNQ5*-positive progenitor-like astrocyte-associated cells) and 2 (*AURKA*-positive proliferative astrocyte-associated precursor-like cells). (D) Transcription factor regulatory network for *ASCL1* within the Astro-M7 module. Bar plot displays XGBoost-derived regulatory scores for *ASCL1* target genes (regulatory scores ≥ 0.01). (E) GO enrichment analysis (KEGG and GO Biological Process) of *ASCL1* targets. Upper: *ASCL1* negative regulon (repressed targets, correlation ≤ -0.05) enriched for GABA metabolism and circadian processes. Lower: *ASCL1* positive regulon (activated targets) enriched for Notch signaling and Th1/Th2 cell differentiation. Color = -log₁₀(FDR); bar length = fold enrichment. All FDR < 0.05. (F) *ASCL1* regulatory activity across subclusters. Left: *ASCL1* gene expression (log-normalized counts) shows elevated expression in subclusters 0 and 2. Middle: *ASCL1* positive regulon score (aggregated expression of positively correlated target genes) mirrors *ASCL1* expression pattern. Right: *ASCL1* negative regulon score (aggregated expression of negatively correlated target genes, correlation ≤ -0.05) shows complementary expression pattern, with higher activity in subclusters 1 (*THRB1*-positive developing astrocyte-associated cells) and 3 (*SHH/NKX2-4/SLIT2*-positive ventral gliogenic-like astrocyte-associated cells).

### 2.6 Transcription Factor ASCL1 Orchestrates Subtype-Specific Regulation in Astrocytes

To identify TFs involved in regulating Astro-M7 transcriptional programs in heat call-associated developmental programming, we performed TF regulatory network analysis using a motif-informed, XGBoost-based framework (reference). We inferred TF-target relationships by assigning regulons using a regulatory gain threshold of 0.01. We identified *ASCL1* (*Achaete-scute homolog 1*) as a central regulatory transcription factor with 32 target genes (gain ≥ 0.01) with top targets such as *MTNR1A* (*melatonin receptor 1A*; gain = 0.103), *HAT1* (*histone acetyltransferase 1*; gain = 0.084), *TCF3* (transcription factor 3; gain = 0.077), *DLL1* (*delta-like 1*; gain = 0.075), and *GSX2* (*insert full gene name here*, gain = 0.068), all showing strong positive correlations with *ASCL1* expression (correlation coefficients > 0.50; Figure 3D). To characterize the functional consequences of *ASCL1* regulatory activity, we performed gene ontology enrichment analysis on *ASCL1* regulon targets identified using a relaxed regulatory gain threshold (≥ 0.005), separately analyzing positive regulon targets (positively correlated with *ASCL1*) and negative regulon targets (negatively correlated with *ASCL1*). *ASCL1* negative regulon targets were enriched for gamma-aminobutyric acid (GABA) metabolism (Fold Enrichment = 342.7, FDR < 0.05) and circadian rhythm processes (Figure 3E). *ASCL1* positive regulon targets were enriched for Notch signaling pathway (Fold Enrichment = 22.54, FDR < 0.05) and Th1/Th2 cell differentiation processes (Fold Enrichment = 19.78, FDR < 0.05; Figure 3E). Finally, *ASCL1* had a subcluster-specific regulatory response with elevated gene expression and positive regulon scores (aggregated expression of activated targets) in subcluster 0 (*FOXG1*/*KCNQ5*-positive progenitor-like astrocyte-associated cells), and in subcluster 2 (*AURKA*-positive proliferative astrocyte-associated precursor-like cells), while *ASCL1* negative regulon scores (aggregated expression of repressed targets) showed complementary enrichment in subcluster 1 (*THRB1*-positive developing astrocyte-associated cells) and subcluster 3 (*SHH*/*NKX2-4*/*SLIT2*-positive ventral gliogenic-like astrocyte-associated cells) (Figure 3F). Together, these results suggest that prenatal heat call playback may influence *ASCL1* regulatory activity in a tightly controlled way across astrocyte-associated subpopulations by suppressing GABAergic differentiation programs in developing astrocyte-associated cells and by shifting the balance of Notch-*ASCL1* interactions, consistent with altered astrocyte developmental-state regulation across subpopulations.

### 2.7 Heat Call Playback Drives Extensive Chromatin Reorganization in Astrocytes

To investigate gene regulatory programs associated with heat call playback, we performed chromatin co-accessibility analysis using Cicero (Pliner et al., 2018) on astrocyte snATAC-seq data. We identified 3,903,816 genome-wide chromatin connections in astrocytes using optimized parameters (250 kb distance constraint, 500 kb window, smoothing parameter s = 0.75). We defined condition-specific “chromatin rewiring” as connections active in one playback condition (both endpoint peaks accessible in ≥ 5% of astrocytes, Cicero co-accessibility ≥ 0.15) but inactive in the other (at least one endpoint accessible in < 5% of astrocytes) (Figure 4A). Heat call playback induced high chromatin re-organization, establishing 9,502 heat call-specific loops involving 3,891 unique regulatory elements, compared to only 248 control-specific loops (Figure 4B). Next, we identified heat call-specific *cis*-regulatory loops to gene promoters (± 2 kb from Transcription Start Site (TSS)) and stratified 198 genes by rewiring burden: high (≥ 10 promoter-anchored connections; n = 19), moderate (5-9 connections; n = 18), and low (1-4 connections; n = 161). RNA log₂ fold-changes (Heat Call vs. Control) showed no relationship with rewiring extent (Pearson r = −0.002, p = 0.976; Supplemental Figure 2). These results suggest that heat call playback induces regulatory chromatin rewiring, which is decoupled from immediate transcriptional changes.

**Figure 4:**
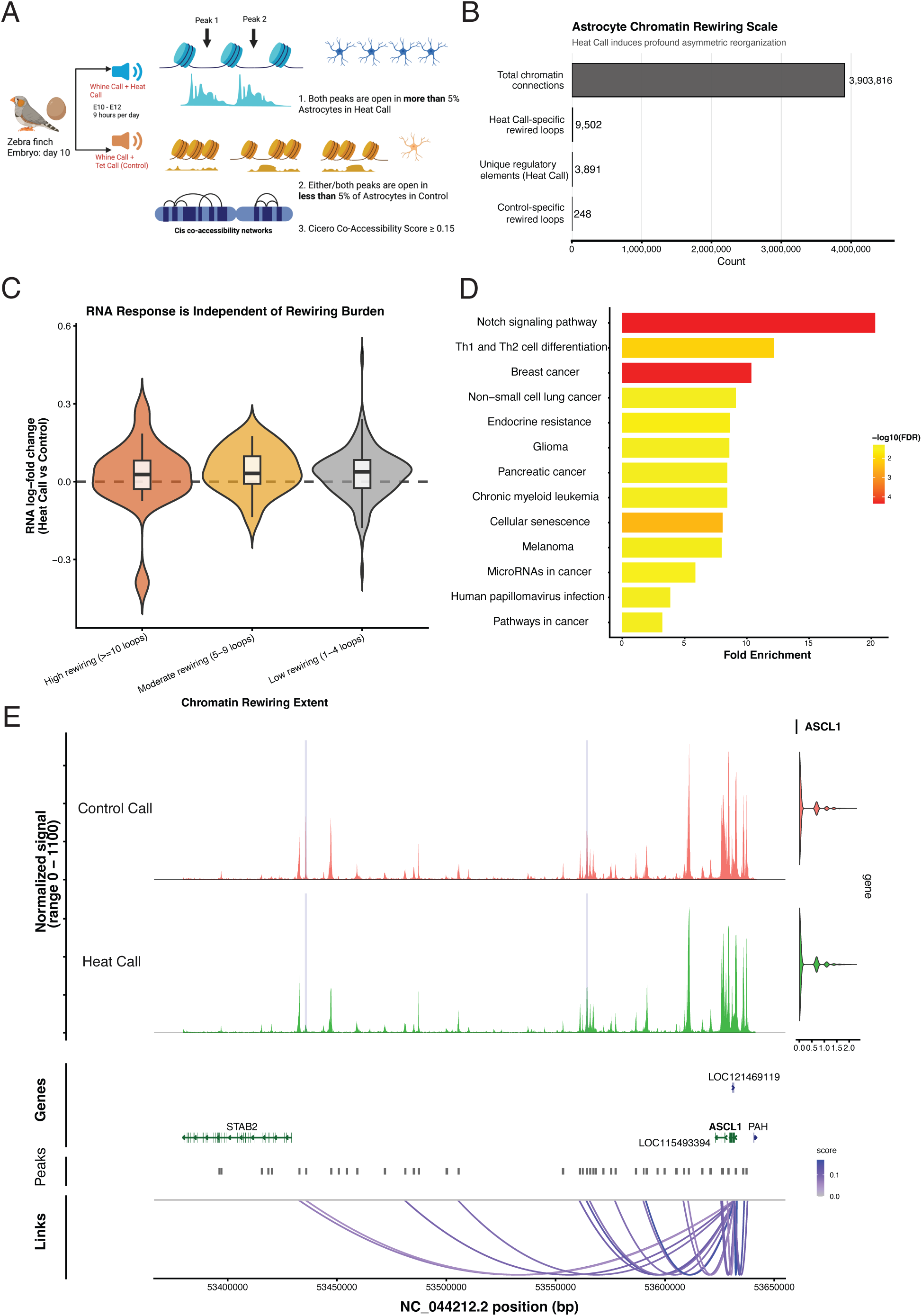
Heat call playback drives chromatin reorganization in astrocytes. Chromatin co-accessibility analysis identifies playback-specific rewiring in zebra finch embryonic astrocytes. (A) Cicero analysis workflow defines rewired connections as active in heat call playback (both peaks accessible in ≥ 5% of astrocytes, co-accessibility ≥ 0.15) but inactive in control (at least one peak accessible in < 5% of astrocytes). (B) Scale of astrocyte chromatin reorganization in response to heat call playback. Bars report the total number of chromatin connections assessed (3,903,816), heat call-specific rewired loops (9,502), the number of unique regulatory elements participating in heat call rewiring (3,891), and control-specific rewired loops (248). (C) Relationship between chromatin rewiring extent and transcriptional response. Genes are grouped by the number of heat call-specific promoter-anchored links (high: ≥ 10; moderate: 5-9; low: 1-4), and violin plots show RNA log₂ fold-change (Heat Call vs Control) for each group with embedded boxplots (median, interquartile range, 1.5×IQR whiskers). No group-level difference in RNA response is detected (Kruskal-Wallis p = 0.763). (D) GO enrichment reveals Notch signaling and developmental pathways in genes with rewired promoters (FDR < 0.05). (E) Chromatin co-accessibility at the *ASCL1* locus in astrocytes from heat call (n = 4) versus control (n = 4) E13 zebra finch embryos, where n represents pooled biological replicates (5 medial hypothalamic punches per pool). Top: ATAC-seq coverage tracks (normalized signal 0-1100) for control (red) and heat call (blue). Middle: Gene annotations on chromosome NC_044212.2. Bottom: Differentially accessible peaks and heat call-specific co-accessibility loops (purple arcs, Cicero score 0.0-0.1). Two flanking peaks show increased accessibility in heat call astrocytes and participate in enhancer-promoter loops (dual-hit peaks). Peak-to-gene correlation confirms association with *ASCL1* expression (Pearson r=0.138, p=3.6×10⁻^9^). Differential accessibility by MAST test (two-tailed, FDR < 0.05, log₂FC > 0.25). Co-accessibility parameters: 250 kb distance, 500 kb window, score ≥ 0.15.

Genes with high *cis*-regulatory chromatin rewiring in the promoter region included *LOC100226434* (*HES-5-like*; 52 loops), *NFIA* (45 loops), *ZFP36L2* (22 loops), and *DLL1* (18 loops) (Supplemental Figure 2). *NOTCH1*, a receptor of the Notch signaling pathway, included 6 *cis*-regulatory loops and was also significantly upregulated in heat call playback (RNA log_2_FC = 0.18, Bonferroni-adjusted P = 0.016, MAST test; Supplemental Figures 2-3). *DLL1*, a Notch ligand, was among the few rewired genes showing significant differential expression, consistent with activation of Notch-mediated developmental signaling cascades (Supplemental Figure 2). Gene ontology enrichment analysis of all genes with moderate and high promoter rewiring revealed significant enrichment for Notch signaling pathway components (Fold Enrichment = 20.3; FDR < 0.05; Figure 4D), as well as pathways involved in cell differentiation and developmental processes. Together, these results suggest that heat call playback is associated with *cis*-regulatory changes in Notch pathway loci consistent with altered astrocyte developmental programs.

To identify regulatory elements undergoing coordinated chromatin state changes, we identified the intersection of heat-call-specific rewired connections with DA peaks. Differential accessibility analysis identified 226 heat call-gained peaks (FDR < 0.05, |log₂FC| > 0.25) in astrocytes (Supplemental Table 12). A total of 57 dual-hit peaks (25% of DA-gained peaks) displayed both differentially accessible and co-accessible *cis*-regulatory loops (Supplemental Table 13). We performed peak-to-gene linking analysis to correlate peak accessibility at these dual-hit peaks with RNA expression of genes within 500 kb. This identified 10 significant peak-gene correlations (Pearson |*r*| > 0.1, p < 0.05) involving 9 unique genes, confirming that chromatin rewiring at these peaks associates with transcriptional regulation of nearby genes (Supplemental Table 14). Genes linked to these peaks included the proneural transcription factor *ASCL1* (*r* = 0.138, p = 3.6×10⁻^9^), the axon guidance receptor *PLXNA4* (*r* = 0.273, p = 9.1×10⁻¹¹), and the fibroblast growth factor receptor-like 1 *FGFRL1* (*r* = 0.130, p = 7.3×10⁻⁴), as well as genes involved in chromatin organization (*PRDM16*), neural differentiation (*KCNQ5*), and cell cycle regulation (*TACC3*; Supplemental Table 14). Two peaks flanking the *ASCL1* promoter exhibited increased accessibility specifically in heat-call-exposed astrocytes and were identified as dual-hit elements participating in heat call-specific enhancer-promoter loops (Figure 4E, Supplemental Table 14). Taken together, these chromatin connections were specific to heat call playback and were associated with coordinated opening of distal regulatory elements and establishment of long-range promoter-enhancer loops at neuronal differentiation genes like *ASCL1*.

### 2.8 Pseudotemporal Trajectory Analysis Identifies Condition-Associated Developmental Trajectory Shifts

To determine whether prenatal heat call playback was associated with altered developmental trajectory structure across hypothalamic cell types (26,758 control cells; 29,192 heat call cells), we performed pseudotemporal trajectory inference using Slingshot (reference) on weighted nearest neighbor (WNN) UMAP coordinates. Slingshot trajectories were inferred separately for control and heat call cells on the same WNN embedding, using cluster 0 as the root state (Figure 5A). Both conditions showed broadly similar trajectory structure across shared WNN clusters, consistent with conserved overall lineage organization. However, pseudotime mapping revealed condition-associated shifts in cell positions within multiple clusters, indicating differences in pseudotime distribution rather than overt trajectory divergence (Figure 5A).

**Figure 5:**
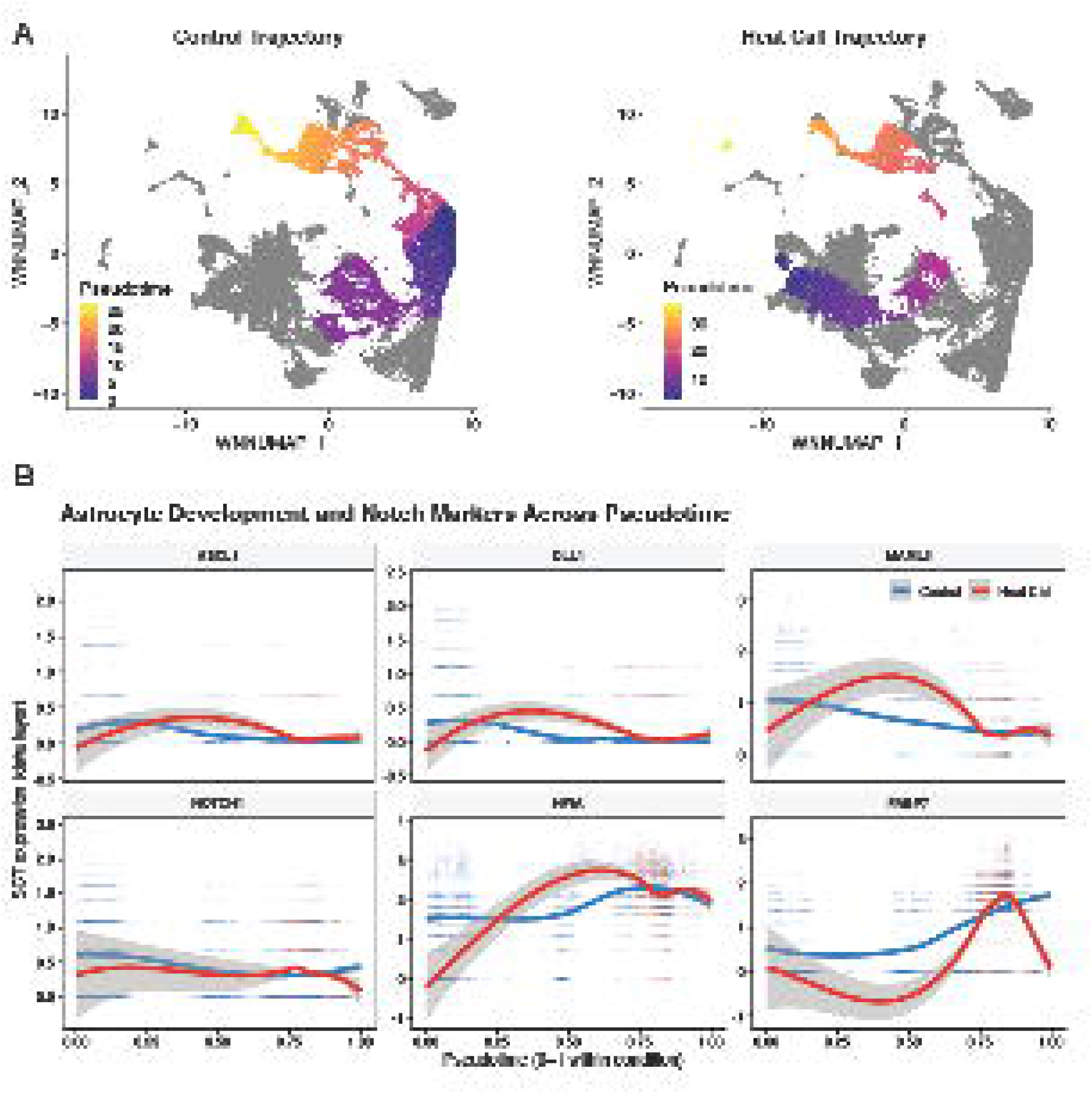
Pseudotemporal trajectory analysis identifies condition-associated shifts in hypothalamic developmental trajectories and astrocyte gene-expression dynamics. (A) Slingshot trajectories were inferred separately for control and heat call conditions on the full hypothalamic WNN-UMAP embedding containing all cell types, using WNN cluster identities (resolution 0.6) as input and cluster 0 as the root state. Cells are colored by principal-lineage Slingshot pseudotime values within each condition. (B) Astrocyte-specific expression dynamics of proneural transcription factor *ASCL1*, radial glial marker *FABP7*, and Notch pathway and astrocyte-development-state-associated genes (*DLL1, NOTCH1, MAML3, NFIA*) across rescaled pseudotime (0-1 within each condition). Points represent per-cell SCT-normalized expression values from astrocytes only; LOESS smoothers (span = 0.75) with 95% confidence intervals are overlaid for control (blue) and heat call (red) conditions.

To identify non-linear expression dynamics along developmental trajectories, we applied tradeSeq generalized additive modeling (GAM) (reference) with 6 knots per playback condition. We performed association testing and identified 5,594 pseudotime-dynamic genes in the control samples and 5,494 in the heat call samples, whose expression is significantly associated with developmental progression (Supplemental File 1). Of these, 3,999 genes were shared between both conditions, while 1,595 and 1,495 genes showed condition-specific dynamics in the control and heat call samples, respectively. The comparable numbers of condition-specific dynamic genes indicate that both conditions retain extensive pseudotime-dependent gene-expression programs, while differing in the specific sets of genes that vary along their respective trajectories.

To assess whether condition-associated pseudotime shifts were especially evident in astrocyte populations, we conducted a per-cluster pseudotime distribution analysis across all 42 WNN clusters and identified significant developmental shifts in 20 clusters (FDR < 0.05, two-sided Wilcoxon rank-sum test; Supplemental Table 15). The largest positive mean pseudotime shift in heat call cells was observed in cluster 1, an astrocyte-annotated WNN cluster (Δpt = +18.4, FDR = 4.7 × 10⁻³²; Supplemental Table 15). Because trajectories were inferred from gene expression separately by condition, we visualized expression of selected developmental-state genes associated with Notch signaling and astrocyte development along normalized pseudotime (0-1 scale) in the astrocyte subpopulation (Figure 5B). The proneural transcription factor *ASCL1* exhibited an early to mid-trajectory peak in both conditions (peak pseudotime 0.3-0.5), with a later pseudotime peak in heat call playback (Figure 5B). *ASCL1* and *DLL1* (ligand of the Notch pathway) exhibited broadly similar mid-pseudotime dynamics across conditions, with only modest separation between control and heat call trajectories (Figure 5B). *NOTCH1* did not show consistent late-stage elevation in heat call astrocytes, with only a small, localized bump in heat call near late pseudotime (Figure 5B). In contrast, *MAML3* displayed a pronounced mid-pseudotime increase in heat call astrocytes compared with the control condition, consistent with a condition-specific shift in Notch co-activator dynamics along the astrocyte developmental trajectory (Figure 5B). *NFIA*, an astrocyte-associated differentiation marker, increased across pseudotime in both conditions but remained higher in heat call across much of the trajectory, consistent with altered astrocyte trajectory-linked transcriptional dynamics in heat call cells (Figure 5B). *FABP7*, a radial glial cell marker, showed divergent late-stage behavior: heat call astrocytes exhibited a sharp late pseudotime spike followed by a decline toward pseudotime ∼1.0, whereas control astrocytes increased more gradually and remained elevated at the endpoint (Figure 5B). Together, these results indicate condition-associated differences in astrocyte trajectory-linked expression dynamics, including genes related to Notch signaling and astrocyte-associated developmental programs, in heat-call-exposed embryos.

## 3 Discussion

The mechanisms by which prenatal sensory experience establishes anticipatory developmental programs remain poorly understood, particularly at the cell-type-specific regulatory level. In zebra finches, parental heat call vocalizations program offspring for thermal environments (Mariette and Buchanan, 2016), inducing transcriptional changes enriched in hypothalamic non-neuronal cell populations (Subba et al., 2026). However, the cellular heterogeneity of the developing hypothalamus has obscured which specific cell types are primary targets of acoustic programming, and the regulatory mechanisms by which early experience drives lasting phenotypic change. Here, using single-nucleus multiome sequencing, we demonstrate that prenatal heat calls trigger astrocyte-selective chromatin reorganization in the embryonic hypothalamus. We found that extensive enhancer-promoter rewiring of developmental regulators precedes large-scale transcriptional output, revealing an epigenetic priming mechanism whereby acoustic experience poises glial differentiation programs for anticipated thermal stress. Our findings support a model in which prenatal acoustic cues epigenetically prime astrocyte regulatory programs in the hypothalamus, potentially preconditioning metabolic and homeostatic machinery for enhanced thermal resilience in post-hatch environments. This astrocyte-centric response challenges neuron-focused models of developmental programming and raises three key questions. Why are astrocytes preferential targets? What regulatory networks coordinate the astrocyte response? What are the functional consequences of altered astrocyte developmental-state regulation? We address each question in turn, beginning with the biological rationale for astrocyte-selective programming.

Our study shows a striking enrichment in chromatin regulatory rewiring occurs in astrocytes of heat-call-exposed embryos. This result challenges a neuron-centric developmental programming model with a cell-type specificity for astrocytes (204/236 DA peaks in astrocytes vs. 4 peaks in glutamatergic, and 25 peaks in GABAergic neurons) as key mediators of anticipatory adaptation (Figure 2C, Supplemental Table 6). Astrocytes are uniquely positioned as homeostatic integrators with processes ensheathing synapses and brain capillaries, allowing them to couple neural activity with metabolic supply and vascular dynamics (Bélanger and Magistretti, 2009; Verkhratsky and Nedergaard, 2018). This architectural positioning enables astrocytes to regulate energy metabolism through the astrocyte-neuron lactate shuttle (ANLS), in which astrocytes take up glucose via glucose transporter 1 (*GLUT1*), undergo glycolysis, and release lactate that supports neuronal energy demand and provides stress-buffering signals under oxidative challenge (Beard et al., 2021). Beyond metabolic support, astrocytes maintain extracellular K⁺ homeostasis through spatial buffering and control glutamate clearance via high-affinity transporters (EAAT1/GLAST), preventing hyperexcitability and excitotoxicity during periods of elevated neuronal activity (Ambroziak et al., 2025; Verkhratsky and Nedergaard, 2018). Astrocytes also regulate critical period plasticity through release of extracellular matrix components and regulation of synaptic pruning (Brandt and Ackerman, 2025; Ribot et al., 2021; Starkey et al., 2023), and recent work demonstrates their role in stabilizing mature neural circuits (Sancho et al., 2026). Taken together, these functional roles position astrocytes as ideal candidates for integrating prenatal acoustic cues such as heat calls and translating them into lasting metabolic and developmental reprogramming that enhances future thermal resilience.

The multiomic changes we observed in astrocytes likely reflect the developmental state of the embryonic hypothalamus during the E9-E13 exposure window. Our playback window spans from E9 to E13, representing the second half of the 14-day zebra finch incubation period. By E13, one day before hatching, the single-nucleus profiling identified 14,949 astrocyte nuclei (∼27% of all cell types), suggesting that astrocytes are abundant and transcriptionally active in the hypothalamus of the late-stage zebra finch embryo. Although direct markers of astrocyte developmental timing in zebra finch embryos have not yet been established, comparative avian systems such as the chicken embryo offer instructive parallels. In the chicken optic tectum, astroglial markers including *GFAP* and glutamine synthetase identify astroglial cells as early as embryonic day 9, with continued maturation toward hatching (Linser and Perkins, 1987). While extrapolation across species requires caution, given potential differences in developmental timing between precocial chicken and altricial zebra finch lineages, the timing of our E9-E13 playback window (mid to late-stage 14-day incubation period) aligns with the predicted neurogenesis-to-gliogenesis transition. This inference is supported by our own results with the abundance of astrocytes at E13, combined with extensive chromatin remodeling in astrocyte populations (204 of 236 differentially accessible peaks), indicating that we captured hypothalamic astrocytes during an active fate-commitment window. In the developing vertebrate brain, radial glial progenitors are multipotent neural stem cells that sequentially produce neurons during early development, then transition to producing glial cells during mid-to-late embryogenesis in a process called the “gliogenic switch” (Deneen et al., 2006; Molofsky and Deneen, 2015). This fate transition requires extensive chromatin remodeling in which Polycomb repressive complexes are reconfigured to progressively silence neurogenic programs while promoting astroglial fate specification (Hirabayashi et al., 2009; Pereira et al., 2010). This chromatin rewiring during the gliogenic switch could render progenitor cells particularly responsive to environmental inputs during this plastic chromatin state, which may stabilize as the cell commits to a mature astrocytic fate. This predicts that prenatal heat call exposure acts by engaging master regulators of the gliogenic switch - transcription factors and signaling pathways that coordinate astrocyte specification and developmental-state programs.

The molecular profile of heat-call-exposed astrocytes reveals a coordinated regulatory program consistent with engagement of the gliogenic switch. First, the convergent enrichment of NFIC motifs suggests that heat call exposure specifically engages a master regulator of astrocyte fate commitment, one that drives gliogenesis by opening gliogenic enhancers and linking the abrogation of neurogenesis to glial-fate specification (Deneen et al., 2006; Glasgow et al., 2017). Second, the reorganization of chromatin architecture around Notch signaling components points to a coordinated rewiring of the neurogenic-to-gliogenic transition machinery. The Notch pathway promotes the maintenance of radial glial progenitors and regulates the timing of this transition (Gaiano et al., 2000; Ge et al., 2002). Notch pathway genes, including *NOTCH1* and *DLL1*, exhibited both differential accessibility and condition-specific enhancer-promoter rewiring in heat call-exposed astrocytes (Supplemental Figures 2-3). Third, the positioning of *ASCL1* as a hub linking positive regulation of Notch signaling components to suppression of GABAergic neuronal differentiation programs suggests it may serve as a molecular switch that biases progenitor transcriptional output away from neuronal and toward glial fates. Together, these chromatin accessibility shifts, chromatin rewiring events, and transcriptional network changes indicate a coordinated regulatory program rather than isolated epigenetic alterations. We interpret these findings as evidence that prenatal heat call exposure alters the regulation and timing of gliogenic programs in late-stage hypothalamic astrocyte-associated populations. Alternatively, heat call playback could bias lineage choice toward astrocytes at the expense of neurons or stabilize already-committed astrocyte identity. However, these alternative explanations are less consistent with our observation that the most extensive chromatin remodeling occurs in progenitor-associated regulatory elements (e.g., *NFIA* and *HES5-like* genes) rather than mature astrocyte markers. This model predicts that heat-call-exposed embryos may exhibit altered timing of astrocyte-state acquisition and associated developmental plasticity windows in hypothalamic circuits.

Consistent with the activation of gliogenic regulatory programs, pseudotemporal trajectory analysis revealed condition-associated shifts in pseudotime distributions and astrocyte gene-expression dynamics in heat call-exposed embryos. Heat-call-exposed astrocytes showed shifted pseudotime distributions within developmental trajectories, with the largest positive mean pseudotime shift observed in an astrocytic cluster (WNN cluster 1; Δpt = +18.4, FDR = 4.7 × 10⁻³²; Figure 5A, Supplemental Table 14). Transcription factors involved in astrocytic differentiation programs, such as *NFIA* (Nuclear Factor I A), a gliogenic switch, and *FABP7* (fatty acid binding protein-7), a radial glial marker, showed altered temporal dynamics along the astrocyte differentiation continuum (Figure 5B) (Tchieu et al., 2019; Young et al., 2013). Specifically, *NFIA* remained elevated in heat call astrocytes across much of the pseudotime trajectory, while *FABP7* exhibited a decline toward the trajectory endpoint in astrocytes from heat call playback. Further, components of the Notch signaling pathway and *ASCL1* were elevated in early to mid-pseudotime in heat call astrocytes compared to controls, consistent with condition-specific shifts in Notch signaling dynamics during astrocyte maturation. Notch signaling has also been shown to inhibit neuronal differentiation programs by suppressing *ASCL1* activity, suggesting that co-expression of both Notch pathway components and *ASCL1* could reprogram cell fate toward a gliogenic fate (Park et al., 2017). These trajectory-associated transcriptional differences are accompanied by extensive chromatin rewiring, with 9,502 heat-call-specific enhancer-promoter loops involving 3,891 unique regulatory elements compared to only 248 control-specific loops (Figure 4B). Genes with the highest *cis*-regulatory co-accessibility, including *LOC100226434* (*HES-5*-like, 52 loops), *NFIA* (45 loops), *ZFP36L2* (22 loops), and *DLL1* (18 loops), are established regulators of glial differentiation and Notch-mediated developmental signaling (Supplemental Figure 2). Peak-to-gene correlation analysis identified a significant association between differentially accessible dual-hit peaks (both gained accessibility and participating in heat call-specific co-accessibility loops) and gene expression at two *cis*-regulatory peaks in *ASCL1* (r = 0.138, p = 3.6×10^-9^; Figure 4E, Supplemental Table 13). Together, these pseudotemporal and chromatin architectural changes support a model in which prenatal heat call exposure alters astrocyte developmental-state regulation through coordinated establishment of long-range regulatory interactions at lineage-associated loci.

Taken together, our chromatin and trajectory analysis results support a capacity priming model in which prenatal heat calls reshape astrocyte regulatory potential and alter trajectory-linked developmental programs by establishing regulatory changes at lineage and signaling hubs (e.g., NFI/Notch-associated architecture) that are only partially translated into immediate steady-state RNA differences at E13 (Supplemental Figure 2). This interpretation is consistent with our observation that heat call-induced *cis*-regulatory rewiring is extensive and strongly astrocyte-biased, yet the magnitude of promoter rewiring is largely decoupled from concurrent gene expression changes. This suggests that acoustic experience may set gene regulatory potential before downstream effector programs fully activate. A plausible functional consequence is that thermoregulatory hypothalamic networks may enter post-hatch life with an altered glial support state, increasing their margin for maintaining circuit stability when neuronal activity and metabolic demand rise during heat exposure. This hypothesis is motivated by evidence that heat acclimation requires plastic changes in the preoptic area circuitry of the hypothalamus, including the emergence of intrinsically warm-sensitive, pacemaker-like firing in preoptic hypothalamic neurons that is necessary for heat tolerance (Ambroziak et al., 2025). Under this framework, the key prediction is not that heat calls must induce maximal expression of heat-adapted effectors at E13, but rather that earlier developmental time points and post-hatch assays may reveal altered timing of astrocyte-state feature acquisition and altered thermoregulatory circuit dynamics in heat call-exposed offspring (Mariette and Buchanan, 2016).

Another notable result is the stress-thyroid interface suggested by transthyretin (*TTR*) regulation. In our snRNA-seq analysis, *TTR* shows a robust playback-associated signal: it is downregulated in heat call-exposed embryos in a glutamatergic cluster in the non-sex-stratified analysis, and in females it is downregulated across many clusters (20/49 clusters), while a *SIM1*-high glutamatergic population consistent with PVN identity shows sexually dimorphic regulation (up in males, down in females; Figure 2A-B, Supplemental Tables 3-5). This signature provides a mechanistic link between prenatal heat call playbacks and neuroendocrine programs, because PVN circuitry is a canonical hub for coordinating endocrine and homeostatic responses to environmental challenge (Herman and Tasker, 2016; Swanson and Sawchenko, 1980), and because hypothalamic TTR itself can modulate PVN neuropeptide signaling that governs feeding, metabolism, and stress-adaptive physiology (Zheng et al., 2016).

Mechanistically, TTR is best understood as a key component of thyroid hormone distribution to the brain via the blood-cerebrospinal fluid (CSF) interface. In vertebrates, the choroid plexus is a major site of TTR synthesis and secretion into CSF, where TTR contributes to the transport of thyroid hormone during period of brain development (Richardson et al., 2015). *TTR* expression in the hypothalamus has been shown to be sensitive to early-life peripubertal stress in a sex-dependent manner. Peripubertal stress in male mice drastically reduced hypothalamic *TTR* (via promoter hypermethylation) while simultaneously increasing prefrontal cortex *TTR*, and this sex-biased hypothalamic suppression is sufficient to reduce local thyroid hormone availability and alter stress-responsive behavior in adulthood (Rawat et al., 2022). In rats, psychosocial stress can also regulate *TTR* expression in the choroid plexus via glucocorticoids acting on mineralocorticoid and glucocorticoid receptors, providing a testable route through which anticipatory acoustic cues could engage endocrine-relevant transport machinery. While our data do not establish changes in thyroid hormone levels *in ovo*, or in the embryo itself, the changes in *TTR* expression associated with heat call playback are consistent with the specific hypothesis that heat-call exposure modulates thyroid-hormone buffering or delivery capacity in a sex-dependent manner, potentially interacting with hypothalamic maturation, metabolic programming, and stress-axis development. This *TTR*-based endocrine interface also provides a mechanistic link to our astrocyte-centric findings: changes in local thyroid hormone availability can shift the timing of thyroid hormone-dependent developmental programs (Bernal, 2007), raising the possibility that the neuronal *TTR* signature and astrocyte developmental-state program reflect a coordinated, thyroid-sensitive response to prenatal heat-call exposure.

The convergence of *TTR* downregulation and altered regulation of astrocyte developmental state in heat-call-exposed embryos suggests a potentially coordinated developmental program linking thyroid hormone availability to glial differentiation programs. Thyroid hormone signaling is a well-established regulator of neural progenitor development and differentiation, with T3 acting directly on neural stem cells through thyroid hormone receptors (TRα1) to modulate JAK-STAT signaling, thereby influencing neuronal versus glial fate decisions (Bernal, 2007; Chen et al., 2012; Mohácsik et al., 2011). While the directional effects of T3 on progenitor fate choices are context- and stage-dependent, with embryonic neural stem cells showing enhanced neuronal differentiation and suppressed astrocytic fate under T3 exposure (Chen et al., 2012), T3 robustly promotes the morphological differentiation and functional maturation of committed astrocytes, including upregulation of *GFAP*, cytoskeletal remodeling, and acquisition of mature astrocytic morphology (Lima et al., 1997; Mohácsik et al., 2011; Trentin and Moura Neto, 1995). In the developing brain, local thyroid hormone availability is actively regulated by cell-type-specific deiodinases: astrocytes and tanycytes express *DIO2*, which generates T3 from T4 (Guadaño-Ferraz et al., 1997), while neurons express *DIO3*, which degrades T3 (Mohácsik et al., 2011), providing spatial and temporal control of thyroid signaling during critical developmental windows. It remains unknown whether the *TTR* downregulation that we observe in PVN neurons alters local thyroid hormone availability and whether such changes contribute to altered astrocyte developmental-state regulation. However, the coordinated regulation of *TTR* (in PVN neurons) and gliogenic chromatin remodeling (in astrocytes) in response to the same prenatal acoustic stimulus raises the possibility that heat-call exposure engages thyroid-sensitive developmental programs in multiple cell types to coordinate hypothalamic developmental regulation in anticipation of a predicted thermal challenge. Testing this hypothesis will require direct measurement of thyroid hormone levels, cell-type-specific profiling of deiodinase expression, and functional manipulation of thyroid signaling during the exposure window.

Our findings establish that prenatal acoustic stress induces cell-type-specific chromatin remodeling and transcriptional reprogramming in the embryonic hypothalamus. Several considerations frame interpretation and prioritize future research directions. First, our E13 multiome snapshot cannot address whether heat-call-associated chromatin changes persist post-hatch or whether their developmental timing has functional consequences; longitudinal profiling across critical post-hatch windows will be necessary to address these questions. Second, validating the *TTR* finding requires protein-level confirmation in micro-dissected PVN tissue, direct measurement of T3/T4 levels in the brain and plasma, and assessment of *TTR* promoter accessibility. We note that rigorous single-nucleus quality control, including identification of a discrete choroid plexus epithelial cluster (marked by *TTR* and *ENPP2*) and absence of choroid marker contamination in neuronal populations, excludes dissection artifacts as a source of the PVN-specific *TTR* signal (Figure 2A, Supplemental Table 4) (Olney et al., 2022). Third, while motif enrichment and network architecture show NFI, AP-1, and Notch-linked programs as upstream regulators of the heat call responses, establishing direct transcription factor binding and causal hierarchies will require targeted epigenomic approaches (e.g., CUT&Tag (reference) in sorted PVN-lineage nuclei) paired with pathway-specific perturbations. Fourth, whether altered PVN *TTR* expression affects local thyroid hormone availability, neuropeptide processing (Liz et al., 2009; Zheng et al., 2016), or both, and how these molecular changes translate to physiological outcomes, remains to be determined through integrated developmental, endocrine, and behavioral phenotyping. Collectively, this work provides the first chromatin-resolution atlas of early-life stress programming in the developing avian hypothalamus and identifies *TTR* as a novel stress-responsive locus in neuroendocrine circuits.

In summary, this work establishes chromatin remodeling as a mechanistic basis for acoustic developmental programming and identifies astrocytes as primary cellular targets of prenatal sensory experience. We report three conceptual advances. First, the astrocyte-selective response challenges neuron-centric models of developmental plasticity and positions glial cells as active integrators of environmental anticipation signals. Second, extensive enhancer-promoter rewiring at gliogenic master regulators with limited concurrent transcriptional output supports an epigenetic priming mechanism in which prenatal experience presets the regulatory potential realized under subsequent thermal challenge, consistent with predictive adaptive response frameworks (Gluckman, 2005). Third, the *TTR* signature provides a candidate link between acoustic experience and thyroid-dependent developmental timing, suggesting that sensory programming may coordinate glial and neuronal developmental programs by modulating hormone availability. This study reveals that developmental programming operates through cell-type-specific chromatin reorganization that reshapes cellular competence landscapes, providing a molecular framework for understanding how prenatal acoustic cues prepare offspring for predicted environmental challenges.

## 4 Methods

**4.1 Experimental Design**

To conduct playback experiments on zebra finch embryos, we allowed adult zebra finches to pair and breed in an indoor aviary at Clemson University, USA. We transferred viable eggs at embryonic development stage 15 (Embryonic Day 2.75) to a shared single artificial incubator (Ovation EX, Brinsea) maintained at 37.5°C and 60% humidity to equalize the immediate pre-playback environment. On the evening of incubation day 8 (E8), we randomly assigned eggs to one of two experimental incubators (Octagon eco, Brinsea), each equipped with a different sound playback treatment. The control group (n = 20 embryos) received playback of whine calls combined with tet calls, while the experimental group (n = 20 embryos) received playback of whine calls combined with heat calls. Whine calls are complex contact calls with acoustic structures that facilitate auditory system development (Boucaud, 2016), whereas tet calls are typical contact calls commonly uttered by parents at the nest but unrelated to temperature.

Prenatal heat call playbacks occurred daily from E9 to E13 for 9 hours per day (9 am to 6 pm) at intensities of 60-70 dB. We delivered playbacks via speakers (PUI Audio Speaker 3W 8OHM 85DB 300 HZ, Part Number AS05508MO) positioned inside each experimental incubator and externally connected to an amplifier (Facmogu F900s) and an Apple iPad (9^th^ generation) using automated playbacks. All eggs were within 5cm of the speaker, and we rotated the incubation trays daily to provide a similar acoustic exposure to all eggs in the tray. The embryos were exposed to a 77-minute sequence of respective playbacks that looped continuously over the 9-hour playback period, thus replicating the acoustic experience of natural incubation (Katsis et al., 2018; Mariette and Buchanan, 2016). This looped playback consisted of randomly alternating sequences of whine calls (2 minutes each, 6 minutes total) and either heat calls or control calls (5 minutes each, 20 minutes total), separated by periods of silence (Katsis et al., 2018; Mariette and Buchanan, 2016). On the morning of E13, the day before hatching, we extracted embryos from their eggs and euthanized them by rapid decapitation. We flash froze the isolated heads in optimal cutting temperature (OCT) embedded blocks and stored at −80°C until needed for hypothalamic tissue collection.

### 4.2 Nuclei Isolation and Preparation for Single Nucleus RNA/ATAC-Sequencing

To prepare the stored hypothalamic tissue for downstream sequencing, we designed an optimized nuclei isolation protocol to preserve nuclear integrity and minimize RNA degradation. We first collected medial hypothalamic tissue samples from the frozen heads using a double-punch technique while cutting 100 μm coronal sections on a cryostat (Leica CM3050 S). Specifically, we made two circular punches, each 1 mm in diameter, in a vertically aligned, overlapping configuration resembling a “figure-8 shape”. We punched the tissue when the third ventricle became visible, approximately 5,000 μm caudal to the beak, collected in cryotubes on dry ice, and stored in -80C. We pooled tissue samples according to sex and experimental condition (n = 4 pooled heat call playback, 4 pooled control call playback samples). Each pooled sample contained five biological tissue punches, and we prepared two pooled replicates for each sex and condition combination.

We lysed the pooled samples for 5 minutes using 500 μL chilled 0.1x lysis buffer [10 mM Tris-HCl, 10 mM NaCl, 3 mM MgCl2, 0.01% Tween-20, 0.01% Igepal CA-630, 0.001% Digitonin, 1% Bovine Serum Albumin (BSA), 1mM dithiothreitol (DTT), RNase inhibitor (40 U/μL), and nuclease free water] and 15 homogenization strokes. We washed the lysed tissue with 500 μL wash buffer [10 mM Tris-HCl, 10 mM NACl, 3 mM MgCl2, 1% BSA, 0.1% Tween-20, 1 mM DTT, RNase inhibitor (40 U/μL), and nuclease free water] and filtered using Flowmi 70 μm cell strainer such that only 330 μL of nuclei suspension was passed through each 70 μm cell strainer at a time. Next, we passed the nuclei suspension through a Flowmi 40 μm cell strainer once, centrifuged at 500 rcf at 4°C for 5 minutes, and resuspended it in 1 mL cold wash buffer and pipette mixed 5 times. This was followed by a final round of cell straining using a pluriStrainer Mini 20 µm Cell Strainer, centrifugation at 500 rcf at 4°C for 5 minutes, and resuspension in chilled 1% BSA in 1x Phosphate-Buffered Saline (PBS) to obtain a final nuclei concentration of 8,000 nuclei/uL. The suspension thus obtained was quantified (target concentration between 3,200 and 8,000 nuclei/µL) on a Countess 3 FL Cell Counter and assessed for intact nuclei quality under a fluorescence microscope using DAPI staining.

### 4.3 Single Nucleus ATAC-seq and gene expression library preparation

To construct ATAC and gene expression libraries, we used the 10x Genomics Chromium Next GEM Single Cell Multiome ATAC + Gene Expression Reagent kit (v1, 10x Genomics, Pleasanton, CA), following the manufacturer’s protocol (CG000338, Rev G). Briefly, we transposed the the extracted nuclei with ATAC Buffer B and ATAC Enzyme B at 37°C for 60 minutes. We performed simultaneous barcoding of transposed ATAC fragments and mRNA-derived cDNA on the Chromium iX instrument by generating Gel Beads-in-emulsion (GEM). GEMs were obtained by combining transposed nuclei with a barcoding master mix, barcoded gel beads, and partitioning oil that were loaded onto a Chromium Next GEM Chip J. We obtained purified nucleic acids using Dynabeads MyOne SILANE and SPRIselect beads. The purified material underwent pre-amplification PCR and was split for ATAC and gene expression library construction. Next, ATAC libraries were generated by performing sample index PCR using SI-PCR Primer B and Amp Mix (14-16 cycles), followed by double-sided SPRIselect size selection. Gene expression libraries were generated by PCR amplifying cDNA for 12 cycles, fragmentation, end repair, A-tailing, adaptor ligation, indexing PCR (14-16 cycles), and double-sided SPRIselect size selection. We performed fragment quality control using the Agilent TapeStation High Sensitivity D5000 (ATAC) and High Sensitivity D1000 (gene expression) kits (Agilent Technologies, Santa Clara, CA). We performed library concentration quality control using the Qubit 1X dsDNA HS kit (Invitrogen, Waltham, MA). We then pooled, denatured, diluted as per manufacturer’s instructions, and sequenced the two libraries on the Illumina NovaSeq X+ (Illumina, San Diego, CA) (Hao et al., 2024).

### 4.4 Data processing and quality control

We demultiplexed the BCL files generated from the multiome sequencing and converted them to Fastq format using BCL Convert (version 4.1.23). We used 10x Cellranger-arc (v.2.0.2) to align ATAC and gene expression reads to the reference genome (bTaeGut1.4.pri) as well as to filter, count barcodes, and call peaks. We imported gene expression and chromatin accessibility profiles generated from Cellranger-arc into Seurat (v 5.2.1) (Hao et al., 2024) and Signac (v 1.14.0) (Stuart et al., 2021) respectively. We performed quality control on gene expression data by filtering cells based on gene count (200-3,500), unique molecular identifier (UMI) count (500-25,000), and mitochondrial content (< 5%). For chromatin accessibility data, we merged sample-specific peaks, and performed quality control by retaining specific peak length (20 bp - 10,000 bp). We retained cells for downstream single-nucleus ATAC-seq analysis based on total number of ATAC fragments (100-100,000 fragments), fraction of reads in peaks (> 15%), nucleosome signal (< 4), and Transcription Start Site (TSS) enrichment (> 2). After quality control, 57,388 of 60,138 single nuclei were retained for RNA-seq (average 1,714.4 RNA UMIs per nuclei), and 70,720 of 76,730 nuclei were retained for ATAC-seq (average 3,657.1 unique ATAC fragments per nuclei, with 35.6% of fragments overlapping called peaks).

To normalize gene expression count data, we used SCTransform (Hafemeister and Satija, 2019) and principal component analysis (PCA) was performed. We used the first 35 principal components in the downstream analyses. For ATAC-seq chromatin accessibility data, we normalized chromatin peaks using the term frequency-inverse document frequency (TF-IDF) transformation, implemented via the RunTFIDF function in Signac. The top 20% of the most frequently observed peaks were retained in subsequent analysis using the FindTopFeatures function. Next, we performed singular value decomposition (SVD) on the normalized and feature-selected matrix using the RunSVD function in Signac. The top 50 latent semantic indexing (LSI) components were used in downstream analysis and we excluded the first LSI, which is typically associated with technical variation such as sequencing depth (Stuart et al., 2021).

### 4.5 Clustering and visualization

To identify cell populations and visualize data, we used graph-based approaches on single and multi-modal datasets. For single modality data, we constructed shared nearest neighbor graphs using the FindNeighbors function in Seurat, applying PCA for gene expression data and LSI for chromatin accessibility data as described previously. We then performed clustering using the original Louvain algorithm for RNA-seq, and Smart Local Moving algorithm for ATAC-seq via the FindClusters function. We tested a range of resolutions for optimum cluster granularity, and we selected a final resolution of 1.1 (RNA) and 1.3 (ATAC) where cluster numbers stabilized. For multimodal analysis, we used the FindMultiModalNeighbors function in Seurat to construct a weighted nearest neighbor (WNN) graph that integrated RNA and ATAC dimensional reductions. The WNN graph (parameter setting: nn.name = “weighted.nn”) was used for multimodal clustering (Smart Local Moving algorithm and resolution = 0.6) and visualized with Uniform Manifold Approximation and Projection (UMAP).

To identify cluster-specific marker genes, we used the FindAllMarkers function with only.pos = TRUE. For differential gene expression, the MAST test was used with a minimum percentage detection threshold of 0.1, log-fold change threshold of 0.25, and genes with adjusted p-values < 0.05 were retained as cluster markers (positive values only). We obtained the top 20 cluster zebra finch markers, or their gene description (if unannotated for zebra finch), to identify clusters. We used these markers were used as input gene names for the Gene Expression tool on the CZ CELLxGENE database (organism = *Homo sapiens*; tissue = brain) and we obtained the cell types and count data for each of the 20 marker genes in a CSV file (CZI Single-Cell Biology Program et al., 2023). The frequency, average normalized expression, and percent of cells (number of cells expressing the gene / total cell count for cell type) expressing each marker gene across cell types were summarized and ranked to find the cell types most frequently associated with the marker set. This approach was supplemented by literature review of the marker genes and the cell types they are expressed in. Overall, the 49 clusters obtained were grouped into 13 hypothalamic clusters based on their cell-type annotation in the literature (Chaturvedi et al., 2025).

To perform astrocyte-specific analyses, we isolated the astrocyte cluster (n = 14,949 nuclei) from the integrated multimodal dataset based on cluster annotations. We performed subclustering using weighted nearest neighbor (WNN) integration of RNA principal components (PCs 1-30) and ATAC latent semantic indexing (LSI) dimensions (2-40). We computed WNN-based UMAP embeddings, and performed graph-based clustering across multiple resolutions (0.01-0.20) using the Leiden algorithm. We selected a resolution of 0.06 based on elbow plot analysis, yielding biologically interpretable subclusters. The resulting subclusters and WNN-based UMAP embedding were used as the basis for downstream visualization and transcription factor regulon analyses in astrocyte-associated subpopulations. This approach identified four transcriptionally distinct astrocyte-associated subpopulations with marker profiles suggestive of different developmental states: subcluster 0, *FOXG1*/*KCNQ5*-positive progenitor-like astrocyte-associated cells (5,885 nuclei); subcluster 1, *THRB1*-positive developing astrocyte-associated cells (4,205 nuclei); subcluster 2, *AURKA*-positive proliferative astrocyte-associated precursor-like cells (2,439 nuclei); and subcluster 3, *SHH*/*NKX2-4*/*SLIT2*-positive ventral gliogenic-like astrocyte-associated cells (2,420 nuclei) (Figure 3B). We ascribed these identities based on marker combinations consistent with astroglial developmental progression rather than any single marker alone, with *FOXG1* linked to astrocyte lineage diversity, *THRB1* consistent with thyroid hormone-dependent astrocyte maturation, the *AURKA*-positive subcluster also showing a proliferative/cell-cycle signature consistent with precursor-like cells, and *SHH/SLIT2* expression consistent with ventral astroglial programs (Supplemental Table 8) (Bose et al., 2025; Graziano et al., 2024; Guadaño-Ferraz et al., 1997; Li et al., 2025; Minocha et al., 2015; Minocha et al., 2017; Trentin, 2006; Wang et al., 2011).

### 4.6 Differential Accessibility and Gene Expression Analyses

To identify gene expression and chromatin differences between heat call and control exposed embryos, we performed analyses at three levels: globally (across all cell types), within each cell cluster, and stratified by sex. We achieved this by creating a combined identity variable by combining cluster, condition and sex (i.e., stratified by sex), or by combining cluster and condition (i.e., cluster wide, non-sex stratified). For each analysis, we used the MAST test with adjusted p-value < 0.05, and an absolute log2 fold change (log_2_FC) threshold as appropriate for the analysis context. We corrected p-values for multiple comparisons using Bonferroni adjustment as implemented in Seurat’s FindMarkers function (p.adjust.method = ‘bonferroni’). We annotated differentially accessible chromatin peaks using the ClosestFeature function from Signac and bedtools intersect to identify peaks overlapping gene features (i.e., within gene bodies and regulatory region). We visualized the results from these analyses using dot matrix plots to assess gene expression and chromatin accessibility differences across all clusters.

### 4.7 Single-Nucleus Transcription Factor Activity Analysis

To investigate gene regulatory mechanisms underlying differential chromatin accessibility, we analyzed single-nucleus transcription factor activity on condition-specific differentially accessible peaks. This analysis employed the ChromVAR-based differential transcription factor activity estimation and motif scanning by leveraging the JASPAR2020 vertebrate motif database and the zebra finch reference genome (bTaeGut1.4.pri assembly). We obtained position frequency matrices for vertebrate transcription factors from JASPAR 2020 and we used them to scan open chromatin regions in the ATAC assay. We indexed the reference genome for sequence scanning and GC content calculation using RegionStats function in Signac to control for sequence composition bias. We used CreateMotifMatrix to scan motifs in ATAC peaks, and assessed differential motif activity between heat call and control using FindMarkers on the ChromVar assay. We visualized results using motif plots, violin plots and feature plots using SCpubr (v 2.0.2) (Blanco-Carmona, 2022).

### 4.8 High-dimensional Weighted Gene Co-Expression Network Analysis

To identify cell-type-specific gene programs associated with heat call playback, we performed high-dimensional weighted gene co-expression network analysis (hdWGCNA) on each annotated hypothalamic cell type using the hdWGCNA R package (v 0.4.07) (Morabito et al., 2023). We normalized RNA expression counts for all nuclei (n = 55,950 nuclei) using log-normalization with a scale factor of 10,000. We identified the top 2,000 genes with the highest variance across the integrated dataset using variance-stabilizing transformation. For each cell type, we isolated clusters based on WNN-derived cluster annotations and subset for independent network analysis. We aggregated transcriptionally similar cell types into metacells using the hdWGCNA framework to decrease the sparsity of the dataset. This metacells aggregation was based on similarity in the WNN embedding space and the grouping was stratified by the cell type and sample ID to preserve batch structure.

To construct the co-expression network, we retained genes expressed in at least 5% of nuclei across resulting in 10,901 genes expressed across all cell types. We scaled the expression values to zero mean and unit variance within each cell type before network construction. Gene selection for network construction employed fraction-based filtering, retaining genes expressed in at least 5% of nuclei within any annotated cell type across the complete hypothalamic dataset. This cross-cell-type selection strategy identified 10,901 genes for analysis, ensuring consistent gene sets across cell-type-specific networks while capturing both ubiquitously expressed and cell-type-enriched genes. Expression values for selected genes were scaled to zero mean and unit variance within each cell type prior to network construction. For each cell type, we constructed signed co-expression networks to preserve whether genes increase or decrease together. To optimize network structure, we selected soft-thresholding power independently for each cell type by testing a range of powers from 1 to 30. Soft-thresholding power that best achieved scale-free topology (R² ≥ 0.8) while maintaining reasonable connectivity levels (mean connections < 300) was picked for network construction. We computed topological overlap matrices from weighted adjacency matrices to quantify connection strength between genes based on shared network neighbors. Next, we identified modules using gene expression patterns and network neighborhoods using hierarchical clustering of topological overlap matrices. We defined modules using dynamic tree cutting with parameters optimized to detect specific, distinct gene programs: modules required at least 10 genes, we used aggressive splitting to identify specialized groups, and we merged modules that were more than 75% similar to avoid redundancy. We assigned modules cell-type-specific identifiers and genes not assigned to any module were designated as background (grey module). Finally, for each module, we calculated eigengenes as the first principal component of the module’s expression profile and removed technical variation between datasets by harmonizing the module eigengenes across all eight samples using Harmony batch correction.

To quantify relationships between co-expression modules and heat call playback across all cell types, we correlated harmonized module eigengenes with a binary treatment indicator (Control = 0, Heat Call Playback = 1) using Pearson correlation. We assessed statistical significance using Student’s asymptotic test, and adjusted p-values for multiple testing within each cell type using the Benjamini-Hochberg false discovery rate procedure. Modules with FDR < 0.05 and absolute correlation coefficient > 0.05 were considered significantly associated with heat call playback. We identified hub genes by calculating intramodular connectivity (kME) for functional interpretation of treatment-associated modules.

We performed gene ontology enrichment analysis on module gene lists to identify biological pathways and functional categories associated with heat call exposure (Ge et al., 2020). To mitigate incomplete functional annotation in the zebra finch genome, we mapped zebra finch gene symbols to human orthologs using the biomaRt R package (v 2.60.1) to query the Ensembl BioMart database (Ensembl release 110, *Taeniopygia guttata* genome assembly GCF_003957565.2_bTaeGut1.4.pri) (Kinsella et al., 2011). We analyzed genes with identifiable human orthologs using ShinyGO v0.85.1 (species: *Homo sapiens*). We provided a custom background gene set consisting of all 21,407 zebra finch genes in the RNA-seq analysis that successfully mapped to human orthologs (N = 11,310 unique human genes) to ensure the statistical universe matched measurable genes in our experiment. We tested enrichment against GO Biological Process, GO Molecular Function, GO Cellular Component, and KEGG pathway databases. In parallel, we performed enrichment analysis using native zebra finch annotation (ShinyGO v0.85.1, species: *Taeniopygia guttata*) with the zebra finch background gene set to assess concordance between approaches. For both analyses, we calculated enrichment p-values using the hypergeometric test and corrected using the Benjamini-Hochberg FDR method. Pathways meeting FDR < 0.05 were considered statistically significant. Pathway size limits were set to a minimum of 2 and a maximum of 5,000 genes. We enabled redundancy reduction to collapse highly overlapping pathways. We ranked results by fold enrichment, and top pathways per module were reported as bar plots displaying fold enrichment and FDR values.

### 4.9 Transcription Factor Regulatory Network Analysis

To identify transcription factors (TF) associated with heat call playback, we integrated TF binding site predictions with co-expression network analysis using hdWGCNA. We used the extreme gradient boosting algorithm (XGBoost) to predict TF-gene regulatory relationships by modeling each gene’s expression based on all TFs with binding motifs in that gene’s promoter region. This approach identifies which TFs are most predictive of each gene’s expression patterns by generating importance scores for each TF-gene pair: gain (how much the TF improves prediction accuracy), cover (proportion of cells influenced by the TF), and frequency (how often the TF is selected across models). We calculated Pearson correlations to distinguish activating (positive correlation) versus repressive (negative correlation) TF regulatory relationships.

From the complete transcription factor network, we selected high-confidence TF-target gene pairs, called “regulons”, by choosing the top-ranked TFs for each target gene based on XGBoost importance scores. We retained TF-gene pairs with regulatory scores ≥ 0.01 and limited each gene to its top 10 predicted TF regulators. Next, we computed regulon activity scores representing the aggregated expression of target genes within each TF’s regulon. We calculated separate regulon activity scores for positive regulons (positive correlation with TF expression, using default correlation thresholds) and negative regulons (correlation ≤ -0.05).

We compared regulon activity scores between heat call and control call playback embryos to identify coordinated changes in gene expression using Wilcoxon rank-sum tests with FDR correction using the Benjamini-Hochberg procedure. We classified TFs as heat call-responsive if they showed coordinated regulatory changes, specifically increased positive regulon activity coupled with decreased negative regulon activity (both FDR < 0.05). We prioritized these heat call-responsive TFs based on their membership in heat call-associated co-expression modules. Within each module, we ranked TFs by statistical significance (FDR) and effect size (log₂ fold-change in regulon activity) to identify key transcription factor regulators influencing gene expression in the associated co-expression module.

We visualized gene regulatory networks showing direct targets or secondary connections, with edge weights representing correlation strength. For gene ontology enrichment analysis, we prepared gene lists using permissive thresholds (correlation ≥ 0.005) to capture indirect regulatory connections. We performed gene ontology enrichment using ShinyGo v0.85.1 as previously described.

For *ASCL1* network visualization, we constructed the regulatory network at depth 2 to include *ASCL1*, its direct target genes, and secondary targets. The network was generated using correlation-weighted edges to represent the directionality of TF-target relationships (positive correlation: activation; negative correlation: repression). We scaled edge thickness by regulatory gain scores (XGBoost importance) and applied a permissive correlation threshold of 0.005 to capture indirect regulatory connections for visualization purposes. We plotted network using the TFNetworkPlot function from hdWGCNA with custom node labeling for *ASCL1* and key Notch pathway components (Supplemental Figure 4; *DLL1*, *NOTCH1*, *MAML3*).

### 4.10 Pseudotemporal Trajectory Analysis

To characterize condition-associated developmental trajectory structure in response to heat call playback, we performed pseudotemporal trajectory analysis using Slingshot v2.2.1 (Street et al., 2018). We converted the integrated Seurat object containing all hypothalamic cell types to SingleCellExperiment format, and provided weighted nearest neighbor (WNN) UMAP coordinates, which integrate transcriptomic (RNA-seq) and chromatin accessibility (ATAC-seq) information into a unified low-dimensional embedding, as the input dimensionality reduction space. We performed trajectory inference separately for control and heat call conditions on the same WNN UMAP embedding to assess condition-associated differences in pseudotime structure across the full cellular diversity of the hypothalamus. For each condition, we applied Slingshot to WNN UMAP coordinates using WNN cluster identities (resolution 0.6) to guide lineage construction. We designated Cluster 0 as the root state because it represented the largest balanced cluster with substantial representation from both conditions, minimizing sensitivity to condition-dependent cell abundance differences. We extracted pseudotime values from the principal lineage for each condition and were mapped back to the Seurat object metadata using cell barcodes as unique identifiers.

To quantify condition-associated differences in pseudotime distributions across hypothalamic cell populations, we performed per-cluster pseudotime comparisons between heat call and control conditions. For each WNN cluster, we extracted pseudotime values for all cells assigned to that cluster in each condition. We excluded clusters with fewer than 5 cells in either condition from statistical testing. For each cluster, we calculated mean pseudotime, cell counts per condition, and the pseudotime shift (Δpt) as the difference between heat call and control mean pseudotime values. We assessed statistical significance using two-sided Wilcoxon rank-sum tests comparing the pseudotime distributions between conditions within each cluster. We adjusted p-values for multiple testing across all clusters using the Benjamini-Hochberg method. Clusters with FDR < 0.05 were considered to exhibit significant condition-associated pseudotime shifts.

To identify genes whose expression varied along developmental trajectories, we used tradeSeq v1.8.0 in R (Van den Berge et al., 2020) to model pseudotime-associated expression patterns. For each condition, we fitted generalized additive models (GAMs) separately after retaining cells with valid pseudotime values and extracting their corresponding Slingshot curve weights for the principal lineage. We fitted GAMs using fitGAM with SCT-normalized expression values, pseudotime as the continuous variable, lineage curve weights, and 6 knots. We then applied associationTest, which tests whether average gene expression is associated with pseudotime rather than remaining constant across pseudotime within a lineage, to identify pseudotime-dynamic genes within each condition. Genes passing a p-value threshold of 1×10⁻⁴ were retained as significant pseudotime-dynamic genes.

For focused analysis of astrocyte trajectory-associated gene-expression dynamics, we subset the integrated Seurat object to astrocytes by identifying cells with astrocyte annotations in the cell type metadata. We retained pseudotime values from the full-dataset trajectories for astrocytes to maintain developmental context. To enable direct comparison of gene expression trajectories between conditions, we independently rescaled pseudotime values to a 0-1 range within each condition using min-max normalization. We visualized gene expression dynamics for astrocyte-state-development-associated genes (*NFIA* and *FABP7*) and Notch signaling genes (*ASCL1, DLL1, NOTCH1, MAML3*) as scatterplots using SCT-normalized expression values from astrocytes plotted against normalized pseudotime, overlaid with LOESS regression curves (span = 0.75) and 95% confidence intervals to capture smooth, non-linear expression trends along the developmental trajectory.

### 4.11 Chromatin Co-Accessibility and Regulatory Network Analysis

To identify playback-specific changes in chromatin architecture, we performed co-accessibility analysis using Cicero v1.3.9 with Monocle3 v1.4.26 (Pliner et al., 2018). We converted the integrated ATAC assay to a Monocle3 cell_data_set (CDS) format. We binarized ATAC-seq counts (counts greater than 0 were set to 1) to focus on accessibility patterns instead of signal intensity. We annotated the genome by extracting peak coordinates and chromosome information using the GRanges object associated with the ATAC assay, and obtained genome sizes from the indexed zebra finch reference genome (GCF_003957565.2_bTaeGut1.4.pri) FASTA file.

To find patterns of co-accessibility due to playbacks, we created a Cicero CDS with WNN UMAP coordinates as the reduced dimension input for cell aggregation to aggregate similar cells for robust co-accessibility estimation. To determine optimal parameters for co-accessibility estimation, we tested multiple parameter combinations varying distance constraints (100-250 kb), window sizes (500-750 kb), and smoothing parameters (s = 0.75-0.85). We compared results using Jaccard similarity indices to quantify overlap of detected connections (co-accessibility ≥ 0.15) across parameter sets, along with median co-accessibility scores, number of connections, and number of co-accessible modules (CCANs). Based on this sensitivity analysis, we found standard mammalian genome parameters as optimal: smoothing parameter s = 0.75, distance constraint of 250 kb, sliding window of 500 kb, and 100 samples per window. We generated co-accessibility models with the estimated distance parameters to estimate correlations in chromatin accessibility between peak pairs, and these correlations were reconciled into a single genome-wide network to produce peak co-accessibility scores. We identified co-accessible modules (CCANs) that represent putative *cis*-regulatory interactions, and we applied a co-accessibility score threshold of 0.15 to define high-confidence regulatory connections.

To identify playback-specific regulatory changes, we subset the analysis to astrocytes to investigate gene-regulatory programs in the cell cluster. We calculated peak accessibility, defined as the mean binarized accessibility across cells, separately for heat call and control call playbacks, and peaks with accessibility scores greater than 0.01 in each playback condition were retained for downstream analyses. We annotated these endpoint peak co-accessibility connections in heat call and control playback conditions, and connections where both endpoint peaks exhibited accessibility ≥ 0.05 in a given condition were considered active in that condition. To identify chromatin loops that are gained or lost in response to heat call playback, we explored chromatin rewiring in each playback condition, defining rewired connections as those that were active in one playback condition (both peaks ≥ 0.05 accessible), inactive in the other condition (at least one peak < 0.05 accessible), and had a co-accessibility score ≥ 0.15.

Further, we identified chromatin rewiring anchored at gene promoters (defined as ± 2 kb windows surrounding transcription start sites) using the zebra finch genome annotation (GCF_003957565.2_bTaeGut1.4.pri genomic GTF). We assigned regulatory connections to genes where at least one end point overlapped the promoter region of that gene, and for each gene, heat call playback-specific promoter-anchored connections were quantified along with the mean co-accessibility scores of those connections. We integrated these chromatin rewiring data with differential gene expression results from the snRNA-seq analysis to assess concordance between chromatin remodeling and transcriptional changes, and we quantified the relationship between the number of heat call-specific promoter connections and gene expression fold changes using Pearson correlation. To test for differences in RNA expression across rewiring burden groups (high: ≥ 10 promoter-anchored loops; moderate: 5-9 loops; low: 1-4 loops), we applied the Kruskal-Wallis rank sum test. Finally, we converted Cicero connections and CCANs to Signac-compatible formats, and we visualized locus-specific chromatin architecture using CoveragePlot from Signac, displaying ATAC-seq coverage, peak locations, gene annotations, and playback-specific chromatin connections for selected genomic regions.

To identify regulatory peaks undergoing chromatin state changes in astrocytes, we performed differential accessibility analysis on astrocyte ATAC-seq peaks using the MAST test in Seurat’s FindMarkers function, comparing heat call versus control playback. We classified peaks with FDR-adjusted p-values < 0.05 and absolute log₂ fold-change > 0.25 as differentially accessible. We classified peaks that were both (i) significantly gained in heat call playback (positive log₂FC, FDR < 0.05), and (ii) participating as endpoint *cis*-regulatory peaks in heat call-specific rewired Cicero connections as “dual-hit” peaks. These dual-hit peaks were used to identify regulatory peaks experiencing simultaneous change in chromatin accessibility, and likely three-dimensional genome organization. We further assessed regulatory relationships between dual-hit peaks and gene expression by performing peak-to-gene linkage analysis Signac’s LinkPeaks function. We computed Pearson correlations between ATAC-seq accessibility at each peak and SCT-normalized RNA expression of genes within 500 kb across all astrocytes for peak-gene pairs with at least 10 cells expressing both signals. We defined significant peak-gene correlations as those with |Pearson r| > 0.1 and p < 0.05. For genes with multiple linked peaks, we summarized the maximum absolute correlation, mean correlation, and minimum p-value. We generated locus-specific visualization of dual-hit peak architecture using Signac’s CoveragePlot, highlighting dual-hit peaks and displaying playback-specific Cicero connections as arc diagrams overlaid on ATAC-seq coverage tracks.

### 4.12 Use of AI Tools

We employed large language models accessed via the Perplexity platform (including Grok 4.1, GPT 5.1, Sonar, and Claude Sonnet 4.5) to assist with R script optimization and debugging of computational workflows, including snRNA/ATAC-seq, hdWGCNA, trajectory analysis, and Cicero co-accessibility pipelines. We also used these models to help edit and refine portions of the manuscript, particularly the Materials and Methods and figure legends, to ensure adherence to the target journal’s formatting and reporting guidelines. All AI-assisted code and text were critically reviewed, modified, and validated prior to inclusion in the manuscript, and the authors take full responsibility for the accuracy and integrity of all content. No AI tools were used for image creation, editing, or enhancement.

## 6 Author Contributions

P.S. performed embryonic dissections, hypothalamic punching, nuclei isolation, and tissue collection for snRNA/ATAC-seq, conceived the bioinformatic analyses, performed all computational and statistical analyses, generated figures and tables, and led the drafting and revision of the manuscript.

K.W. performed nuclei isolation and barcoding, library preparation, and sequencing on the 10x Chromium platform, and contributed to manuscript editing. V.S. contributed to conceptualization and experimental design, conceived and supervised computational, bioinformatic, and statistical analyses, contributed to data interpretation, and co-wrote and edited the manuscript. P.S.F. supervised nuclei isolation and quality control steps prior to library preparation, supervised aspects of laboratory work, and contributed to manuscript editing. T.F.C.M. contributed to conceptualization and experimental design, supervised aspects of laboratory work, provided access to laboratory space and 10x Chromium instrumentation, supported the sequencing experiments, obtained funding, contributed to data analysis and interpretation, and co-wrote and edited the manuscript. N.A.S. performed embryonic dissections, sex identification, and tissue collection for snRNA/ATAC-seq, assisted in bioinformatic analysis for cell cluster annotation, and co-wrote and edited the manuscript. S.R.L. performed embryonic dissections, sex identification, and tissue collection for snRNA/ATAC-seq, and contributed to manuscript editing. A.M.R. performed embryonic dissections, sex identification, and tissue collection for snRNA/ATAC-seq, led the sound playback design for incubators, and contributed to manuscript editing. J.R. performed embryonic dissections, sex identification, and tissue collection for snRNA/ATAC-seq, and contributed to manuscript editing. J.M.G. conceived and supervised the overall project, contributed to experimental design, performed embryonic dissections and nuclei isolation, contributed to data analysis and interpretation, obtained funding, and co-wrote and edited the manuscript. All authors read and approved the final manuscript.

## 7 Acknowledgments

Clemson Institute for Human Genetics Research Computing is acknowledged for their generous allotment of compute time on the Secretariat Cluster. David Clayton is acknowledged for their contribution to data interpretation. Townsend Mcdonald is acknowledged for their contribution to the histological analysis of these results.

## 9 Ethics Approval

All procedures were approved by the Clemson University Institutional Animal Care and Use Committee (IACUC), under protocols AUP2022-0486 and AUP2023-0106.

## 10 Data Accessibility Statement

Raw single-nucleus multiome sequencing data (snRNA-seq and snATAC-seq), along with processed gene expression and chromatin accessibility matrices, have been deposited in NCBI’s Gene Expression Omnibus (GEO) under accession GSE322480 and are publicly available at https://www.ncbi.nlm.nih.gov/geo/query/acc.cgi?acc=GSE322480. Raw sequencing reads are available through the NCBI Sequence Read Archive under BioProject accession PRJNA1429414.

Sample metadata, RNA count matrices, ATAC peak matrices, and processed Seurat/Signac objects are available as Supplemental files linked to the GEO submission. Analysis code is publicly available at https://github.com/praxsubba/Heat_Call_Single_Nucleus_Multiome. All other relevant data are included in the article and its Supplemental information.

