## Supplemental Data for "Multiome Profiling Reveals Astrocyte and Neuroendocrine Targets of Prenatal Acoustic Programming in Zebra Finch Embryos"

**Supplemental Table 1:** Per-sample quality control (QC) summary for the snATAC-seq dataset, including nuclei counts before and after filtering and median ATAC-seq QC metrics for each sample across heat call and control playback conditions.

| Sample | Condition | Sex | ATAC Nuclei Pre-QC | ATAC Nuclei Post-QC | ATAC Median Count | ATAC Median TSS Enrichment | ATAC Median % Reads in Peaks | ATAC Median Nucleosome Signal | ATAC Median Total Fragments | ATAC Median Peak Fragments | ATAC Median TSS Fragments |
|---|---|---|---|---|---|---|---|---|---|---|---|
| **S1** | Heat Call | Male | 13379 | 12586 | 2256.00 | 4.49 | 30.83 | 0.57 | 5916.50 | 1948.00 | 1732.50 |
| **S2** | Heat Call | Female | 8761 | 8155 | 3694.00 | 4.36 | 30.83 | 0.53 | 9430.00 | 3132.00 | 2847.00 |
| **S3** | Control | Male | 9513 | 8196 | 2653.50 | 4.21 | 27.15 | 0.57 | 7439.00 | 2147.50 | 2109.00 |
| **S4** | Control | Female | 7985 | 7486 | 3105.00 | 4.62 | 32.51 | 0.56 | 7851.50 | 2646.00 | 2358.00 |
| **S5** | Heat Call | Male | 9575 | 8842 | 3754.00 | 4.92 | 36.07 | 0.54 | 9047.00 | 3296.00 | 2894.00 |
| **S6** | Heat Call | Female | 8635 | 7859 | 3654.00 | 4.64 | 32.99 | 0.60 | 9305.00 | 3100.00 | 2814.00 |
| **S7** | Control | Male | 10918 | 10245 | 2633.00 | 5.08 | 36.47 | 0.60 | 6258.00 | 2314.00 | 1982.00 |
| **S8** | Control | Female | 7964 | 7351 | 3301.00 | 4.80 | 34.22 | 0.66 | 8103.00 | 2853.00 | 2508.00 |

**Supplemental Table 2**: Per-sample quality control (QC) summary for the snRNA-seq dataset, including RNA nuclei counts before (Pre-QC) and after (Post-QC) filtering, and median RNA QC metrics - including median RNA nuclei count, median nFeature (number of unique genes detected per nucleus), and median % mt (percentage of mitochondrial gene reads) - for each sample across heat call and control playback conditions.

| Sample | Condition | Sex | RNA Nuclei Pre-QC | RNA Nuclei Post-QC | Median RNA Nuclei Count | RNA Median nFeature | RNA Median % mt |
|---|---|---|---|---|---|---|---|
| S1 | Heat Call | Male | 10749 | 9864 | 1652 | 1064 | 0 |
| S2 | Heat Call | Female | 6676 | 6501 | 1852 | 1144 | 0 |
| S3 | Control | Male | 7201 | 6724 | 1565 | 997 | 0 |
| S4 | Control | Female | 6335 | 6023 | 1676 | 1061 | 0 |
| S5 | Heat Call | Male | 7462 | 7232 | 1747 | 1074 | 0 |
| S6 | Heat Call | Female | 6568 | 6380 | 1729.5 | 1090.5 | 0 |
| S7 | Control | Male | 8660 | 8463 | 1694 | 1070 | 0 |
| S8 | Control | Female | 6487 | 6201 | 1773 | 1106 | 0 |

**Supplemental Table 3**: Summary of WNN clusters showing numeric cluster identity, cell type annotation, total nuclei per cluster, and nuclei counts by condition and sex.

| WNN Cluster | Cell Type Annotation | Number Nuclei Total | Number Heat Call Nuclei | Number Control Nuclei | % Heat Call Nuclei | % Control Nuclei | Number Male Nuclei | Number Female Nuclei |
|---|---|---|---|---|---|---|---|---|
| 0 | Glutamatergic Neuron | 3878 | 1955 | 1923 | 50.4 | 49.6 | 2297 | 1581 |
| 1 | Astrocyte | 3277 | 1801 | 1476 | 55 | 45 | 1683 | 1594 |
| 2 | Astrocyte | 2988 | 1556 | 1432 | 52.1 | 47.9 | 1621 | 1367 |
| 3 | Glutamatergic/GABA | 2871 | 1451 | 1420 | 50.5 | 49.5 | 1625 | 1246 |
| 4 | Glutamatergic Neuron | 2812 | 1358 | 1454 | 48.3 | 51.7 | 1671 | 1141 |
| 5 | Astrocyte | 2444 | 1377 | 1067 | 56.3 | 43.7 | 1391 | 1053 |
| 6 | GABAergic Neuron | 2343 | 1306 | 1037 | 55.7 | 44.3 | 1317 | 1026 |
| 7 | Glutamatergic Neuron | 2119 | 1084 | 1035 | 51.2 | 48.8 | 1212 | 907 |
| 8 | Glutamatergic Neuron | 2053 | 1005 | 1048 | 49 | 51 | 1143 | 910 |
| 9 | GABAergic Neuron | 1994 | 1056 | 938 | 53 | 47 | 986 | 1008 |
| 10 | GABAergic Neuron | 1965 | 1138 | 827 | 57.9 | 42.1 | 1085 | 880 |
| 11 | Astrocyte | 1876 | 1117 | 759 | 59.5 | 40.5 | 1048 | 828 |
| 12 | Glutamatergic Neuron | 1548 | 814 | 734 | 52.6 | 47.4 | 850 | 698 |
| 13 | GABAergic Neuron | 1515 | 815 | 700 | 53.8 | 46.2 | 807 | 708 |
| 14 | Glutamatergic Neuron | 1469 | 745 | 724 | 50.7 | 49.3 | 837 | 632 |
| 15 | Astrocyte | 1467 | 751 | 716 | 51.2 | 48.8 | 828 | 639 |
| 16 | Glutamatergic Neuron | 1401 | 650 | 751 | 46.4 | 53.6 | 770 | 631 |
| 17 | GABAergic Neuron | 1326 | 793 | 533 | 59.8 | 40.2 | 737 | 589 |
| 18 | GABAergic Neuron | 1294 | 708 | 586 | 54.7 | 45.3 | 667 | 627 |
| 19 | Glutamatergic/GABA | 1277 | 649 | 628 | 50.8 | 49.2 | 658 | 619 |
| 20 | Astrocyte | 1189 | 613 | 576 | 51.6 | 48.4 | 705 | 484 |
| 21 | Microglia | 1156 | 624 | 532 | 54 | 46 | 658 | 498 |
| 22 | Oligodendrocyte | 992 | 521 | 471 | 52.5 | 47.5 | 523 | 469 |
| 23 | Astrocyte | 877 | 490 | 387 | 55.9 | 44.1 | 536 | 341 |

| WNN Cluster | Cell Type Annotation | Number Nuclei Total | Number Heat Call Nuclei | Number Control Nuclei | % Heat Call Nuclei | % Control Nuclei | Number Male Nuclei | Number Female Nuclei |
|---|---|---|---|---|---|---|---|---|
| 24 | GABAergic Neuron | 814 | 430 | 384 | 52.8 | 47.2 | 408 | 406 |
| 25 | Astrocyte | 713 | 386 | 327 | 54.1 | 45.9 | 401 | 312 |
| 26 | Glutamatergic/GABA | 702 | 67 | 635 | 9.5 | 90.5 | 667 | 35 |
| 27 | Ependymal Cell | 633 | 339 | 294 | 53.6 | 46.4 | 341 | 292 |
| 28 | Glutamatergic Neuron | 628 | 314 | 314 | 50 | 50 | 304 | 324 |
| 29 | Glutamatergic Neuron | 607 | 344 | 263 | 56.7 | 43.3 | 318 | 289 |
| 30 | Astrocyte | 540 | 270 | 270 | 50 | 50 | 303 | 237 |
| 31 | Glutamatergic Neuron | 505 | 282 | 223 | 55.8 | 44.2 | 330 | 175 |
| 32 | Fibroblast | 431 | 206 | 225 | 47.8 | 52.2 | 268 | 163 |
| 33 | Astrocyte | 417 | 219 | 198 | 52.5 | 47.5 | 218 | 199 |
| 34 | Glutamatergic Neuron | 383 | 222 | 161 | 58 | 42 | 282 | 101 |
| 35 | Glutamatergic Neuron | 368 | 235 | 133 | 63.9 | 36.1 | 271 | 97 |
| 36 | Astrocyte | 362 | 201 | 161 | 55.5 | 44.5 | 196 | 166 |
| 37 | Macrophage | 341 | 178 | 163 | 52.2 | 47.8 | 169 | 172 |
| 38 | Capillary Endothelial | 328 | 125 | 203 | 38.1 | 61.9 | 205 | 123 |
| 39 | Glutamatergic Neuron | 306 | 136 | 170 | 44.4 | 55.6 | 163 | 143 |
| 40 | Endothelial Cell | 269 | 87 | 182 | 32.3 | 67.7 | 140 | 129 |
| 41 | Astrocyte | 266 | 157 | 109 | 59 | 41 | 133 | 133 |
| 42 | Vascular Associated Smooth Muscle Cell | 247 | 108 | 139 | 43.7 | 56.3 | 158 | 89 |
| 43 | Endothelial Cell | 210 | 110 | 100 | 52.4 | 47.6 | 121 | 89 |
| 44 | GABAergic Neuron | 193 | 120 | 73 | 62.2 | 37.8 | 106 | 87 |
| 45 | Glutamatergic Neuron | 187 | 96 | 91 | 51.3 | 48.7 | 105 | 82 |
| 46 | Glutamatergic Neuron | 138 | 67 | 71 | 48.6 | 51.4 | 85 | 53 |
| 47 | Glutamatergic Neuron | 128 | 63 | 65 | 49.2 | 50.8 | 74 | 54 |
| 48 | Choroid Plexus Epithelial Cell | 103 | 53 | 50 | 51.5 | 48.5 | 28 | 75 |

**Supplemental Table 4: WNN cluster marker genes and cell type annotations.** The top 20 differentially expressed marker genes defining each of 49 WNN clusters in the E13 zebra finch hypothalamus are given. These markers were identified using FindAllMarkers (MAST; adjusted P < 0.05, |log2FC| ≥ 0.25, pct.1 ≥ 0.1) and ranked by log2 fold-change. Cell type annotations were assigned by cross-referencing marker genes with human ortholog expression in brain tissue (CZ CELLxGENE Discover) and literature-based validation; clusters were grouped into 13 hypothalamic cell types. Columns: Gene (marker gene symbol), P-Value (unadjusted P-value from MAST), Average log2FC (average log2 fold-change for the cluster versus all other clusters), %.1 (fraction of cells in the cluster expressing the gene), %.2 (fraction of cells in all other clusters expressing the gene), FDR (FDR-adjusted P-value), cluster (WNN cluster ID), and Annotation (assigned cell type).

| Gene | P-Value | FDR | Average $log_2FC$ | Fraction 1 | Fraction 2 | Cluster | Annotation |
|---|---|---|---|---|---|---|---|
| ***LOC115496780*** | 0.00E+00 | 0.00E+00 | 2.51 | 0.60 | 0.14 | 0 | Glutamatergic Neuron |
| ***LOC105759036*** | 0.00E+00 | 0.00E+00 | 2.50 | 0.34 | 0.06 | 0 | Glutamatergic Neuron |
| ***EYS*** | 0.00E+00 | 0.00E+00 | 1.96 | 0.58 | 0.18 | 0 | Glutamatergic Neuron |
| ***NETO1*** | 0.00E+00 | 0.00E+00 | 1.94 | 0.43 | 0.15 | 0 | Glutamatergic Neuron |
| ***LOC100221646*** | 0.00E+00 | 0.00E+00 | 1.57 | 0.53 | 0.19 | 0 | Glutamatergic Neuron |
| ***KCNC2*** | 5.18E-290 | 9.86E-286 | 1.54 | 0.40 | 0.15 | 0 | Glutamatergic Neuron |
| ***MGAT4C*** | 0.00E+00 | 0.00E+00 | 1.52 | 0.59 | 0.21 | 0 | Glutamatergic Neuron |
| ***GRIK3*** | 0.00E+00 | 0.00E+00 | 1.49 | 0.46 | 0.17 | 0 | Glutamatergic Neuron |
| ***NRP2*** | 1.80E-305 | 3.42E-301 | 1.46 | 0.42 | 0.16 | 0 | Glutamatergic Neuron |
| ***ZFHX3*** | 3.41E-299 | 6.49E-295 | 1.45 | 0.41 | 0.15 | 0 | Glutamatergic Neuron |
| ***LOC100229124*** | 0.00E+00 | 0.00E+00 | 1.42 | 0.47 | 0.18 | 0 | Glutamatergic Neuron |
| ***SHISA9*** | 0.00E+00 | 0.00E+00 | 1.41 | 0.62 | 0.28 | 0 | Glutamatergic Neuron |
| ***LOC100219930*** | 2.10E-283 | 4.00E-279 | 1.38 | 0.47 | 0.20 | 0 | Glutamatergic Neuron |
| ***GPR158*** | 2.52E-295 | 4.79E-291 | 1.38 | 0.46 | 0.19 | 0 | Glutamatergic Neuron |
| ***UNC5D*** | 8.47E-298 | 1.61E-293 | 1.37 | 0.48 | 0.21 | 0 | Glutamatergic Neuron |
| ***CDH12*** | 6.21E-311 | 1.18E-306 | 1.35 | 0.55 | 0.26 | 0 | Glutamatergic Neuron |
| ***UNC80*** | 0.00E+00 | 0.00E+00 | 1.35 | 0.63 | 0.25 | 0 | Glutamatergic Neuron |
| ***ERC2*** | 0.00E+00 | 0.00E+00 | 1.32 | 0.82 | 0.38 | 0 | Glutamatergic Neuron |
| ***SLITRK1*** | 1.36E-303 | 2.59E-299 | 1.31 | 0.49 | 0.21 | 0 | Glutamatergic Neuron |
| ***RYR2*** | 0.00E+00 | 0.00E+00 | 1.31 | 0.57 | 0.25 | 0 | Glutamatergic Neuron |

| Gene | P-Value | FDR | Average $log_2FC$ | Fraction 1 | Fraction 2 | Cluster | Annotation |
|---|---|---|---|---|---|---|---|
| *LOC105758645* | 0.00E+00 | 0.00E+00 | 2.89 | 0.91 | 0.26 | 1 | Astrocyte |
| *FAM53A* | 0.00E+00 | 0.00E+00 | 2.74 | 0.68 | 0.21 | 1 | Astrocyte |
| *KCNQ5* | 0.00E+00 | 0.00E+00 | 2.28 | 0.62 | 0.19 | 1 | Astrocyte |
| *GLCCI1* | 0.00E+00 | 0.00E+00 | 2.27 | 0.67 | 0.28 | 1 | Astrocyte |
| *NOTCH1* | 0.00E+00 | 0.00E+00 | 2.20 | 0.53 | 0.17 | 1 | Astrocyte |
| *BCAR3* | 0.00E+00 | 0.00E+00 | 2.14 | 0.43 | 0.14 | 1 | Astrocyte |
| *LRP5* | 0.00E+00 | 0.00E+00 | 2.06 | 0.54 | 0.19 | 1 | Astrocyte |
| *IGDCC3* | 0.00E+00 | 0.00E+00 | 2.00 | 0.39 | 0.14 | 1 | Astrocyte |
| *GLI3* | 0.00E+00 | 0.00E+00 | 1.91 | 0.53 | 0.16 | 1 | Astrocyte |
| *PRDM16* | 0.00E+00 | 0.00E+00 | 1.81 | 0.66 | 0.22 | 1 | Astrocyte |
| *FOXG1* | 0.00E+00 | 0.00E+00 | 1.81 | 0.86 | 0.42 | 1 | Astrocyte |
| *MLC1* | 2.73E-302 | 5.19E-298 | 1.73 | 0.42 | 0.15 | 1 | Astrocyte |
| *FGFRL1* | 0.00E+00 | 0.00E+00 | 1.59 | 0.59 | 0.19 | 1 | Astrocyte |
| *TMEM131* | 0.00E+00 | 0.00E+00 | 1.54 | 0.65 | 0.40 | 1 | Astrocyte |
| *TTYH3* | 0.00E+00 | 0.00E+00 | 1.53 | 0.57 | 0.31 | 1 | Astrocyte |
| *LOC105758648* | 0.00E+00 | 0.00E+00 | 1.52 | 0.72 | 0.34 | 1 | Astrocyte |
| *CREB5* | 5.52E-285 | 1.05E-280 | 1.51 | 0.47 | 0.19 | 1 | Astrocyte |
| *PLXNB2* | 0.00E+00 | 0.00E+00 | 1.43 | 0.80 | 0.52 | 1 | Astrocyte |
| *LRIG1* | 0.00E+00 | 0.00E+00 | 1.42 | 0.66 | 0.29 | 1 | Astrocyte |
| *SOX2* | 0.00E+00 | 0.00E+00 | 1.29 | 0.99 | 0.68 | 1 | Astrocyte |
| *EBF1* | 0.00E+00 | 0.00E+00 | 5.33 | 0.90 | 0.07 | 10 | GABAergic Neuron |
| *ST8SIA5* | 0.00E+00 | 0.00E+00 | 3.12 | 0.40 | 0.07 | 10 | GABAergic Neuron |
| *VCAN* | 0.00E+00 | 0.00E+00 | 2.72 | 0.88 | 0.36 | 10 | GABAergic Neuron |
| *BCL11B* | 0.00E+00 | 0.00E+00 | 2.59 | 0.97 | 0.39 | 10 | GABAergic Neuron |
| *MEIS2* | 0.00E+00 | 0.00E+00 | 2.46 | 0.94 | 0.29 | 10 | GABAergic Neuron |
| *FOXP1* | 0.00E+00 | 0.00E+00 | 2.40 | 0.66 | 0.28 | 10 | GABAergic Neuron |
| *OXR1* | 0.00E+00 | 0.00E+00 | 2.38 | 0.70 | 0.33 | 10 | GABAergic Neuron |

| Gene | P-Value | FDR | Average $log_2FC$ | Fraction 1 | Fraction 2 | Cluster | Annotation |
|---|---|---|---|---|---|---|---|
| *NKAIN3* | 0.00E+00 | 0.00E+00 | 2.21 | 0.75 | 0.35 | 10 | GABAergic Neuron |
| *FOXP2* | 0.00E+00 | 0.00E+00 | 2.14 | 0.79 | 0.37 | 10 | GABAergic Neuron |
| *EPHA4* | 0.00E+00 | 0.00E+00 | 2.12 | 0.57 | 0.19 | 10 | GABAergic Neuron |
| *ADCY5* | 0.00E+00 | 0.00E+00 | 2.12 | 0.60 | 0.25 | 10 | GABAergic Neuron |
| *RAPGEF5* | 0.00E+00 | 0.00E+00 | 2.04 | 0.62 | 0.23 | 10 | GABAergic Neuron |
| *ISL1* | 1.42E-267 | 2.69E-263 | 1.94 | 0.46 | 0.13 | 10 | GABAergic Neuron |
| *ARPP21* | 0.00E+00 | 0.00E+00 | 1.93 | 0.92 | 0.52 | 10 | GABAergic Neuron |
| *LOC115494452* | 0.00E+00 | 0.00E+00 | 1.85 | 0.85 | 0.41 | 10 | GABAergic Neuron |
| *BCL11A* | 0.00E+00 | 0.00E+00 | 1.84 | 0.81 | 0.34 | 10 | GABAergic Neuron |
| *TSPAN9* | 6.08E-259 | 1.16E-254 | 1.82 | 0.51 | 0.20 | 10 | GABAergic Neuron |
| *PHACTR1* | 0.00E+00 | 0.00E+00 | 1.81 | 0.87 | 0.48 | 10 | GABAergic Neuron |
| *HS6ST1* | 0.00E+00 | 0.00E+00 | 1.66 | 0.67 | 0.36 | 10 | GABAergic Neuron |
| *KIF26A* | 2.72E-181 | 5.17E-177 | 1.66 | 0.43 | 0.16 | 10 | GABAergic Neuron |
| *LOC105758645* | 0.00E+00 | 0.00E+00 | 3.32 | 0.97 | 0.27 | 11 | Astrocyte |
| *GLI3* | 0.00E+00 | 0.00E+00 | 3.01 | 0.79 | 0.16 | 11 | Astrocyte |
| *GLI2* | 0.00E+00 | 0.00E+00 | 2.97 | 0.49 | 0.09 | 11 | Astrocyte |
| *HMGA2* | 2.52E-242 | 4.80E-238 | 2.62 | 0.32 | 0.07 | 11 | Astrocyte |
| *KCNH8* | 0.00E+00 | 0.00E+00 | 2.41 | 0.52 | 0.14 | 11 | Astrocyte |
| *KIAA1217* | 0.00E+00 | 0.00E+00 | 2.41 | 0.48 | 0.12 | 11 | Astrocyte |
| *LOC115493838* | 0.00E+00 | 0.00E+00 | 2.33 | 0.56 | 0.19 | 11 | Astrocyte |
| *KCNQ5* | 0.00E+00 | 0.00E+00 | 2.30 | 0.70 | 0.20 | 11 | Astrocyte |
| *PLXNB2* | 0.00E+00 | 0.00E+00 | 2.23 | 0.92 | 0.53 | 11 | Astrocyte |
| *PRDM16* | 0.00E+00 | 0.00E+00 | 2.16 | 0.76 | 0.23 | 11 | Astrocyte |
| *MID1* | 0.00E+00 | 0.00E+00 | 2.15 | 0.72 | 0.26 | 11 | Astrocyte |
| *LOC100223646* | 0.00E+00 | 0.00E+00 | 2.12 | 0.77 | 0.37 | 11 | Astrocyte |
| *CDK6* | 5.46E-321 | 1.04E-316 | 2.06 | 0.51 | 0.19 | 11 | Astrocyte |
| *FAM53A* | 6.70E-286 | 1.27E-281 | 1.90 | 0.56 | 0.23 | 11 | Astrocyte |

| Gene | P-Value | FDR | Average $log_2FC$ | Fraction 1 | Fraction 2 | Cluster | Annotation |
|---|---|---|---|---|---|---|---|
| ***FGFRL1*** | 0.00E+00 | 0.00E+00 | 1.82 | 0.69 | 0.20 | 11 | Astrocyte |
| ***FOXG1*** | 0.00E+00 | 0.00E+00 | 1.70 | 0.87 | 0.43 | 11 | Astrocyte |
| ***NOTCH1*** | 1.39E-197 | 2.64E-193 | 1.70 | 0.47 | 0.18 | 11 | Astrocyte |
| ***CREB5*** | 5.39E-245 | 1.03E-240 | 1.67 | 0.55 | 0.20 | 11 | Astrocyte |
| ***SEMA5B*** | 3.02E-243 | 5.74E-239 | 1.62 | 0.57 | 0.25 | 11 | Astrocyte |
| ***LRP5*** | 2.62E-175 | 4.98E-171 | 1.55 | 0.48 | 0.20 | 11 | Astrocyte |
| ***LHX8*** | 0.00E+00 | 0.00E+00 | 4.43 | 0.49 | 0.03 | 12 | Glutamatergic Neuron |
| ***ADARB2*** | 0.00E+00 | 0.00E+00 | 3.39 | 0.73 | 0.23 | 12 | Glutamatergic Neuron |
| ***LOC121469413*** | 9.29E-275 | 1.77E-270 | 3.37 | 0.30 | 0.04 | 12 | Glutamatergic Neuron |
| ***SNTG2*** | 1.13E-202 | 2.15E-198 | 2.65 | 0.34 | 0.07 | 12 | Glutamatergic Neuron |
| ***LHX6*** | 3.77E-246 | 7.17E-242 | 2.31 | 0.34 | 0.06 | 12 | Glutamatergic Neuron |
| ***GABRG3*** | 0.00E+00 | 0.00E+00 | 2.17 | 0.71 | 0.23 | 12 | Glutamatergic Neuron |
| ***CACNA1I*** | 6.18E-172 | 1.17E-167 | 2.15 | 0.36 | 0.10 | 12 | Glutamatergic Neuron |
| ***KLHL1*** | 1.86E-172 | 3.54E-168 | 1.92 | 0.44 | 0.15 | 12 | Glutamatergic Neuron |
| ***LOC121469249*** | 6.13E-186 | 1.17E-181 | 1.69 | 0.50 | 0.17 | 12 | Glutamatergic Neuron |
| ***VSTM2A*** | 9.31E-143 | 1.77E-138 | 1.62 | 0.44 | 0.17 | 12 | Glutamatergic Neuron |
| ***CNTN5*** | 1.09E-168 | 2.08E-164 | 1.59 | 0.59 | 0.27 | 12 | Glutamatergic Neuron |
| ***RYR2*** | 6.31E-229 | 1.20E-224 | 1.57 | 0.65 | 0.26 | 12 | Glutamatergic Neuron |
| ***ZNF385D*** | 1.16E-290 | 2.21E-286 | 1.43 | 0.84 | 0.45 | 12 | Glutamatergic Neuron |
| ***SLITRK1*** | 3.17E-150 | 6.03E-146 | 1.43 | 0.52 | 0.22 | 12 | Glutamatergic Neuron |
| ***GABRA1*** | 1.16E-123 | 2.21E-119 | 1.41 | 0.44 | 0.17 | 12 | Glutamatergic Neuron |
| ***RGS7*** | 3.13E-210 | 5.95E-206 | 1.35 | 0.70 | 0.32 | 12 | Glutamatergic Neuron |
| ***EPB41L3*** | 1.15E-141 | 2.18E-137 | 1.32 | 0.55 | 0.25 | 12 | Glutamatergic Neuron |
| ***NCAM2*** | 7.24E-281 | 1.38E-276 | 1.32 | 0.89 | 0.55 | 12 | Glutamatergic Neuron |
| ***LOC100219930*** | 1.08E-114 | 2.06E-110 | 1.29 | 0.48 | 0.22 | 12 | Glutamatergic Neuron |
| ***TSHZ2*** | 1.11E-117 | 2.12E-113 | 1.27 | 0.54 | 0.27 | 12 | Glutamatergic Neuron |
| ***TMEM272*** | 6.51E-291 | 1.24E-286 | 3.48 | 0.31 | 0.03 | 13 | GABAergic Neuron |

| Gene | P-Value | FDR | Average $log_2FC$ | Fraction 1 | Fraction 2 | Cluster | Annotation |
|---|---|---|---|---|---|---|---|
| ***KCNH8*** | 0.00E+00 | 0.00E+00 | 3.22 | 0.73 | 0.14 | 13 | GABAergic Neuron |
| ***PENK*** | 0.00E+00 | 0.00E+00 | 2.82 | 0.62 | 0.25 | 13 | GABAergic Neuron |
| ***SMOC2*** | 3.42E-246 | 6.50E-242 | 2.69 | 0.38 | 0.10 | 13 | GABAergic Neuron |
| ***MEIS2*** | 0.00E+00 | 0.00E+00 | 2.64 | 0.92 | 0.30 | 13 | GABAergic Neuron |
| ***SEMA3A*** | 5.56E-246 | 1.06E-241 | 2.61 | 0.40 | 0.09 | 13 | GABAergic Neuron |
| ***CACNA2D3*** | 0.00E+00 | 0.00E+00 | 2.50 | 0.79 | 0.23 | 13 | GABAergic Neuron |
| ***LOC115494452*** | 0.00E+00 | 0.00E+00 | 2.45 | 0.94 | 0.41 | 13 | GABAergic Neuron |
| ***RUNX1T1*** | 0.00E+00 | 0.00E+00 | 2.27 | 0.86 | 0.36 | 13 | GABAergic Neuron |
| ***BCL11B*** | 0.00E+00 | 0.00E+00 | 2.20 | 0.93 | 0.40 | 13 | GABAergic Neuron |
| ***LOC121469248*** | 1.70E-198 | 3.23E-194 | 2.14 | 0.41 | 0.12 | 13 | GABAergic Neuron |
| ***AFF3*** | 3.38E-259 | 6.43E-255 | 2.10 | 0.53 | 0.19 | 13 | GABAergic Neuron |
| ***ZNF804A*** | 7.24E-222 | 1.38E-217 | 1.95 | 0.55 | 0.25 | 13 | GABAergic Neuron |
| ***PHACTR1*** | 0.00E+00 | 0.00E+00 | 1.70 | 0.81 | 0.48 | 13 | GABAergic Neuron |
| ***LOC116808653*** | 3.20E-311 | 6.09E-307 | 1.48 | 0.81 | 0.45 | 13 | GABAergic Neuron |
| ***PBX3*** | 6.83E-222 | 1.30E-217 | 1.47 | 0.68 | 0.40 | 13 | GABAergic Neuron |
| ***EPHA4*** | 2.14E-115 | 4.06E-111 | 1.44 | 0.45 | 0.20 | 13 | GABAergic Neuron |
| ***MYT1L*** | 0.00E+00 | 0.00E+00 | 1.26 | 0.97 | 0.64 | 13 | GABAergic Neuron |
| ***BCL11A*** | 9.24E-124 | 1.76E-119 | 1.19 | 0.62 | 0.35 | 13 | GABAergic Neuron |
| ***RASGEF1A*** | 4.41E-140 | 8.38E-136 | 1.16 | 0.68 | 0.41 | 13 | GABAergic Neuron |
| ***PROX1*** | 0.00E+00 | 0.00E+00 | 2.75 | 0.53 | 0.12 | 14 | Glutamatergic Neuron |
| ***ISL1*** | 9.38E-173 | 1.78E-168 | 1.93 | 0.43 | 0.14 | 14 | Glutamatergic Neuron |
| ***SORCS1*** | 1.13E-162 | 2.15E-158 | 1.66 | 0.54 | 0.26 | 14 | Glutamatergic Neuron |
| ***LRFN2*** | 4.79E-128 | 9.10E-124 | 1.56 | 0.47 | 0.20 | 14 | Glutamatergic Neuron |
| ***CACNA2D3*** | 1.55E-158 | 2.94E-154 | 1.53 | 0.56 | 0.24 | 14 | Glutamatergic Neuron |
| ***LOC101234143*** | 3.12E-121 | 5.94E-117 | 1.39 | 0.45 | 0.18 | 14 | Glutamatergic Neuron |
| ***LRP1B*** | 3.99E-221 | 7.58E-217 | 1.36 | 0.75 | 0.35 | 14 | Glutamatergic Neuron |
| ***LINGO2*** | 1.97E-205 | 3.74E-201 | 1.32 | 0.76 | 0.38 | 14 | Glutamatergic Neuron |

| Gene | P-Value | FDR | Average $log_2FC$ | Fraction 1 | Fraction 2 | Cluster | Annotation |
|---|---|---|---|---|---|---|---|
| ***LOC115491243*** | 2.03E-264 | 3.85E-260 | 1.31 | 0.91 | 0.58 | 14 | Glutamatergic Neuron |
| ***PCDH7*** | 1.40E-161 | 2.66E-157 | 1.28 | 0.76 | 0.47 | 14 | Glutamatergic Neuron |
| ***CDH18*** | 1.48E-109 | 2.81E-105 | 1.24 | 0.46 | 0.20 | 14 | Glutamatergic Neuron |
| ***LOC100224567*** | 9.74E-220 | 1.85E-215 | 1.22 | 0.79 | 0.40 | 14 | Glutamatergic Neuron |
| ***LRRTM3*** | 9.13E-96 | 1.74E-91 | 1.18 | 0.51 | 0.26 | 14 | Glutamatergic Neuron |
| ***DPP6*** | 3.86E-197 | 7.34E-193 | 1.17 | 0.88 | 0.54 | 14 | Glutamatergic Neuron |
| ***EPB41L3*** | 9.76E-101 | 1.86E-96 | 1.13 | 0.51 | 0.25 | 14 | Glutamatergic Neuron |
| ***FAM135B*** | 1.26E-107 | 2.40E-103 | 1.13 | 0.51 | 0.24 | 14 | Glutamatergic Neuron |
| ***KCNB2*** | 2.33E-97 | 4.43E-93 | 1.11 | 0.56 | 0.30 | 14 | Glutamatergic Neuron |
| ***RASGEF1A*** | 8.86E-144 | 1.68E-139 | 1.11 | 0.73 | 0.41 | 14 | Glutamatergic Neuron |
| ***GRM5*** | 8.72E-186 | 1.66E-181 | 1.10 | 0.71 | 0.33 | 14 | Glutamatergic Neuron |
| ***RGS7*** | 4.14E-139 | 7.87E-135 | 1.10 | 0.65 | 0.33 | 14 | Glutamatergic Neuron |
| ***PDGFRA*** | 0.00E+00 | 0.00E+00 | 6.32 | 0.93 | 0.02 | 15 | Oligodendrocyte |
| ***CA8*** | 0.00E+00 | 0.00E+00 | 6.10 | 0.80 | 0.02 | 15 | Oligodendrocyte |
| ***LOC121468099*** | 0.00E+00 | 0.00E+00 | 5.63 | 0.26 | 0.01 | 15 | Oligodendrocyte |
| ***SPRY4*** | 0.00E+00 | 0.00E+00 | 4.91 | 0.31 | 0.01 | 15 | Oligodendrocyte |
| ***COLEC12*** | 0.00E+00 | 0.00E+00 | 4.87 | 0.83 | 0.05 | 15 | Oligodendrocyte |
| ***ZCCHC24*** | 0.00E+00 | 0.00E+00 | 4.25 | 0.93 | 0.12 | 15 | Oligodendrocyte |
| ***NKX2-2*** | 4.10E-306 | 7.79E-302 | 3.82 | 0.29 | 0.02 | 15 | Oligodendrocyte |
| ***GAB3*** | 0.00E+00 | 0.00E+00 | 3.75 | 0.34 | 0.03 | 15 | Oligodendrocyte |
| ***NCAN*** | 0.00E+00 | 0.00E+00 | 3.69 | 0.68 | 0.09 | 15 | Oligodendrocyte |
| ***REPS2*** | 0.00E+00 | 0.00E+00 | 3.49 | 0.61 | 0.08 | 15 | Oligodendrocyte |
| ***LOC121470875*** | 0.00E+00 | 0.00E+00 | 3.42 | 0.44 | 0.05 | 15 | Oligodendrocyte |
| ***CNTFR*** | 0.00E+00 | 0.00E+00 | 3.34 | 0.78 | 0.15 | 15 | Oligodendrocyte |
| ***ARHGEF3*** | 0.00E+00 | 0.00E+00 | 3.25 | 0.55 | 0.07 | 15 | Oligodendrocyte |
| ***SH3RF1*** | 0.00E+00 | 0.00E+00 | 3.17 | 0.82 | 0.19 | 15 | Oligodendrocyte |
| ***PIEZO2*** | 0.00E+00 | 0.00E+00 | 3.10 | 0.52 | 0.06 | 15 | Oligodendrocyte |

| Gene | P-Value | FDR | Average $log_2FC$ | Fraction 1 | Fraction 2 | Cluster | Annotation |
|---|---|---|---|---|---|---|---|
| ***SOX8*** | 3.62E-283 | 6.89E-279 | 3.08 | 0.35 | 0.04 | 15 | Oligodendrocyte |
| ***SERPINE2*** | 0.00E+00 | 0.00E+00 | 3.03 | 0.64 | 0.12 | 15 | Oligodendrocyte |
| ***STARD9*** | 0.00E+00 | 0.00E+00 | 3.03 | 0.83 | 0.19 | 15 | Oligodendrocyte |
| ***SPRY2*** | 0.00E+00 | 0.00E+00 | 2.97 | 0.60 | 0.09 | 15 | Oligodendrocyte |
| ***TUB*** | 0.00E+00 | 0.00E+00 | 2.93 | 0.90 | 0.27 | 15 | Oligodendrocyte |
| ***SIM1*** | 0.00E+00 | 0.00E+00 | 5.42 | 0.47 | 0.02 | 16 | Glutamatergic Neuron |
| ***EBF3*** | 0.00E+00 | 0.00E+00 | 4.62 | 0.67 | 0.04 | 16 | Glutamatergic Neuron |
| ***LOC100231196*** | 0.00E+00 | 0.00E+00 | 3.68 | 0.40 | 0.04 | 16 | Glutamatergic Neuron |
| ***NRN1*** | 0.00E+00 | 0.00E+00 | 3.00 | 0.45 | 0.06 | 16 | Glutamatergic Neuron |
| ***NHLH2*** | 4.63E-302 | 8.80E-298 | 2.92 | 0.40 | 0.06 | 16 | Glutamatergic Neuron |
| ***LOC100221630*** | 0.00E+00 | 0.00E+00 | 2.76 | 0.65 | 0.17 | 16 | Glutamatergic Neuron |
| ***SLC17A6*** | 1.66E-301 | 3.16E-297 | 2.53 | 0.48 | 0.09 | 16 | Glutamatergic Neuron |
| ***LOC100226371*** | 1.04E-272 | 1.97E-268 | 2.40 | 0.48 | 0.10 | 16 | Glutamatergic Neuron |
| ***HS6ST3*** | 2.85E-176 | 5.42E-172 | 2.37 | 0.41 | 0.13 | 16 | Glutamatergic Neuron |
| ***CDH18*** | 0.00E+00 | 0.00E+00 | 2.36 | 0.72 | 0.19 | 16 | Glutamatergic Neuron |
| ***CACNA2D1*** | 0.00E+00 | 0.00E+00 | 2.28 | 0.92 | 0.42 | 16 | Glutamatergic Neuron |
| ***TMEM163*** | 5.51E-183 | 1.05E-178 | 2.13 | 0.39 | 0.10 | 16 | Glutamatergic Neuron |
| ***CPNE4*** | 1.98E-144 | 3.76E-140 | 2.10 | 0.39 | 0.13 | 16 | Glutamatergic Neuron |
| ***SLIT3*** | 1.87E-270 | 3.56E-266 | 2.10 | 0.62 | 0.23 | 16 | Glutamatergic Neuron |
| ***ABCC5*** | 1.39E-247 | 2.64E-243 | 1.99 | 0.53 | 0.24 | 16 | Glutamatergic Neuron |
| ***TAFA1*** | 1.57E-199 | 2.99E-195 | 1.95 | 0.49 | 0.15 | 16 | Glutamatergic Neuron |
| ***GALNT9*** | 3.08E-152 | 5.85E-148 | 1.78 | 0.43 | 0.14 | 16 | Glutamatergic Neuron |
| ***SCG2*** | 4.80E-171 | 9.12E-167 | 1.76 | 0.54 | 0.22 | 16 | Glutamatergic Neuron |
| ***LRP1B*** | 1.12E-317 | 2.13E-313 | 1.72 | 0.82 | 0.35 | 16 | Glutamatergic Neuron |
| ***KLHL1*** | 5.06E-122 | 9.63E-118 | 1.72 | 0.41 | 0.15 | 16 | Glutamatergic Neuron |
| ***LOC121470376*** | 0.00E+00 | 0.00E+00 | 3.43 | 0.97 | 0.25 | 17 | GABAergic Neuron |
| ***LOC121470375*** | 0.00E+00 | 0.00E+00 | 3.39 | 0.81 | 0.15 | 17 | GABAergic Neuron |

| Gene | P-Value | FDR | Average $log_2FC$ | Fraction 1 | Fraction 2 | Cluster | Annotation |
|---|---|---|---|---|---|---|---|
| *EPHA3* | 0.00E+00 | 0.00E+00 | 2.90 | 0.90 | 0.34 | 17 | GABAergic Neuron |
| *MEIS2* | 0.00E+00 | 0.00E+00 | 2.62 | 0.93 | 0.30 | 17 | GABAergic Neuron |
| *SHB* | 2.30E-273 | 4.37E-269 | 2.62 | 0.35 | 0.10 | 17 | GABAergic Neuron |
| *FOXP2* | 0.00E+00 | 0.00E+00 | 2.49 | 0.77 | 0.37 | 17 | GABAergic Neuron |
| *LOC115496358* | 0.00E+00 | 0.00E+00 | 2.36 | 0.72 | 0.25 | 17 | GABAergic Neuron |
| *DPF3* | 2.53E-314 | 4.82E-310 | 2.26 | 0.49 | 0.18 | 17 | GABAergic Neuron |
| *PLXNA2* | 0.00E+00 | 0.00E+00 | 2.25 | 0.66 | 0.26 | 17 | GABAergic Neuron |
| *ESRRG* | 0.00E+00 | 0.00E+00 | 2.24 | 0.57 | 0.32 | 17 | GABAergic Neuron |
| *LOC116808643* | 0.00E+00 | 0.00E+00 | 2.22 | 0.62 | 0.27 | 17 | GABAergic Neuron |
| *MYT1* | 0.00E+00 | 0.00E+00 | 2.20 | 0.68 | 0.27 | 17 | GABAergic Neuron |
| *SH3RF3* | 0.00E+00 | 0.00E+00 | 2.14 | 0.93 | 0.57 | 17 | GABAergic Neuron |
| *MN1* | 3.17E-252 | 6.03E-248 | 1.97 | 0.50 | 0.20 | 17 | GABAergic Neuron |
| *SLAIN1* | 6.43E-287 | 1.22E-282 | 1.96 | 0.63 | 0.29 | 17 | GABAergic Neuron |
| *ST18* | 7.25E-134 | 1.38E-129 | 1.92 | 0.39 | 0.14 | 17 | GABAergic Neuron |
| *LOC100227840* | 1.91E-168 | 3.63E-164 | 1.90 | 0.47 | 0.21 | 17 | GABAergic Neuron |
| *ARX* | 1.15E-143 | 2.18E-139 | 1.64 | 0.49 | 0.21 | 17 | GABAergic Neuron |
| *PLXNB2* | 0.00E+00 | 0.00E+00 | 1.53 | 0.81 | 0.53 | 17 | GABAergic Neuron |
| *SOX4* | 6.83E-180 | 1.30E-175 | 1.43 | 0.62 | 0.33 | 17 | GABAergic Neuron |
| *CXCL12* | 0.00E+00 | 0.00E+00 | 4.38 | 0.49 | 0.04 | 18 | GABAergic Neuron |
| *ETV1* | 0.00E+00 | 0.00E+00 | 3.47 | 0.63 | 0.10 | 18 | GABAergic Neuron |
| *LHX6* | 1.39E-232 | 2.64E-228 | 2.82 | 0.37 | 0.06 | 18 | GABAergic Neuron |
| *OPRM1* | 0.00E+00 | 0.00E+00 | 2.67 | 0.65 | 0.17 | 18 | GABAergic Neuron |
| *RASA3* | 0.00E+00 | 0.00E+00 | 2.58 | 0.64 | 0.20 | 18 | GABAergic Neuron |
| *ST18* | 0.00E+00 | 0.00E+00 | 2.58 | 0.61 | 0.13 | 18 | GABAergic Neuron |
| *ARX* | 0.00E+00 | 0.00E+00 | 2.38 | 0.76 | 0.21 | 18 | GABAergic Neuron |
| *NRXN3* | 0.00E+00 | 0.00E+00 | 2.29 | 1.00 | 0.70 | 18 | GABAergic Neuron |
| *OSBPL5* | 5.00E-324 | 9.39E-320 | 2.22 | 0.53 | 0.20 | 18 | GABAergic Neuron |

| Gene | P-Value | FDR | Average $log_2FC$ | Fraction 1 | Fraction 2 | Cluster | Annotation |
|---|---|---|---|---|---|---|---|
| ***SLAIN1*** | 6.87E-322 | 1.31E-317 | 2.00 | 0.71 | 0.28 | 18 | GABAergic Neuron |
| ***ZNF536*** | 0.00E+00 | 0.00E+00 | 1.94 | 0.92 | 0.47 | 18 | GABAergic Neuron |
| ***DNAAF9*** | 0.00E+00 | 0.00E+00 | 1.85 | 0.95 | 0.64 | 18 | GABAergic Neuron |
| ***NFIB*** | 0.00E+00 | 0.00E+00 | 1.78 | 0.86 | 0.43 | 18 | GABAergic Neuron |
| ***NYAP2*** | 5.65E-280 | 1.07E-275 | 1.77 | 0.65 | 0.31 | 18 | GABAergic Neuron |
| ***ROBO1*** | 0.00E+00 | 0.00E+00 | 1.74 | 1.00 | 0.75 | 18 | GABAergic Neuron |
| ***KIAA1549*** | 4.32E-312 | 8.21E-308 | 1.57 | 0.71 | 0.45 | 18 | GABAergic Neuron |
| ***DACH2*** | 9.82E-115 | 1.87E-110 | 1.46 | 0.49 | 0.21 | 18 | GABAergic Neuron |
| ***GAD2*** | 6.07E-191 | 1.15E-186 | 1.42 | 0.71 | 0.34 | 18 | GABAergic Neuron |
| ***SOX4*** | 1.66E-148 | 3.15E-144 | 1.35 | 0.63 | 0.33 | 18 | GABAergic Neuron |
| ***BCL11A*** | 1.16E-132 | 2.21E-128 | 1.23 | 0.65 | 0.35 | 18 | GABAergic Neuron |
| ***COL19A1*** | 0.00E+00 | 0.00E+00 | 3.77 | 0.48 | 0.06 | 19 | Glutamatergic/GABA |
| ***LOC115496226*** | 4.11E-319 | 7.81E-315 | 3.70 | 0.35 | 0.05 | 19 | Glutamatergic/GABA |
| ***ELFN1*** | 0.00E+00 | 0.00E+00 | 3.37 | 0.73 | 0.16 | 19 | Glutamatergic/GABA |
| ***LHX8*** | 0.00E+00 | 0.00E+00 | 3.37 | 0.38 | 0.03 | 19 | Glutamatergic/GABA |
| ***INSYN2A*** | 0.00E+00 | 0.00E+00 | 2.82 | 0.50 | 0.10 | 19 | Glutamatergic/GABA |
| ***SMOC2*** | 1.58E-224 | 3.00E-220 | 2.81 | 0.40 | 0.10 | 19 | Glutamatergic/GABA |
| ***LHX6*** | 6.91E-312 | 1.31E-307 | 2.80 | 0.43 | 0.06 | 19 | Glutamatergic/GABA |
| ***ADARB2*** | 0.00E+00 | 0.00E+00 | 2.76 | 0.78 | 0.23 | 19 | Glutamatergic/GABA |
| ***SNTG2*** | 2.68E-174 | 5.10E-170 | 2.75 | 0.35 | 0.08 | 19 | Glutamatergic/GABA |
| ***LOC121469604*** | 1.69E-183 | 3.21E-179 | 2.70 | 0.31 | 0.05 | 19 | Glutamatergic/GABA |
| ***SRRM4*** | 0.00E+00 | 0.00E+00 | 2.54 | 0.61 | 0.15 | 19 | Glutamatergic/GABA |
| ***MEF2C*** | 6.74E-314 | 1.28E-309 | 2.53 | 0.57 | 0.14 | 19 | Glutamatergic/GABA |
| ***GABRA1*** | 0.00E+00 | 0.00E+00 | 2.49 | 0.60 | 0.17 | 19 | Glutamatergic/GABA |
| ***PDE3A*** | 0.00E+00 | 0.00E+00 | 2.44 | 0.73 | 0.26 | 19 | Glutamatergic/GABA |
| ***SULF2*** | 0.00E+00 | 0.00E+00 | 2.38 | 0.65 | 0.25 | 19 | Glutamatergic/GABA |
| ***VSTM2A*** | 1.57E-298 | 2.98E-294 | 2.37 | 0.58 | 0.17 | 19 | Glutamatergic/GABA |

| Gene | P-Value | FDR | Average $log_2FC$ | Fraction 1 | Fraction 2 | Cluster | Annotation |
|---|---|---|---|---|---|---|---|
| *VWC2* | 2.26E-141 | 4.29E-137 | 2.23 | 0.33 | 0.08 | 19 | Glutamatergic/GABA |
| *PTPN5* | 4.78E-272 | 9.09E-268 | 2.13 | 0.56 | 0.17 | 19 | Glutamatergic/GABA |
| *C2H8orf34* | 5.00E-323 | 9.39E-319 | 2.10 | 0.68 | 0.26 | 19 | Glutamatergic/GABA |
| *GRM1* | 6.97E-163 | 1.32E-158 | 2.08 | 0.42 | 0.16 | 19 | Glutamatergic/GABA |
| *PTPRZ1* | 0.00E+00 | 0.00E+00 | 3.74 | 1.00 | 0.46 | 2 | Astrocyte |
| *GPC5* | 0.00E+00 | 0.00E+00 | 3.58 | 0.80 | 0.10 | 2 | Astrocyte |
| *SLC4A4* | 0.00E+00 | 0.00E+00 | 3.45 | 0.80 | 0.13 | 2 | Astrocyte |
| *ADAM12* | 0.00E+00 | 0.00E+00 | 3.36 | 0.34 | 0.04 | 2 | Astrocyte |
| *LOC115495208* | 0.00E+00 | 0.00E+00 | 3.25 | 0.35 | 0.04 | 2 | Astrocyte |
| *VIT* | 0.00E+00 | 0.00E+00 | 3.08 | 0.31 | 0.04 | 2 | Astrocyte |
| *SLC1A3* | 0.00E+00 | 0.00E+00 | 2.93 | 0.83 | 0.18 | 2 | Astrocyte |
| *PROCA1* | 0.00E+00 | 0.00E+00 | 2.89 | 0.30 | 0.05 | 2 | Astrocyte |
| *RFX4* | 0.00E+00 | 0.00E+00 | 2.84 | 0.63 | 0.10 | 2 | Astrocyte |
| *FABP7* | 0.00E+00 | 0.00E+00 | 2.84 | 0.93 | 0.38 | 2 | Astrocyte |
| *THRB* | 0.00E+00 | 0.00E+00 | 2.82 | 0.46 | 0.08 | 2 | Astrocyte |
| *TNS1* | 0.00E+00 | 0.00E+00 | 2.73 | 0.30 | 0.05 | 2 | Astrocyte |
| *BCAN* | 0.00E+00 | 0.00E+00 | 2.71 | 0.40 | 0.07 | 2 | Astrocyte |
| *EGF* | 0.00E+00 | 0.00E+00 | 2.70 | 0.86 | 0.18 | 2 | Astrocyte |
| *SLC1A2* | 0.00E+00 | 0.00E+00 | 2.69 | 0.75 | 0.22 | 2 | Astrocyte |
| *PITPNC1* | 0.00E+00 | 0.00E+00 | 2.68 | 0.50 | 0.11 | 2 | Astrocyte |
| *SMOC1* | 0.00E+00 | 0.00E+00 | 2.65 | 0.71 | 0.16 | 2 | Astrocyte |
| *SLC6A11* | 0.00E+00 | 0.00E+00 | 2.65 | 0.57 | 0.16 | 2 | Astrocyte |
| *FGFR3* | 0.00E+00 | 0.00E+00 | 2.64 | 0.76 | 0.17 | 2 | Astrocyte |
| *GRM3* | 0.00E+00 | 0.00E+00 | 2.64 | 0.63 | 0.17 | 2 | Astrocyte |
| *CRACR2A* | 0.00E+00 | 0.00E+00 | 5.02 | 0.39 | 0.02 | 20 | Astrocyte |
| *MOXD1* | 5.00E-324 | 9.39E-320 | 4.24 | 0.34 | 0.03 | 20 | Astrocyte |
| *LPAR1* | 5.18E-276 | 9.86E-272 | 4.00 | 0.30 | 0.02 | 20 | Astrocyte |

| Gene | P-Value | FDR | Average $log_2FC$ | Fraction 1 | Fraction 2 | Cluster | Annotation |
|---|---|---|---|---|---|---|---|
| ***KCNQ1*** | 0.00E+00 | 0.00E+00 | 3.99 | 0.56 | 0.05 | 20 | Astrocyte |
| ***SHH*** | 0.00E+00 | 0.00E+00 | 3.76 | 0.68 | 0.08 | 20 | Astrocyte |
| ***SLC15A2*** | 0.00E+00 | 0.00E+00 | 3.60 | 0.66 | 0.08 | 20 | Astrocyte |
| ***MECOM*** | 2.59E-266 | 4.92E-262 | 3.56 | 0.32 | 0.03 | 20 | Astrocyte |
| ***LOC121469790*** | 3.32E-271 | 6.31E-267 | 3.54 | 0.35 | 0.04 | 20 | Astrocyte |
| ***COL23A1*** | 1.24E-301 | 2.35E-297 | 3.52 | 0.39 | 0.06 | 20 | Astrocyte |
| ***LOC121470332*** | 1.24E-235 | 2.37E-231 | 3.36 | 0.33 | 0.04 | 20 | Astrocyte |
| ***FILIP1*** | 9.36E-220 | 1.78E-215 | 3.05 | 0.35 | 0.05 | 20 | Astrocyte |
| ***LOC100232212*** | 0.00E+00 | 0.00E+00 | 2.99 | 0.55 | 0.08 | 20 | Astrocyte |
| ***ASPG*** | 1.64E-231 | 3.11E-227 | 2.98 | 0.37 | 0.06 | 20 | Astrocyte |
| ***PCSK5*** | 0.00E+00 | 0.00E+00 | 2.90 | 0.59 | 0.14 | 20 | Astrocyte |
| ***SLIT2*** | 0.00E+00 | 0.00E+00 | 2.86 | 0.96 | 0.43 | 20 | Astrocyte |
| ***FGFRL1*** | 0.00E+00 | 0.00E+00 | 2.80 | 0.92 | 0.20 | 20 | Astrocyte |
| ***VSTM2B*** | 1.90E-191 | 3.61E-187 | 2.71 | 0.35 | 0.06 | 20 | Astrocyte |
| ***SEMA5A*** | 0.00E+00 | 0.00E+00 | 2.70 | 0.63 | 0.20 | 20 | Astrocyte |
| ***JAM2*** | 1.01E-173 | 1.92E-169 | 2.61 | 0.33 | 0.06 | 20 | Astrocyte |
| ***SMOC1*** | 0.00E+00 | 0.00E+00 | 2.61 | 0.73 | 0.18 | 20 | Astrocyte |
| ***PAX2*** | 0.00E+00 | 0.00E+00 | 7.87 | 0.86 | 0.01 | 21 | Microglia |
| ***PAX5*** | 0.00E+00 | 0.00E+00 | 6.82 | 0.42 | 0.01 | 21 | Microglia |
| ***WIPF3*** | 0.00E+00 | 0.00E+00 | 6.21 | 0.81 | 0.03 | 21 | Microglia |
| ***COL15A1*** | 0.00E+00 | 0.00E+00 | 5.62 | 0.28 | 0.01 | 21 | Microglia |
| ***RUNX1*** | 0.00E+00 | 0.00E+00 | 5.45 | 0.37 | 0.01 | 21 | Microglia |
| ***LOC115496633*** | 0.00E+00 | 0.00E+00 | 5.37 | 0.71 | 0.04 | 21 | Microglia |
| ***GDF10*** | 0.00E+00 | 0.00E+00 | 5.13 | 0.43 | 0.02 | 21 | Microglia |
| ***LOC100232659*** | 0.00E+00 | 0.00E+00 | 4.93 | 0.29 | 0.01 | 21 | Microglia |
| ***CCBE1*** | 0.00E+00 | 0.00E+00 | 4.65 | 0.63 | 0.05 | 21 | Microglia |
| ***VIT*** | 0.00E+00 | 0.00E+00 | 4.65 | 0.61 | 0.04 | 21 | Microglia |

| Gene | P-Value | FDR | Average $log_2FC$ | Fraction 1 | Fraction 2 | Cluster | Annotation |
|---|---|---|---|---|---|---|---|
| ***ADAMTS17*** | 0.00E+00 | 0.00E+00 | 4.59 | 0.68 | 0.06 | 21 | Microglia |
| ***MOXD1*** | 0.00E+00 | 0.00E+00 | 4.50 | 0.43 | 0.02 | 21 | Microglia |
| ***GYPC*** | 0.00E+00 | 0.00E+00 | 4.48 | 0.47 | 0.03 | 21 | Microglia |
| ***STON1*** | 3.63E-298 | 6.91E-294 | 4.15 | 0.32 | 0.02 | 21 | Microglia |
| ***VCL*** | 0.00E+00 | 0.00E+00 | 3.99 | 0.65 | 0.07 | 21 | Microglia |
| ***SLC4A11*** | 0.00E+00 | 0.00E+00 | 3.97 | 0.41 | 0.04 | 21 | Microglia |
| ***EDNRA*** | 1.07E-251 | 2.04E-247 | 3.92 | 0.29 | 0.02 | 21 | Microglia |
| ***CLYBL*** | 0.00E+00 | 0.00E+00 | 3.92 | 0.69 | 0.07 | 21 | Microglia |
| ***LOC115495827*** | 0.00E+00 | 0.00E+00 | 3.90 | 0.84 | 0.09 | 21 | Microglia |
| ***PAK1*** | 0.00E+00 | 0.00E+00 | 3.83 | 0.51 | 0.05 | 21 | Microglia |
| ***BCAS1*** | 0.00E+00 | 0.00E+00 | 6.61 | 0.54 | 0.02 | 22 | Oligodendrocyte |
| ***ANTXRL*** | 0.00E+00 | 0.00E+00 | 4.42 | 0.38 | 0.03 | 22 | Oligodendrocyte |
| ***BMPER*** | 0.00E+00 | 0.00E+00 | 4.27 | 0.83 | 0.20 | 22 | Oligodendrocyte |
| ***NKX2-2*** | 1.99E-273 | 3.79E-269 | 4.17 | 0.33 | 0.02 | 22 | Oligodendrocyte |
| ***PDGFRA*** | 0.00E+00 | 0.00E+00 | 3.77 | 0.50 | 0.04 | 22 | Oligodendrocyte |
| ***COLEC12*** | 0.00E+00 | 0.00E+00 | 3.70 | 0.55 | 0.06 | 22 | Oligodendrocyte |
| ***SOX8*** | 1.65E-278 | 3.14E-274 | 3.60 | 0.40 | 0.05 | 22 | Oligodendrocyte |
| ***AFAP1L2*** | 9.32E-198 | 1.77E-193 | 3.55 | 0.30 | 0.04 | 22 | Oligodendrocyte |
| ***CA8*** | 2.07E-273 | 3.94E-269 | 3.41 | 0.39 | 0.04 | 22 | Oligodendrocyte |
| ***ZCCHC24*** | 0.00E+00 | 0.00E+00 | 3.32 | 0.76 | 0.13 | 22 | Oligodendrocyte |
| ***ARHGEF3*** | 8.63E-293 | 1.64E-288 | 3.22 | 0.48 | 0.08 | 22 | Oligodendrocyte |
| ***MAML2*** | 0.00E+00 | 0.00E+00 | 3.12 | 0.92 | 0.27 | 22 | Oligodendrocyte |
| ***SLC44A5*** | 0.00E+00 | 0.00E+00 | 3.07 | 0.55 | 0.20 | 22 | Oligodendrocyte |
| ***DOCK10*** | 0.00E+00 | 0.00E+00 | 2.97 | 0.57 | 0.11 | 22 | Oligodendrocyte |
| ***EPB41L2*** | 0.00E+00 | 0.00E+00 | 2.77 | 0.44 | 0.18 | 22 | Oligodendrocyte |
| ***ZNF516*** | 1.37E-211 | 2.61E-207 | 2.68 | 0.39 | 0.11 | 22 | Oligodendrocyte |
| ***LOC121470875*** | 4.80E-138 | 9.13E-134 | 2.61 | 0.32 | 0.06 | 22 | Oligodendrocyte |

| Gene | P-Value | FDR | Average $log_2FC$ | Fraction 1 | Fraction 2 | Cluster | Annotation |
|---|---|---|---|---|---|---|---|
| ***DLL1*** | 4.76E-213 | 9.05E-209 | 2.52 | 0.48 | 0.09 | 22 | Oligodendrocyte |
| ***REPS2*** | 1.10E-161 | 2.08E-157 | 2.45 | 0.41 | 0.09 | 22 | Oligodendrocyte |
| ***BBS9*** | 0.00E+00 | 0.00E+00 | 2.42 | 0.63 | 0.28 | 22 | Oligodendrocyte |
| ***SULF1*** | 0.00E+00 | 0.00E+00 | 4.11 | 0.85 | 0.13 | 23 | Astrocyte |
| ***SLIT2*** | 0.00E+00 | 0.00E+00 | 3.90 | 0.98 | 0.43 | 23 | Astrocyte |
| ***LOC115494283*** | 1.66E-202 | 3.16E-198 | 3.78 | 0.32 | 0.03 | 23 | Astrocyte |
| ***LTBP1*** | 0.00E+00 | 0.00E+00 | 3.17 | 0.55 | 0.13 | 23 | Astrocyte |
| ***SHH*** | 5.11E-214 | 9.72E-210 | 3.02 | 0.44 | 0.09 | 23 | Astrocyte |
| ***LOC116808485*** | 2.73E-165 | 5.18E-161 | 2.86 | 0.38 | 0.06 | 23 | Astrocyte |
| ***LOC100232212*** | 1.50E-244 | 2.84E-240 | 2.80 | 0.53 | 0.09 | 23 | Astrocyte |
| ***FGFRL1*** | 0.00E+00 | 0.00E+00 | 2.62 | 0.88 | 0.21 | 23 | Astrocyte |
| ***GLIS3*** | 1.18E-179 | 2.25E-175 | 2.62 | 0.43 | 0.11 | 23 | Astrocyte |
| ***SLC1A3*** | 0.00E+00 | 0.00E+00 | 2.61 | 0.80 | 0.20 | 23 | Astrocyte |
| ***PDE1A*** | 2.61E-144 | 4.97E-140 | 2.52 | 0.38 | 0.12 | 23 | Astrocyte |
| ***LOC115493780*** | 3.49E-222 | 6.64E-218 | 2.41 | 0.58 | 0.13 | 23 | Astrocyte |
| ***SMOC1*** | 1.84E-249 | 3.50E-245 | 2.29 | 0.69 | 0.19 | 23 | Astrocyte |
| ***LMO1*** | 0.00E+00 | 0.00E+00 | 2.29 | 0.86 | 0.32 | 23 | Astrocyte |
| ***BCAR3*** | 1.75E-186 | 3.32E-182 | 2.28 | 0.56 | 0.15 | 23 | Astrocyte |
| ***IGSF11*** | 2.18E-115 | 4.15E-111 | 2.13 | 0.40 | 0.11 | 23 | Astrocyte |
| ***LAMA4*** | 2.73E-154 | 5.20E-150 | 2.03 | 0.53 | 0.15 | 23 | Astrocyte |
| ***THSD4*** | 7.31E-125 | 1.39E-120 | 1.95 | 0.49 | 0.17 | 23 | Astrocyte |
| ***AKAP13*** | 1.56E-176 | 2.96E-172 | 1.94 | 0.65 | 0.24 | 23 | Astrocyte |
| ***SLC1A2*** | 3.61E-139 | 6.87E-135 | 1.94 | 0.57 | 0.25 | 23 | Astrocyte |
| ***FOXP4*** | 1.47E-160 | 2.79E-156 | 3.48 | 0.30 | 0.04 | 24 | GABAergic Neuron |
| ***FOXP2*** | 0.00E+00 | 0.00E+00 | 3.39 | 0.96 | 0.37 | 24 | GABAergic Neuron |
| ***TSHZ1*** | 0.00E+00 | 0.00E+00 | 3.02 | 0.61 | 0.15 | 24 | GABAergic Neuron |
| ***RUNX1T1*** | 0.00E+00 | 0.00E+00 | 2.66 | 0.94 | 0.37 | 24 | GABAergic Neuron |

| Gene | P-Value | FDR | Average $log_2FC$ | Fraction 1 | Fraction 2 | Cluster | Annotation |
|---|---|---|---|---|---|---|---|
| *ADARB2* | 0.00E+00 | 0.00E+00 | 2.66 | 0.95 | 0.23 | 24 | GABAergic Neuron |
| *LOC121469248* | 9.17E-215 | 1.74E-210 | 2.65 | 0.51 | 0.12 | 24 | GABAergic Neuron |
| *MEIS2* | 0.00E+00 | 0.00E+00 | 2.60 | 0.98 | 0.31 | 24 | GABAergic Neuron |
| *CHRM3* | 2.06E-152 | 3.92E-148 | 2.24 | 0.51 | 0.22 | 24 | GABAergic Neuron |
| *CACNA2D3* | 1.47E-179 | 2.80E-175 | 2.09 | 0.66 | 0.24 | 24 | GABAergic Neuron |
| *MPPED2* | 1.01E-130 | 1.92E-126 | 1.94 | 0.48 | 0.20 | 24 | GABAergic Neuron |
| *DSCAM* | 1.84E-314 | 3.49E-310 | 1.93 | 0.94 | 0.45 | 24 | GABAergic Neuron |
| *FLRT2* | 1.27E-134 | 2.41E-130 | 1.92 | 0.57 | 0.22 | 24 | GABAergic Neuron |
| *GALNTL6* | 3.93E-163 | 7.47E-159 | 1.88 | 0.68 | 0.32 | 24 | GABAergic Neuron |
| *TSPAN9* | 4.28E-101 | 8.14E-97 | 1.76 | 0.46 | 0.21 | 24 | GABAergic Neuron |
| *PBX3* | 3.14E-192 | 5.97E-188 | 1.73 | 0.71 | 0.40 | 24 | GABAergic Neuron |
| *DLG2* | 1.76E-319 | 3.35E-315 | 1.68 | 0.96 | 0.65 | 24 | GABAergic Neuron |
| *SOX2* | 5.01E-230 | 9.52E-226 | 1.51 | 0.95 | 0.70 | 24 | GABAergic Neuron |
| *GRIP2* | 2.65E-69 | 5.04E-65 | 1.48 | 0.47 | 0.21 | 24 | GABAergic Neuron |
| *CCSER1* | 2.55E-163 | 4.86E-159 | 1.26 | 0.91 | 0.58 | 24 | GABAergic Neuron |
| *LOC115494452* | 7.35E-106 | 1.40E-101 | 1.26 | 0.74 | 0.42 | 24 | GABAergic Neuron |
| *LOC121470554* | 3.02E-169 | 5.74E-165 | 3.89 | 0.32 | 0.03 | 25 | Astrocyte |
| *TNC* | 2.66E-153 | 5.05E-149 | 3.12 | 0.40 | 0.07 | 25 | Astrocyte |
| *GLI2* | 3.75E-218 | 7.13E-214 | 2.93 | 0.56 | 0.09 | 25 | Astrocyte |
| *LOC105758645* | 0.00E+00 | 0.00E+00 | 2.76 | 0.84 | 0.29 | 25 | Astrocyte |
| *GLI3* | 0.00E+00 | 0.00E+00 | 2.65 | 0.83 | 0.18 | 25 | Astrocyte |
| *LOC115495208* | 1.34E-96 | 2.56E-92 | 2.62 | 0.31 | 0.06 | 25 | Astrocyte |
| *BMPR1B* | 3.94E-135 | 7.50E-131 | 2.22 | 0.50 | 0.12 | 25 | Astrocyte |
| *LRP2* | 3.57E-86 | 6.79E-82 | 2.21 | 0.35 | 0.09 | 25 | Astrocyte |
| *PRDM16* | 2.23E-259 | 4.24E-255 | 2.20 | 0.83 | 0.24 | 25 | Astrocyte |
| *SALL1* | 1.63E-88 | 3.09E-84 | 2.04 | 0.39 | 0.11 | 25 | Astrocyte |
| *KIAA1217* | 6.20E-116 | 1.18E-111 | 2.02 | 0.49 | 0.13 | 25 | Astrocyte |

| Gene | P-Value | FDR | Average $log_2FC$ | Fraction 1 | Fraction 2 | Cluster | Annotation |
|---|---|---|---|---|---|---|---|
| *LOC115495827* | 1.70E-93 | 3.23E-89 | 1.99 | 0.40 | 0.10 | 25 | Astrocyte |
| *LAMA4* | 3.26E-86 | 6.20E-82 | 1.95 | 0.45 | 0.15 | 25 | Astrocyte |
| *MLC1* | 6.44E-135 | 1.22E-130 | 1.94 | 0.58 | 0.16 | 25 | Astrocyte |
| *SMOC1* | 2.01E-122 | 3.82E-118 | 1.92 | 0.58 | 0.19 | 25 | Astrocyte |
| *LOC115493780* | 4.26E-93 | 8.10E-89 | 1.91 | 0.44 | 0.13 | 25 | Astrocyte |
| *PLPP3* | 7.48E-148 | 1.42E-143 | 1.90 | 0.64 | 0.19 | 25 | Astrocyte |
| *MEGF10* | 2.84E-73 | 5.41E-69 | 1.89 | 0.38 | 0.12 | 25 | Astrocyte |
| *P4HA1* | 5.18E-120 | 9.84E-116 | 1.87 | 0.58 | 0.22 | 25 | Astrocyte |
| *LFNG* | 4.20E-98 | 7.98E-94 | 1.84 | 0.49 | 0.15 | 25 | Astrocyte |
| *SOX14* | 8.24E-182 | 1.57E-177 | 4.83 | 0.26 | 0.01 | 26 | Glutamatergic/GABA |
| *MTUS2* | 0.00E+00 | 0.00E+00 | 3.97 | 0.63 | 0.13 | 26 | Glutamatergic/GABA |
| *ZFPM2* | 0.00E+00 | 0.00E+00 | 3.56 | 0.57 | 0.17 | 26 | Glutamatergic/GABA |
| *EBF3* | 2.14E-156 | 4.06E-152 | 3.38 | 0.37 | 0.05 | 26 | Glutamatergic/GABA |
| *LEF1* | 9.61E-131 | 1.83E-126 | 2.98 | 0.32 | 0.04 | 26 | Glutamatergic/GABA |
| *LOC100223410* | 0.00E+00 | 0.00E+00 | 2.97 | 0.75 | 0.30 | 26 | Glutamatergic/GABA |
| *ROBO2* | 8.85E-313 | 1.68E-308 | 2.68 | 0.81 | 0.39 | 26 | Glutamatergic/GABA |
| *CRHR2* | 1.60E-103 | 3.04E-99 | 2.64 | 0.38 | 0.10 | 26 | Glutamatergic/GABA |
| *FAT3* | 0.00E+00 | 0.00E+00 | 2.58 | 0.84 | 0.33 | 26 | Glutamatergic/GABA |
| *LOC100218233* | 2.88E-105 | 5.48E-101 | 2.25 | 0.40 | 0.12 | 26 | Glutamatergic/GABA |
| *LOC100230699* | 2.83E-239 | 5.37E-235 | 2.14 | 0.71 | 0.33 | 26 | Glutamatergic/GABA |
| *DAB1* | 6.31E-207 | 1.20E-202 | 2.14 | 0.70 | 0.34 | 26 | Glutamatergic/GABA |
| *DSCAM* | 9.95E-255 | 1.89E-250 | 2.05 | 0.74 | 0.46 | 26 | Glutamatergic/GABA |
| *THSD7A* | 2.63E-124 | 4.99E-120 | 2.00 | 0.55 | 0.22 | 26 | Glutamatergic/GABA |
| *ADCY1* | 6.69E-124 | 1.27E-119 | 1.96 | 0.50 | 0.23 | 26 | Glutamatergic/GABA |
| *EBF1* | 1.06E-88 | 2.02E-84 | 1.88 | 0.39 | 0.10 | 26 | Glutamatergic/GABA |
| *RBFOX1* | 8.46E-232 | 1.61E-227 | 1.87 | 0.92 | 0.49 | 26 | Glutamatergic/GABA |
| *HS3ST5* | 4.52E-67 | 8.58E-63 | 1.82 | 0.42 | 0.17 | 26 | Glutamatergic/GABA |

| Gene | P-Value | FDR | Average $log_2FC$ | Fraction 1 | Fraction 2 | Cluster | Annotation |
|---|---|---|---|---|---|---|---|
| ***GRIP2*** | 8.80E-86 | 1.67E-81 | 1.79 | 0.47 | 0.21 | 26 | Glutamatergic/GABA |
| ***SIPA1L2*** | 2.32E-89 | 4.42E-85 | 1.74 | 0.49 | 0.23 | 26 | Glutamatergic/GABA |
| ***FCN2*** | 0.00E+00 | 0.00E+00 | 9.22 | 0.37 | 0.00 | 27 | Ependymal Cell |
| ***LOC115497384*** | 6.00E-323 | 1.13E-318 | 9.14 | 0.28 | 0.00 | 27 | Ependymal Cell |
| ***CDHR4*** | 0.00E+00 | 0.00E+00 | 8.48 | 0.30 | 0.00 | 27 | Ependymal Cell |
| ***LOC121471091*** | 0.00E+00 | 0.00E+00 | 8.44 | 0.68 | 0.00 | 27 | Ependymal Cell |
| ***SMIM35*** | 3.82E-264 | 7.27E-260 | 8.10 | 0.25 | 0.00 | 27 | Ependymal Cell |
| ***CCDC187*** | 1.12E-269 | 2.14E-265 | 8.01 | 0.26 | 0.00 | 27 | Ependymal Cell |
| ***LOC116808114*** | 7.72E-259 | 1.47E-254 | 6.86 | 0.28 | 0.00 | 27 | Ependymal Cell |
| ***CYP11A1*** | 0.00E+00 | 0.00E+00 | 6.58 | 0.41 | 0.01 | 27 | Ependymal Cell |
| ***SPEF1*** | 1.29E-232 | 2.45E-228 | 6.50 | 0.26 | 0.00 | 27 | Ependymal Cell |
| ***LRRC43*** | 3.07E-264 | 5.84E-260 | 6.47 | 0.30 | 0.00 | 27 | Ependymal Cell |
| ***DEUP1*** | 0.00E+00 | 0.00E+00 | 6.37 | 0.53 | 0.01 | 27 | Ependymal Cell |
| ***MYB*** | 0.00E+00 | 0.00E+00 | 6.29 | 0.56 | 0.01 | 27 | Ependymal Cell |
| ***LOC121470873*** | 6.68E-293 | 1.27E-288 | 6.29 | 0.34 | 0.01 | 27 | Ependymal Cell |
| ***FHAD1*** | 3.13E-260 | 5.94E-256 | 6.24 | 0.30 | 0.01 | 27 | Ependymal Cell |
| ***ANKRD66*** | 3.58E-268 | 6.80E-264 | 6.16 | 0.32 | 0.01 | 27 | Ependymal Cell |
| ***LOC100225564*** | 0.00E+00 | 0.00E+00 | 6.15 | 0.45 | 0.01 | 27 | Ependymal Cell |
| ***KIF6*** | 0.00E+00 | 0.00E+00 | 6.02 | 0.58 | 0.02 | 27 | Ependymal Cell |
| ***LOC115495985*** | 1.33E-221 | 2.53E-217 | 5.92 | 0.28 | 0.01 | 27 | Ependymal Cell |
| ***CCNO*** | 6.07E-238 | 1.15E-233 | 5.90 | 0.30 | 0.01 | 27 | Ependymal Cell |
| ***LOC115495504*** | 4.51E-295 | 8.57E-291 | 5.87 | 0.36 | 0.01 | 27 | Ependymal Cell |
| ***SLC17A8*** | 1.45E-219 | 2.75E-215 | 6.17 | 0.27 | 0.01 | 28 | Glutamatergic Neuron |
| ***RUNX2*** | 9.70E-235 | 1.84E-230 | 4.65 | 0.32 | 0.03 | 28 | Glutamatergic Neuron |
| ***DDC*** | 1.02E-184 | 1.93E-180 | 3.88 | 0.42 | 0.05 | 28 | Glutamatergic Neuron |
| ***RSPO2*** | 7.39E-128 | 1.40E-123 | 3.56 | 0.28 | 0.02 | 28 | Glutamatergic Neuron |
| ***PROX1*** | 1.34E-202 | 2.55E-198 | 3.10 | 0.55 | 0.13 | 28 | Glutamatergic Neuron |

| Gene | P-Value | FDR | Average $log_2FC$ | Fraction 1 | Fraction 2 | Cluster | Annotation |
|---|---|---|---|---|---|---|---|
| ***NKD1*** | 8.60E-178 | 1.63E-173 | 2.98 | 0.52 | 0.11 | 28 | Glutamatergic Neuron |
| ***SYT10*** | 4.32E-152 | 8.22E-148 | 2.95 | 0.49 | 0.10 | 28 | Glutamatergic Neuron |
| ***NR2E1*** | 1.06E-116 | 2.01E-112 | 2.89 | 0.36 | 0.07 | 28 | Glutamatergic Neuron |
| ***PPFIBP1*** | 2.29E-280 | 4.35E-276 | 2.83 | 0.40 | 0.12 | 28 | Glutamatergic Neuron |
| ***LEF1*** | 1.83E-105 | 3.49E-101 | 2.76 | 0.31 | 0.04 | 28 | Glutamatergic Neuron |
| ***PLCL1*** | 2.39E-229 | 4.55E-225 | 2.65 | 0.64 | 0.22 | 28 | Glutamatergic Neuron |
| ***VAV3*** | 1.30E-113 | 2.48E-109 | 2.58 | 0.38 | 0.11 | 28 | Glutamatergic Neuron |
| ***LHX1*** | 7.34E-121 | 1.40E-116 | 2.57 | 0.39 | 0.06 | 28 | Glutamatergic Neuron |
| ***LOC100232083*** | 8.00E-92 | 1.52E-87 | 2.56 | 0.36 | 0.08 | 28 | Glutamatergic Neuron |
| ***EYS*** | 1.03E-237 | 1.96E-233 | 2.45 | 0.78 | 0.20 | 28 | Glutamatergic Neuron |
| ***PRDX1*** | 6.70E-267 | 1.27E-262 | 2.42 | 0.47 | 0.21 | 28 | Glutamatergic Neuron |
| ***THSD7A*** | 2.15E-166 | 4.10E-162 | 2.41 | 0.56 | 0.22 | 28 | Glutamatergic Neuron |
| ***DACH1*** | 1.64E-285 | 3.12E-281 | 2.32 | 0.87 | 0.41 | 28 | Glutamatergic Neuron |
| ***FSTL4*** | 1.19E-119 | 2.26E-115 | 2.27 | 0.51 | 0.20 | 28 | Glutamatergic Neuron |
| ***SLCO3A1*** | 2.38E-117 | 4.52E-113 | 2.25 | 0.46 | 0.13 | 28 | Glutamatergic Neuron |
| ***TBR1*** | 1.78E-161 | 3.38E-157 | 4.15 | 0.31 | 0.02 | 29 | Glutamatergic Neuron |
| ***CHODL*** | 6.18E-248 | 1.17E-243 | 4.11 | 0.47 | 0.05 | 29 | Glutamatergic Neuron |
| ***LOC121470782*** | 1.19E-147 | 2.27E-143 | 4.03 | 0.30 | 0.02 | 29 | Glutamatergic Neuron |
| ***RELN*** | 0.00E+00 | 0.00E+00 | 3.77 | 0.92 | 0.19 | 29 | Glutamatergic Neuron |
| ***ROBO2*** | 0.00E+00 | 0.00E+00 | 3.44 | 0.87 | 0.39 | 29 | Glutamatergic Neuron |
| ***NXPH1*** | 0.00E+00 | 0.00E+00 | 3.30 | 0.97 | 0.34 | 29 | Glutamatergic Neuron |
| ***LOC121469439*** | 3.11E-214 | 5.91E-210 | 3.29 | 0.53 | 0.07 | 29 | Glutamatergic Neuron |
| ***TCF7L2*** | 0.00E+00 | 0.00E+00 | 3.22 | 0.91 | 0.27 | 29 | Glutamatergic Neuron |
| ***CACNA2D1*** | 0.00E+00 | 0.00E+00 | 3.22 | 0.97 | 0.43 | 29 | Glutamatergic Neuron |
| ***LOC100229402*** | 0.00E+00 | 0.00E+00 | 3.17 | 0.94 | 0.28 | 29 | Glutamatergic Neuron |
| ***TMEM272*** | 2.67E-100 | 5.08E-96 | 3.00 | 0.30 | 0.04 | 29 | Glutamatergic Neuron |
| ***LOC115497399*** | 2.90E-103 | 5.51E-99 | 2.89 | 0.33 | 0.05 | 29 | Glutamatergic Neuron |

| Gene | P-Value | FDR | Average $log_2FC$ | Fraction 1 | Fraction 2 | Cluster | Annotation |
|---|---|---|---|---|---|---|---|
| *PHF14* | 0.00E+00 | 0.00E+00 | 2.83 | 0.95 | 0.50 | 29 | Glutamatergic Neuron |
| *GALNTL6* | 0.00E+00 | 0.00E+00 | 2.78 | 0.89 | 0.32 | 29 | Glutamatergic Neuron |
| *PTPRM* | 9.01E-270 | 1.71E-265 | 2.60 | 0.83 | 0.25 | 29 | Glutamatergic Neuron |
| *THSD7A* | 2.53E-181 | 4.81E-177 | 2.39 | 0.66 | 0.22 | 29 | Glutamatergic Neuron |
| *PCDH10* | 0.00E+00 | 0.00E+00 | 2.35 | 0.90 | 0.33 | 29 | Glutamatergic Neuron |
| *DACH1* | 1.49E-307 | 2.83E-303 | 2.32 | 0.90 | 0.41 | 29 | Glutamatergic Neuron |
| *LOC121469488* | 5.41E-109 | 1.03E-104 | 2.19 | 0.49 | 0.14 | 29 | Glutamatergic Neuron |
| *KIF26A* | 5.34E-127 | 1.02E-122 | 2.12 | 0.59 | 0.17 | 29 | Glutamatergic Neuron |
| *TRPC4* | 0.00E+00 | 0.00E+00 | 3.95 | 0.52 | 0.08 | 3 | Glutamatergic/GABA |
| *SHISA6* | 0.00E+00 | 0.00E+00 | 2.35 | 0.50 | 0.14 | 3 | Glutamatergic/GABA |
| *PRDM16* | 0.00E+00 | 0.00E+00 | 2.09 | 0.72 | 0.22 | 3 | Glutamatergic/GABA |
| *PPP4R4* | 0.00E+00 | 0.00E+00 | 2.09 | 0.46 | 0.14 | 3 | Glutamatergic/GABA |
| *LOC100221337* | 0.00E+00 | 0.00E+00 | 2.09 | 0.46 | 0.15 | 3 | Glutamatergic/GABA |
| *GRM5* | 0.00E+00 | 0.00E+00 | 2.01 | 0.75 | 0.32 | 3 | Glutamatergic/GABA |
| *TRPC5* | 9.23E-319 | 1.75E-314 | 1.98 | 0.39 | 0.12 | 3 | Glutamatergic/GABA |
| *GRIP2* | 0.00E+00 | 0.00E+00 | 1.84 | 0.60 | 0.19 | 3 | Glutamatergic/GABA |
| *KCNJ6* | 0.00E+00 | 0.00E+00 | 1.78 | 0.51 | 0.17 | 3 | Glutamatergic/GABA |
| *PEX5L* | 2.43E-289 | 4.63E-285 | 1.77 | 0.47 | 0.18 | 3 | Glutamatergic/GABA |
| *FAM135B* | 0.00E+00 | 0.00E+00 | 1.66 | 0.57 | 0.23 | 3 | Glutamatergic/GABA |
| *TENM2* | 0.00E+00 | 0.00E+00 | 1.63 | 0.87 | 0.46 | 3 | Glutamatergic/GABA |
| *DAB1* | 0.00E+00 | 0.00E+00 | 1.57 | 0.68 | 0.32 | 3 | Glutamatergic/GABA |
| *MEIS2* | 0.00E+00 | 0.00E+00 | 1.50 | 0.66 | 0.30 | 3 | Glutamatergic/GABA |
| *LOC101234143* | 1.96E-214 | 3.74E-210 | 1.48 | 0.42 | 0.17 | 3 | Glutamatergic/GABA |
| *ARPP21* | 0.00E+00 | 0.00E+00 | 1.48 | 0.95 | 0.51 | 3 | Glutamatergic/GABA |
| *GABRG3* | 2.40E-291 | 4.55E-287 | 1.45 | 0.55 | 0.23 | 3 | Glutamatergic/GABA |
| *CCSER1* | 0.00E+00 | 0.00E+00 | 1.39 | 0.93 | 0.57 | 3 | Glutamatergic/GABA |
| *TMEM132B* | 1.49E-247 | 2.84E-243 | 1.38 | 0.53 | 0.24 | 3 | Glutamatergic/GABA |

| Gene | P-Value | FDR | Average $log_2FC$ | Fraction 1 | Fraction 2 | Cluster | Annotation |
|---|---|---|---|---|---|---|---|
| *RAPGEF4* | 3.36E-242 | 6.39E-238 | 1.36 | 0.50 | 0.21 | 3 | Glutamatergic/GABA |
| *NDNF* | 0.00E+00 | 0.00E+00 | 5.99 | 0.64 | 0.03 | 30 | Astrocyte |
| *ASPA* | 9.81E-143 | 1.86E-138 | 4.21 | 0.31 | 0.02 | 30 | Astrocyte |
| *LOC115491117* | 4.87E-157 | 9.25E-153 | 4.14 | 0.34 | 0.02 | 30 | Astrocyte |
| *STON1* | 6.54E-134 | 1.24E-129 | 3.96 | 0.31 | 0.02 | 30 | Astrocyte |
| *LRP4* | 7.10E-265 | 1.35E-260 | 3.90 | 0.58 | 0.07 | 30 | Astrocyte |
| *CDH19* | 3.69E-127 | 7.02E-123 | 3.76 | 0.31 | 0.02 | 30 | Astrocyte |
| *LOC115495827* | 0.00E+00 | 0.00E+00 | 3.73 | 0.83 | 0.10 | 30 | Astrocyte |
| *LOC115497553* | 7.85E-142 | 1.49E-137 | 3.69 | 0.35 | 0.03 | 30 | Astrocyte |
| *ADAMTS17* | 1.78E-205 | 3.39E-201 | 3.63 | 0.54 | 0.07 | 30 | Astrocyte |
| *KANK1* | 4.64E-183 | 8.81E-179 | 3.59 | 0.47 | 0.05 | 30 | Astrocyte |
| *DNAH5* | 0.00E+00 | 0.00E+00 | 3.55 | 0.88 | 0.18 | 30 | Astrocyte |
| *COL26A1* | 5.51E-110 | 1.05E-105 | 3.28 | 0.33 | 0.04 | 30 | Astrocyte |
| *COL5A1* | 1.91E-310 | 3.63E-306 | 3.24 | 0.79 | 0.11 | 30 | Astrocyte |
| *LOC100190440* | 2.44E-133 | 4.65E-129 | 3.24 | 0.39 | 0.05 | 30 | Astrocyte |
| *VIT* | 6.32E-135 | 1.20E-130 | 3.22 | 0.40 | 0.05 | 30 | Astrocyte |
| *GREB1* | 2.58E-126 | 4.90E-122 | 3.17 | 0.39 | 0.05 | 30 | Astrocyte |
| *LMO4* | 1.18E-140 | 2.24E-136 | 3.16 | 0.41 | 0.06 | 30 | Astrocyte |
| *CLYBL* | 1.26E-197 | 2.39E-193 | 3.11 | 0.59 | 0.08 | 30 | Astrocyte |
| *BMPR1B* | 3.40E-208 | 6.46E-204 | 3.07 | 0.65 | 0.12 | 30 | Astrocyte |
| *SAT1* | 2.06E-123 | 3.92E-119 | 3.02 | 0.41 | 0.08 | 30 | Astrocyte |
| *POU6F2* | 4.04E-139 | 7.68E-135 | 3.38 | 0.43 | 0.08 | 31 | Glutamatergic Neuron |
| *MKX* | 2.93E-82 | 5.57E-78 | 2.88 | 0.32 | 0.05 | 31 | Glutamatergic Neuron |
| *LOC115495531* | 9.33E-94 | 1.77E-89 | 2.73 | 0.36 | 0.05 | 31 | Glutamatergic Neuron |
| *LOC115496015* | 1.07E-96 | 2.03E-92 | 2.62 | 0.44 | 0.10 | 31 | Glutamatergic Neuron |
| *FIGN* | 5.06E-160 | 9.62E-156 | 2.41 | 0.62 | 0.23 | 31 | Glutamatergic Neuron |
| *GRIP2* | 7.28E-146 | 1.38E-141 | 2.29 | 0.69 | 0.21 | 31 | Glutamatergic Neuron |

| Gene | P-Value | FDR | Average $log_2FC$ | Fraction 1 | Fraction 2 | Cluster | Annotation |
|---|---|---|---|---|---|---|---|
| ***VSTM2A*** | 3.50E-136 | 6.65E-132 | 2.25 | 0.66 | 0.17 | 31 | Glutamatergic Neuron |
| ***ZNF804A*** | 4.51E-125 | 8.58E-121 | 2.19 | 0.67 | 0.25 | 31 | Glutamatergic Neuron |
| ***RNF220*** | 2.49E-68 | 4.73E-64 | 2.17 | 0.39 | 0.12 | 31 | Glutamatergic Neuron |
| ***COL25A1*** | 1.02E-69 | 1.95E-65 | 2.16 | 0.44 | 0.13 | 31 | Glutamatergic Neuron |
| ***MEIS2*** | 4.78E-235 | 9.08E-231 | 2.13 | 0.93 | 0.31 | 31 | Glutamatergic Neuron |
| ***ISL1*** | 4.48E-118 | 8.52E-114 | 2.11 | 0.59 | 0.14 | 31 | Glutamatergic Neuron |
| ***LOC115493811*** | 5.02E-67 | 9.54E-63 | 2.04 | 0.43 | 0.12 | 31 | Glutamatergic Neuron |
| ***GABRG3*** | 6.39E-110 | 1.21E-105 | 1.98 | 0.68 | 0.24 | 31 | Glutamatergic Neuron |
| ***GALNT9*** | 1.10E-72 | 2.10E-68 | 1.93 | 0.48 | 0.14 | 31 | Glutamatergic Neuron |
| ***CRHR2*** | 4.11E-51 | 7.82E-47 | 1.93 | 0.35 | 0.10 | 31 | Glutamatergic Neuron |
| ***PTPRD*** | 1.03E-173 | 1.96E-169 | 1.90 | 0.90 | 0.62 | 31 | Glutamatergic Neuron |
| ***ZNF804B*** | 1.04E-52 | 1.97E-48 | 1.88 | 0.44 | 0.19 | 31 | Glutamatergic Neuron |
| ***IL1RAPL2*** | 2.81E-58 | 5.35E-54 | 1.87 | 0.41 | 0.12 | 31 | Glutamatergic Neuron |
| ***CNTN5*** | 2.25E-106 | 4.27E-102 | 1.87 | 0.72 | 0.27 | 31 | Glutamatergic Neuron |
| ***CPED1*** | 0.00E+00 | 0.00E+00 | 8.18 | 0.60 | 0.01 | 32 | Fibroblast |
| ***LOC100224093*** | 0.00E+00 | 0.00E+00 | 7.70 | 0.49 | 0.01 | 32 | Fibroblast |
| ***LOC100225050*** | 1.43E-195 | 2.72E-191 | 7.48 | 0.28 | 0.00 | 32 | Fibroblast |
| ***COL6A3*** | 0.00E+00 | 0.00E+00 | 7.32 | 0.65 | 0.01 | 32 | Fibroblast |
| ***SLC13A3*** | 0.00E+00 | 0.00E+00 | 6.60 | 0.56 | 0.04 | 32 | Fibroblast |
| ***DCN*** | 4.67E-310 | 8.88E-306 | 6.36 | 0.50 | 0.01 | 32 | Fibroblast |
| ***COL3A1*** | 3.80E-305 | 7.22E-301 | 6.33 | 0.50 | 0.01 | 32 | Fibroblast |
| ***LOC100230220*** | 8.12E-235 | 1.54E-230 | 6.33 | 0.37 | 0.01 | 32 | Fibroblast |
| ***LOC100223866*** | 3.12E-158 | 5.92E-154 | 6.29 | 0.27 | 0.01 | 32 | Fibroblast |
| ***LOC116807725*** | 5.02E-149 | 9.54E-145 | 6.21 | 0.26 | 0.01 | 32 | Fibroblast |
| ***LOC100220148*** | 1.08E-267 | 2.05E-263 | 5.92 | 0.47 | 0.01 | 32 | Fibroblast |
| ***COL1A2*** | 0.00E+00 | 0.00E+00 | 5.91 | 0.67 | 0.03 | 32 | Fibroblast |
| ***APOA1*** | 0.00E+00 | 0.00E+00 | 5.79 | 0.48 | 0.04 | 32 | Fibroblast |

| Gene | P-Value | FDR | Average $log_2FC$ | Fraction 1 | Fraction 2 | Cluster | Annotation |
|---|---|---|---|---|---|---|---|
| ***RAMP3*** | 1.58E-205 | 3.00E-201 | 5.77 | 0.36 | 0.01 | 32 | Fibroblast |
| ***TBX18*** | 1.51E-209 | 2.88E-205 | 5.68 | 0.38 | 0.01 | 32 | Fibroblast |
| ***LAMA2*** | 0.00E+00 | 0.00E+00 | 5.48 | 0.66 | 0.05 | 32 | Fibroblast |
| ***PI15*** | 1.53E-192 | 2.90E-188 | 5.43 | 0.37 | 0.02 | 32 | Fibroblast |
| ***ADGRD1*** | 2.17E-269 | 4.12E-265 | 5.32 | 0.47 | 0.02 | 32 | Fibroblast |
| ***SIDT1*** | 2.04E-257 | 3.87E-253 | 4.98 | 0.40 | 0.03 | 32 | Fibroblast |
| ***PKP2*** | 2.35E-167 | 4.46E-163 | 4.85 | 0.33 | 0.02 | 32 | Fibroblast |
| ***RSPO2*** | 0.00E+00 | 0.00E+00 | 7.41 | 0.69 | 0.02 | 33 | Astrocyte |
| ***ANGPT1*** | 0.00E+00 | 0.00E+00 | 5.24 | 0.69 | 0.03 | 33 | Astrocyte |
| ***LMX1B*** | 0.00E+00 | 0.00E+00 | 5.23 | 0.68 | 0.03 | 33 | Astrocyte |
| ***WNT7B*** | 6.52E-268 | 1.24E-263 | 4.51 | 0.64 | 0.04 | 33 | Astrocyte |
| ***BAG2*** | 1.62E-145 | 3.08E-141 | 4.47 | 0.34 | 0.02 | 33 | Astrocyte |
| ***PITX2*** | 2.76E-100 | 5.25E-96 | 4.19 | 0.28 | 0.02 | 33 | Astrocyte |
| ***LOC115494283*** | 1.84E-100 | 3.49E-96 | 3.73 | 0.33 | 0.03 | 33 | Astrocyte |
| ***GPC5*** | 1.51E-296 | 2.87E-292 | 3.69 | 0.83 | 0.13 | 33 | Astrocyte |
| ***SULF1*** | 9.96E-258 | 1.89E-253 | 3.37 | 0.84 | 0.13 | 33 | Astrocyte |
| ***NAALADL2*** | 1.25E-84 | 2.38E-80 | 3.30 | 0.33 | 0.04 | 33 | Astrocyte |
| ***LOC100232212*** | 9.14E-185 | 1.74E-180 | 3.22 | 0.65 | 0.09 | 33 | Astrocyte |
| ***GLIS3*** | 2.70E-178 | 5.12E-174 | 3.18 | 0.57 | 0.11 | 33 | Astrocyte |
| ***CELSR1*** | 9.70E-207 | 1.84E-202 | 3.16 | 0.72 | 0.11 | 33 | Astrocyte |
| ***TIMP4*** | 3.82E-143 | 7.26E-139 | 3.15 | 0.53 | 0.09 | 33 | Astrocyte |
| ***LOC115493495*** | 8.78E-120 | 1.67E-115 | 3.15 | 0.47 | 0.06 | 33 | Astrocyte |
| ***NTN1*** | 5.96E-87 | 1.13E-82 | 3.13 | 0.36 | 0.05 | 33 | Astrocyte |
| ***ADAM12*** | 3.99E-97 | 7.59E-93 | 3.00 | 0.41 | 0.06 | 33 | Astrocyte |
| ***CREB5*** | 1.23E-219 | 2.35E-215 | 2.89 | 0.82 | 0.20 | 33 | Astrocyte |
| ***SEMA3A*** | 4.43E-122 | 8.42E-118 | 2.88 | 0.54 | 0.09 | 33 | Astrocyte |
| ***TNC*** | 1.65E-101 | 3.14E-97 | 2.82 | 0.45 | 0.07 | 33 | Astrocyte |

| Gene | P-Value | FDR | Average $log_2FC$ | Fraction 1 | Fraction 2 | Cluster | Annotation |
|---|---|---|---|---|---|---|---|
| ***LOC121470336*** | 2.69E-113 | 5.12E-109 | 4.64 | 0.30 | 0.01 | 34 | Glutamatergic Neuron |
| ***LOC115496198*** | 1.46E-105 | 2.77E-101 | 4.13 | 0.32 | 0.03 | 34 | Glutamatergic Neuron |
| ***LHX9*** | 1.60E-133 | 3.03E-129 | 4.01 | 0.42 | 0.03 | 34 | Glutamatergic Neuron |
| ***RNF220*** | 1.31E-214 | 2.49E-210 | 3.38 | 0.54 | 0.12 | 34 | Glutamatergic Neuron |
| ***MGAT4C*** | 1.29E-168 | 2.44E-164 | 2.83 | 0.82 | 0.23 | 34 | Glutamatergic Neuron |
| ***NWD2*** | 2.99E-56 | 5.68E-52 | 2.77 | 0.31 | 0.05 | 34 | Glutamatergic Neuron |
| ***SLC17A6*** | 2.96E-72 | 5.64E-68 | 2.33 | 0.46 | 0.10 | 34 | Glutamatergic Neuron |
| ***GAS2*** | 1.02E-56 | 1.93E-52 | 2.29 | 0.39 | 0.10 | 34 | Glutamatergic Neuron |
| ***TCF7L2*** | 5.74E-120 | 1.09E-115 | 2.27 | 0.63 | 0.27 | 34 | Glutamatergic Neuron |
| ***ERC2*** | 8.98E-194 | 1.71E-189 | 2.25 | 0.90 | 0.41 | 34 | Glutamatergic Neuron |
| ***ADCYAP1*** | 8.50E-59 | 1.62E-54 | 2.17 | 0.39 | 0.09 | 34 | Glutamatergic Neuron |
| ***VAV3*** | 1.11E-49 | 2.11E-45 | 2.16 | 0.38 | 0.11 | 34 | Glutamatergic Neuron |
| ***HS3ST5*** | 1.00E-73 | 1.91E-69 | 2.15 | 0.57 | 0.17 | 34 | Glutamatergic Neuron |
| ***TMEM163*** | 1.84E-59 | 3.50E-55 | 2.10 | 0.43 | 0.10 | 34 | Glutamatergic Neuron |
| ***DENND1B*** | 1.79E-81 | 3.40E-77 | 2.06 | 0.56 | 0.20 | 34 | Glutamatergic Neuron |
| ***SHISAL1*** | 1.18E-57 | 2.24E-53 | 2.04 | 0.45 | 0.16 | 34 | Glutamatergic Neuron |
| ***C4H4orf50*** | 2.93E-75 | 5.57E-71 | 1.95 | 0.59 | 0.20 | 34 | Glutamatergic Neuron |
| ***JAKMIP1*** | 6.81E-93 | 1.30E-88 | 1.92 | 0.68 | 0.25 | 34 | Glutamatergic Neuron |
| ***NTNG1*** | 3.72E-69 | 7.07E-65 | 1.86 | 0.62 | 0.27 | 34 | Glutamatergic Neuron |
| ***CNTN5*** | 1.84E-58 | 3.50E-54 | 1.84 | 0.63 | 0.27 | 34 | Glutamatergic Neuron |
| ***TFAP2D*** | 1.62E-187 | 3.07E-183 | 6.93 | 0.33 | 0.00 | 35 | Glutamatergic Neuron |
| ***RSPO3*** | 3.07E-164 | 5.84E-160 | 5.56 | 0.37 | 0.01 | 35 | Glutamatergic Neuron |
| ***TBR1*** | 2.16E-191 | 4.11E-187 | 5.01 | 0.47 | 0.02 | 35 | Glutamatergic Neuron |
| ***FIGN*** | 0.00E+00 | 0.00E+00 | 4.55 | 0.89 | 0.23 | 35 | Glutamatergic Neuron |
| ***CRHR2*** | 6.08E-240 | 1.16E-235 | 3.85 | 0.73 | 0.10 | 35 | Glutamatergic Neuron |
| ***PLCL1*** | 2.90E-317 | 5.51E-313 | 3.42 | 0.82 | 0.22 | 35 | Glutamatergic Neuron |
| ***MCTP2*** | 1.50E-70 | 2.85E-66 | 3.38 | 0.30 | 0.04 | 35 | Glutamatergic Neuron |

| Gene | P-Value | FDR | Average $log_2FC$ | Fraction 1 | Fraction 2 | Cluster | Annotation |
|---|---|---|---|---|---|---|---|
| *AFF2* | 0.00E+00 | 0.00E+00 | 2.90 | 0.95 | 0.42 | 35 | Glutamatergic Neuron |
| *MYO16* | 0.00E+00 | 0.00E+00 | 2.85 | 0.90 | 0.43 | 35 | Glutamatergic Neuron |
| *TNC* | 1.83E-69 | 3.48E-65 | 2.81 | 0.39 | 0.07 | 35 | Glutamatergic Neuron |
| *LMO3* | 8.72E-127 | 1.66E-122 | 2.77 | 0.59 | 0.17 | 35 | Glutamatergic Neuron |
| *POU3F2* | 1.67E-72 | 3.18E-68 | 2.53 | 0.44 | 0.09 | 35 | Glutamatergic Neuron |
| *EPHA5* | 5.00E-219 | 9.51E-215 | 2.52 | 0.94 | 0.34 | 35 | Glutamatergic Neuron |
| *ZBTB18* | 4.85E-58 | 9.21E-54 | 2.47 | 0.37 | 0.07 | 35 | Glutamatergic Neuron |
| *PDZD2* | 2.69E-62 | 5.12E-58 | 2.44 | 0.39 | 0.11 | 35 | Glutamatergic Neuron |
| *HS3ST5* | 5.80E-91 | 1.10E-86 | 2.42 | 0.58 | 0.17 | 35 | Glutamatergic Neuron |
| *SLC17A6* | 6.60E-62 | 1.25E-57 | 2.37 | 0.42 | 0.10 | 35 | Glutamatergic Neuron |
| *KCNQ3* | 4.68E-85 | 8.90E-81 | 2.32 | 0.57 | 0.19 | 35 | Glutamatergic Neuron |
| *SDCCAG8* | 1.34E-100 | 2.55E-96 | 2.32 | 0.53 | 0.18 | 35 | Glutamatergic Neuron |
| *TRPM3* | 1.13E-114 | 2.14E-110 | 2.25 | 0.62 | 0.26 | 35 | Glutamatergic Neuron |
| *WNT7B* | 0.00E+00 | 0.00E+00 | 5.94 | 0.83 | 0.04 | 36 | Astrocyte |
| *WIF1* | 3.17E-127 | 6.03E-123 | 5.00 | 0.33 | 0.02 | 36 | Astrocyte |
| *LOC115494438* | 8.68E-146 | 1.65E-141 | 4.59 | 0.41 | 0.02 | 36 | Astrocyte |
| *LOC115497284* | 1.02E-101 | 1.94E-97 | 4.26 | 0.33 | 0.03 | 36 | Astrocyte |
| *LOC121470332* | 2.97E-146 | 5.65E-142 | 3.99 | 0.49 | 0.04 | 36 | Astrocyte |
| *NKX2-4* | 2.18E-119 | 4.14E-115 | 3.93 | 0.40 | 0.03 | 36 | Astrocyte |
| *RASGEF1B* | 1.18E-127 | 2.24E-123 | 3.90 | 0.43 | 0.03 | 36 | Astrocyte |
| *SLC15A2* | 3.26E-202 | 6.20E-198 | 3.59 | 0.71 | 0.09 | 36 | Astrocyte |
| *EMX2* | 1.70E-85 | 3.24E-81 | 3.46 | 0.34 | 0.03 | 36 | Astrocyte |
| *RIPOR2* | 1.99E-217 | 3.78E-213 | 3.26 | 0.70 | 0.13 | 36 | Astrocyte |
| *ARAP2* | 1.69E-95 | 3.21E-91 | 3.21 | 0.44 | 0.06 | 36 | Astrocyte |
| *FGFRL1* | 4.90E-233 | 9.31E-229 | 2.98 | 0.91 | 0.21 | 36 | Astrocyte |
| *SLIT2* | 2.90E-259 | 5.51E-255 | 2.88 | 0.97 | 0.44 | 36 | Astrocyte |
| *MTHFD1L* | 2.14E-69 | 4.06E-65 | 2.80 | 0.37 | 0.07 | 36 | Astrocyte |

| Gene | P-Value | FDR | Average $log_2FC$ | Fraction 1 | Fraction 2 | Cluster | Annotation |
|---|---|---|---|---|---|---|---|
| ***LOC105759165*** | 7.57E-74 | 1.44E-69 | 2.79 | 0.40 | 0.07 | 36 | Astrocyte |
| ***COL14A1*** | 1.42E-62 | 2.69E-58 | 2.78 | 0.36 | 0.06 | 36 | Astrocyte |
| ***WWC1*** | 3.73E-94 | 7.09E-90 | 2.74 | 0.44 | 0.09 | 36 | Astrocyte |
| ***TNFRSF19*** | 1.05E-95 | 2.00E-91 | 2.73 | 0.50 | 0.10 | 36 | Astrocyte |
| ***LRP2*** | 1.17E-94 | 2.23E-90 | 2.69 | 0.51 | 0.09 | 36 | Astrocyte |
| ***RGS6*** | 3.23E-174 | 6.14E-170 | 2.66 | 0.80 | 0.22 | 36 | Astrocyte |
| ***LOC116808418*** | 1.58E-223 | 3.01E-219 | 10.30 | 0.31 | 0.00 | 37 | Macrophage |
| ***CSF1R*** | 0.00E+00 | 0.00E+00 | 9.97 | 0.55 | 0.00 | 37 | Macrophage |
| ***SPI1*** | 0.00E+00 | 0.00E+00 | 9.80 | 0.48 | 0.00 | 37 | Macrophage |
| ***CD2*** | 2.14E-252 | 4.07E-248 | 9.60 | 0.37 | 0.00 | 37 | Macrophage |
| ***LOC100225774*** | 0.00E+00 | 0.00E+00 | 9.24 | 0.58 | 0.00 | 37 | Macrophage |
| ***FYB1*** | 3.28E-189 | 6.24E-185 | 9.23 | 0.28 | 0.00 | 37 | Macrophage |
| ***LOC100226241*** | 4.75E-195 | 9.03E-191 | 9.07 | 0.30 | 0.00 | 37 | Macrophage |
| ***CX3CR1*** | 1.75E-178 | 3.32E-174 | 9.00 | 0.27 | 0.00 | 37 | Macrophage |
| ***PLD4*** | 6.29E-188 | 1.20E-183 | 8.51 | 0.30 | 0.00 | 37 | Macrophage |
| ***TFEC*** | 8.74E-214 | 1.66E-209 | 7.79 | 0.36 | 0.00 | 37 | Macrophage |
| ***LOC115497648*** | 4.43E-196 | 8.43E-192 | 7.56 | 0.34 | 0.00 | 37 | Macrophage |
| ***INPP5D*** | 0.00E+00 | 0.00E+00 | 7.52 | 0.63 | 0.01 | 37 | Macrophage |
| ***IKZF1*** | 0.00E+00 | 0.00E+00 | 7.45 | 0.57 | 0.01 | 37 | Macrophage |
| ***LOC101232863*** | 1.28E-177 | 2.43E-173 | 7.37 | 0.32 | 0.00 | 37 | Macrophage |
| ***LYN*** | 0.00E+00 | 0.00E+00 | 7.35 | 0.63 | 0.01 | 37 | Macrophage |
| ***IRF8*** | 1.70E-152 | 3.23E-148 | 7.30 | 0.27 | 0.00 | 37 | Macrophage |
| ***LCP1*** | 2.20E-262 | 4.19E-258 | 7.21 | 0.46 | 0.01 | 37 | Macrophage |
| ***STAB1*** | 0.00E+00 | 0.00E+00 | 7.19 | 0.67 | 0.01 | 37 | Macrophage |
| ***LY86*** | 4.24E-175 | 8.05E-171 | 7.15 | 0.32 | 0.00 | 37 | Macrophage |
| ***RBM47*** | 5.53E-217 | 1.05E-212 | 7.09 | 0.40 | 0.01 | 37 | Macrophage |
| ***CDH5*** | 0.00E+00 | 0.00E+00 | 7.36 | 0.78 | 0.01 | 38 | Capillary Endothelial |

| Gene | P-Value | FDR | Average $log_2FC$ | Fraction 1 | Fraction 2 | Cluster | Annotation |
|---|---|---|---|---|---|---|---|
| ***RASGRP3*** | 0.00E+00 | 0.00E+00 | 7.26 | 0.67 | 0.01 | 38 | Capillary Endothelial |
| ***ADGRF5*** | 2.68E-193 | 5.10E-189 | 7.19 | 0.36 | 0.00 | 38 | Capillary Endothelial |
| ***TEK*** | 1.43E-160 | 2.73E-156 | 7.14 | 0.30 | 0.00 | 38 | Capillary Endothelial |
| ***EGFL7*** | 2.20E-300 | 4.18E-296 | 7.11 | 0.55 | 0.01 | 38 | Capillary Endothelial |
| ***P2RY8*** | 4.43E-253 | 8.43E-249 | 7.06 | 0.48 | 0.01 | 38 | Capillary Endothelial |
| ***ADGRL4*** | 0.00E+00 | 0.00E+00 | 6.95 | 0.75 | 0.01 | 38 | Capillary Endothelial |
| ***ZNF366*** | 0.00E+00 | 0.00E+00 | 6.93 | 0.63 | 0.01 | 38 | Capillary Endothelial |
| ***TM4SF18*** | 4.69E-171 | 8.91E-167 | 6.93 | 0.33 | 0.00 | 38 | Capillary Endothelial |
| ***VWF*** | 5.22E-272 | 9.93E-268 | 6.70 | 0.53 | 0.01 | 38 | Capillary Endothelial |
| ***MYCT1*** | 3.28E-147 | 6.24E-143 | 6.68 | 0.29 | 0.00 | 38 | Capillary Endothelial |
| ***FOXQ1*** | 3.89E-143 | 7.39E-139 | 6.62 | 0.29 | 0.00 | 38 | Capillary Endothelial |
| ***FLI1*** | 0.00E+00 | 0.00E+00 | 6.61 | 0.68 | 0.01 | 38 | Capillary Endothelial |
| ***KDR*** | 2.80E-312 | 5.33E-308 | 6.58 | 0.61 | 0.01 | 38 | Capillary Endothelial |
| ***PCDH12*** | 6.70E-134 | 1.27E-129 | 6.58 | 0.27 | 0.00 | 38 | Capillary Endothelial |
| ***LOC101233446*** | 5.96E-220 | 1.13E-215 | 6.57 | 0.44 | 0.01 | 38 | Capillary Endothelial |
| ***ERG*** | 1.15E-285 | 2.19E-281 | 6.51 | 0.57 | 0.01 | 38 | Capillary Endothelial |
| ***FLT1*** | 0.00E+00 | 0.00E+00 | 6.48 | 0.92 | 0.03 | 38 | Capillary Endothelial |
| ***LOC100229347*** | 0.00E+00 | 0.00E+00 | 6.34 | 0.80 | 0.02 | 38 | Capillary Endothelial |
| ***FOXF2*** | 2.47E-181 | 4.70E-177 | 6.34 | 0.38 | 0.01 | 38 | Capillary Endothelial |
| ***ESR2*** | 7.60E-105 | 1.45E-100 | 4.56 | 0.36 | 0.02 | 39 | Glutamatergic Neuron |
| ***LOC116808227*** | 1.60E-77 | 3.04E-73 | 4.31 | 0.28 | 0.02 | 39 | Glutamatergic Neuron |
| ***POU6F2*** | 4.41E-235 | 8.39E-231 | 4.03 | 0.81 | 0.08 | 39 | Glutamatergic Neuron |
| ***GREB1*** | 1.76E-101 | 3.35E-97 | 3.68 | 0.46 | 0.05 | 39 | Glutamatergic Neuron |
| ***LOC115494974*** | 1.45E-62 | 2.75E-58 | 3.08 | 0.34 | 0.04 | 39 | Glutamatergic Neuron |
| ***ADCYAP1*** | 6.69E-142 | 1.27E-137 | 3.07 | 0.68 | 0.08 | 39 | Glutamatergic Neuron |
| ***GABRA5*** | 3.42E-51 | 6.50E-47 | 2.97 | 0.30 | 0.04 | 39 | Glutamatergic Neuron |
| ***NPAS2*** | 2.16E-115 | 4.11E-111 | 2.93 | 0.63 | 0.10 | 39 | Glutamatergic Neuron |

| Gene | P-Value | FDR | Average $log_2FC$ | Fraction 1 | Fraction 2 | Cluster | Annotation |
|---|---|---|---|---|---|---|---|
| *LOC115494899* | 3.46E-89 | 6.59E-85 | 2.76 | 0.54 | 0.09 | 39 | Glutamatergic Neuron |
| *EYS* | 8.39E-170 | 1.59E-165 | 2.71 | 0.88 | 0.21 | 39 | Glutamatergic Neuron |
| *HS3ST5* | 1.16E-112 | 2.20E-108 | 2.57 | 0.73 | 0.17 | 39 | Glutamatergic Neuron |
| *GABRB3* | 5.41E-105 | 1.03E-100 | 2.44 | 0.70 | 0.20 | 39 | Glutamatergic Neuron |
| *ZBTB7C* | 3.35E-42 | 6.36E-38 | 2.40 | 0.33 | 0.07 | 39 | Glutamatergic Neuron |
| *COL25A1* | 3.36E-85 | 6.40E-81 | 2.40 | 0.61 | 0.13 | 39 | Glutamatergic Neuron |
| *ALK* | 4.97E-36 | 9.44E-32 | 2.29 | 0.34 | 0.09 | 39 | Glutamatergic Neuron |
| *LRP1B* | 1.89E-107 | 3.59E-103 | 2.25 | 0.84 | 0.36 | 39 | Glutamatergic Neuron |
| *LOC116807738* | 1.62E-56 | 3.09E-52 | 2.24 | 0.46 | 0.10 | 39 | Glutamatergic Neuron |
| *TAFA2* | 3.56E-89 | 6.76E-85 | 2.17 | 0.72 | 0.20 | 39 | Glutamatergic Neuron |
| *TMEM163* | 1.70E-52 | 3.24E-48 | 2.15 | 0.45 | 0.10 | 39 | Glutamatergic Neuron |
| *LOC100221646* | 9.18E-96 | 1.75E-91 | 2.11 | 0.75 | 0.21 | 39 | Glutamatergic Neuron |
| *NRN1* | 0.00E+00 | 0.00E+00 | 3.39 | 0.44 | 0.05 | 4 | Glutamatergic Neuron |
| *ADCYAP1* | 0.00E+00 | 0.00E+00 | 2.44 | 0.39 | 0.07 | 4 | Glutamatergic Neuron |
| *SLC17A6* | 0.00E+00 | 0.00E+00 | 2.23 | 0.41 | 0.08 | 4 | Glutamatergic Neuron |
| *TMEM163* | 0.00E+00 | 0.00E+00 | 2.06 | 0.37 | 0.09 | 4 | Glutamatergic Neuron |
| *CDH18* | 0.00E+00 | 0.00E+00 | 2.06 | 0.59 | 0.18 | 4 | Glutamatergic Neuron |
| *TAFA1* | 0.00E+00 | 0.00E+00 | 1.94 | 0.45 | 0.14 | 4 | Glutamatergic Neuron |
| *MDGA1* | 2.01E-320 | 3.81E-316 | 1.85 | 0.47 | 0.16 | 4 | Glutamatergic Neuron |
| *CDH12* | 0.00E+00 | 0.00E+00 | 1.84 | 0.65 | 0.26 | 4 | Glutamatergic Neuron |
| *CA10* | 0.00E+00 | 0.00E+00 | 1.80 | 0.48 | 0.16 | 4 | Glutamatergic Neuron |
| *GRM1* | 0.00E+00 | 0.00E+00 | 1.74 | 0.47 | 0.15 | 4 | Glutamatergic Neuron |
| *RYR2* | 0.00E+00 | 0.00E+00 | 1.70 | 0.67 | 0.25 | 4 | Glutamatergic Neuron |
| *MGAT4C* | 0.00E+00 | 0.00E+00 | 1.68 | 0.61 | 0.22 | 4 | Glutamatergic Neuron |
| *TENM1* | 0.00E+00 | 0.00E+00 | 1.61 | 0.83 | 0.42 | 4 | Glutamatergic Neuron |
| *KLHL1* | 1.74E-238 | 3.31E-234 | 1.61 | 0.41 | 0.14 | 4 | Glutamatergic Neuron |
| *LOC115496780* | 4.65E-254 | 8.83E-250 | 1.57 | 0.43 | 0.15 | 4 | Glutamatergic Neuron |

| Gene | P-Value | FDR | Average $log_2FC$ | Fraction 1 | Fraction 2 | Cluster | Annotation |
|---|---|---|---|---|---|---|---|
| *GLRA2* | 2.13E-238 | 4.04E-234 | 1.54 | 0.44 | 0.16 | 4 | Glutamatergic Neuron |
| *LOC100221630* | 2.28E-299 | 4.33E-295 | 1.52 | 0.47 | 0.16 | 4 | Glutamatergic Neuron |
| *LRP1B* | 0.00E+00 | 0.00E+00 | 1.46 | 0.74 | 0.35 | 4 | Glutamatergic Neuron |
| *GRM8* | 1.03E-301 | 1.96E-297 | 1.42 | 0.52 | 0.20 | 4 | Glutamatergic Neuron |
| *TAFA2* | 8.37E-252 | 1.59E-247 | 1.41 | 0.48 | 0.19 | 4 | Glutamatergic Neuron |
| *MATN1* | 2.74E-226 | 5.21E-222 | 8.56 | 0.39 | 0.01 | 40 | Endothelial Cell |
| *HAPLN1* | 2.45E-148 | 4.65E-144 | 7.69 | 0.31 | 0.00 | 40 | Endothelial Cell |
| *COL9A3* | 2.07E-264 | 3.94E-260 | 7.00 | 0.39 | 0.01 | 40 | Endothelial Cell |
| *COL2A1* | 0.00E+00 | 0.00E+00 | 6.71 | 0.45 | 0.03 | 40 | Endothelial Cell |
| *COL9A1* | 0.00E+00 | 0.00E+00 | 6.43 | 0.41 | 0.03 | 40 | Endothelial Cell |
| *EYA1* | 1.67E-172 | 3.18E-168 | 6.03 | 0.44 | 0.01 | 40 | Endothelial Cell |
| *COL11A1* | 0.00E+00 | 0.00E+00 | 6.01 | 0.57 | 0.15 | 40 | Endothelial Cell |
| *ACAN* | 2.23E-191 | 4.25E-187 | 5.51 | 0.35 | 0.03 | 40 | Endothelial Cell |
| *ZNF385B* | 1.86E-257 | 3.55E-253 | 5.33 | 0.47 | 0.04 | 40 | Endothelial Cell |
| *COL1A2* | 2.60E-140 | 4.93E-136 | 5.14 | 0.39 | 0.03 | 40 | Endothelial Cell |
| *COL27A1* | 0.00E+00 | 0.00E+00 | 4.64 | 0.45 | 0.11 | 40 | Endothelial Cell |
| *PIEZO2* | 2.53E-88 | 4.82E-84 | 3.60 | 0.35 | 0.07 | 40 | Endothelial Cell |
| *UGDH* | 5.24E-175 | 9.96E-171 | 3.45 | 0.33 | 0.07 | 40 | Endothelial Cell |
| *BNC2* | 4.43E-62 | 8.42E-58 | 3.05 | 0.42 | 0.07 | 40 | Endothelial Cell |
| *KCNQ5* | 1.17E-74 | 2.22E-70 | 2.51 | 0.64 | 0.21 | 40 | Endothelial Cell |
| *FGFR2* | 2.31E-60 | 4.39E-56 | 2.36 | 0.49 | 0.18 | 40 | Endothelial Cell |
| *THSD4* | 3.01E-53 | 5.73E-49 | 2.21 | 0.47 | 0.17 | 40 | Endothelial Cell |
| *RBMS1* | 3.68E-31 | 6.99E-27 | 1.87 | 0.42 | 0.15 | 40 | Endothelial Cell |
| *FNDC3B* | 2.96E-56 | 5.62E-52 | 1.86 | 0.58 | 0.28 | 40 | Endothelial Cell |
| *TCF7L2* | 5.39E-32 | 1.03E-27 | 1.47 | 0.60 | 0.27 | 40 | Endothelial Cell |
| *ANLN* | 2.77E-117 | 5.26E-113 | 4.32 | 0.46 | 0.02 | 41 | Astrocyte |
| *KIF2C* | 6.29E-66 | 1.20E-61 | 4.14 | 0.29 | 0.02 | 41 | Astrocyte |

| Gene | P-Value | FDR | Average $log_2FC$ | Fraction 1 | Fraction 2 | Cluster | Annotation |
|---|---|---|---|---|---|---|---|
| *TPX2* | 3.60E-119 | 6.84E-115 | 4.12 | 0.49 | 0.03 | 41 | Astrocyte |
| *DEPDC1B* | 5.13E-81 | 9.76E-77 | 4.08 | 0.35 | 0.02 | 41 | Astrocyte |
| *CDCA2* | 4.52E-67 | 8.58E-63 | 4.07 | 0.29 | 0.02 | 41 | Astrocyte |
| *CENPF* | 7.80E-111 | 1.48E-106 | 3.95 | 0.49 | 0.04 | 41 | Astrocyte |
| *CENPE* | 4.19E-127 | 7.97E-123 | 3.86 | 0.55 | 0.04 | 41 | Astrocyte |
| *NEK2* | 8.97E-83 | 1.71E-78 | 3.81 | 0.38 | 0.03 | 41 | Astrocyte |
| *NUSAP1* | 4.17E-113 | 7.92E-109 | 3.77 | 0.49 | 0.03 | 41 | Astrocyte |
| *SHCBP1* | 4.32E-86 | 8.22E-82 | 3.76 | 0.40 | 0.03 | 41 | Astrocyte |
| *TACC3* | 2.55E-69 | 4.84E-65 | 3.76 | 0.33 | 0.02 | 41 | Astrocyte |
| *ECT2* | 1.82E-60 | 3.46E-56 | 3.74 | 0.30 | 0.02 | 41 | Astrocyte |
| *PIMREG* | 7.97E-98 | 1.51E-93 | 3.73 | 0.46 | 0.04 | 41 | Astrocyte |
| *KIF20B* | 1.39E-74 | 2.64E-70 | 3.73 | 0.35 | 0.03 | 41 | Astrocyte |
| *TTK* | 1.52E-73 | 2.90E-69 | 3.61 | 0.35 | 0.03 | 41 | Astrocyte |
| *FOXM1* | 2.79E-76 | 5.30E-72 | 3.60 | 0.38 | 0.03 | 41 | Astrocyte |
| *TOP2A* | 1.60E-73 | 3.04E-69 | 3.59 | 0.37 | 0.03 | 41 | Astrocyte |
| *CDK1* | 4.57E-62 | 8.68E-58 | 3.55 | 0.31 | 0.03 | 41 | Astrocyte |
| *TERF1* | 6.27E-60 | 1.19E-55 | 3.54 | 0.30 | 0.02 | 41 | Astrocyte |
| *ASPM* | 6.01E-106 | 1.14E-101 | 3.54 | 0.50 | 0.04 | 41 | Astrocyte |
| *CBY2* | 6.54E-132 | 1.24E-127 | 7.99 | 0.29 | 0.00 | 42 | Vascular Associated Smooth Muscle Cell |
| *LOC115493878* | 7.34E-257 | 1.40E-252 | 7.71 | 0.57 | 0.00 | 42 | Vascular Associated Smooth Muscle Cell |
| *LOC116808812* | 4.33E-150 | 8.22E-146 | 7.69 | 0.35 | 0.00 | 42 | Vascular Associated Smooth Muscle Cell |
| *LOC115494366* | 1.75E-131 | 3.33E-127 | 7.54 | 0.31 | 0.00 | 42 | Vascular Associated Smooth Muscle Cell |
| *LOC100229617* | 1.03E-238 | 1.96E-234 | 7.25 | 0.56 | 0.01 | 42 | Vascular Associated Smooth Muscle Cell |
| *TBX18* | 0.00E+00 | 0.00E+00 | 7.24 | 0.79 | 0.01 | 42 | Vascular Associated Smooth Muscle Cell |
| *SPRY3* | 5.11E-105 | 9.72E-101 | 7.11 | 0.26 | 0.00 | 42 | Vascular Associated Smooth Muscle Cell |
| *LOC100226679* | 1.72E-207 | 3.27E-203 | 6.81 | 0.52 | 0.01 | 42 | Vascular Associated Smooth Muscle Cell |
| *ITGA4* | 8.74E-218 | 1.66E-213 | 6.52 | 0.55 | 0.01 | 42 | Vascular Associated Smooth Muscle Cell |

| Gene | P-Value | FDR | Average $log_2FC$ | Fraction 1 | Fraction 2 | Cluster | Annotation |
|---|---|---|---|---|---|---|---|
| *FN1* | 0.00E+00 | 0.00E+00 | 6.46 | 0.80 | 0.02 | 42 | Vascular Associated Smooth Muscle Cell |
| *ANGPT4* | 1.83E-111 | 3.47E-107 | 6.36 | 0.30 | 0.01 | 42 | Vascular Associated Smooth Muscle Cell |
| *S1PR3* | 1.35E-127 | 2.56E-123 | 6.35 | 0.35 | 0.01 | 42 | Vascular Associated Smooth Muscle Cell |
| *TRPC6* | 1.08E-309 | 2.05E-305 | 6.35 | 0.73 | 0.02 | 42 | Vascular Associated Smooth Muscle Cell |
| *COLEC10* | 8.73E-124 | 1.66E-119 | 6.16 | 0.34 | 0.01 | 42 | Vascular Associated Smooth Muscle Cell |
| *LRRC17* | 9.88E-209 | 1.88E-204 | 6.05 | 0.59 | 0.01 | 42 | Vascular Associated Smooth Muscle Cell |
| *GGT5* | 2.75E-199 | 5.23E-195 | 6.04 | 0.56 | 0.01 | 42 | Vascular Associated Smooth Muscle Cell |
| *TBX2* | 2.09E-158 | 3.97E-154 | 6.03 | 0.46 | 0.01 | 42 | Vascular Associated Smooth Muscle Cell |
| *PCDH18* | 4.17E-248 | 7.92E-244 | 5.99 | 0.66 | 0.02 | 42 | Vascular Associated Smooth Muscle Cell |
| *NID1* | 2.38E-309 | 4.52E-305 | 5.79 | 0.79 | 0.03 | 42 | Vascular Associated Smooth Muscle Cell |
| *OLFML3* | 2.08E-129 | 3.95E-125 | 5.78 | 0.39 | 0.01 | 42 | Vascular Associated Smooth Muscle Cell |
| *ADGRF5* | 1.85E-178 | 3.52E-174 | 7.26 | 0.48 | 0.00 | 43 | Endothelial Cell |
| *ZNF366* | 2.01E-313 | 3.82E-309 | 7.15 | 0.81 | 0.01 | 43 | Endothelial Cell |
| *CD93* | 2.44E-192 | 4.63E-188 | 7.07 | 0.52 | 0.00 | 43 | Endothelial Cell |
| *LOC100224802* | 4.90E-102 | 9.31E-98 | 7.05 | 0.29 | 0.00 | 43 | Endothelial Cell |
| *VWF* | 4.62E-264 | 8.79E-260 | 7.05 | 0.71 | 0.01 | 43 | Endothelial Cell |
| *CDH5* | 0.00E+00 | 0.00E+00 | 6.98 | 0.88 | 0.01 | 43 | Endothelial Cell |
| *LOC115493878* | 5.13E-197 | 9.74E-193 | 6.92 | 0.54 | 0.00 | 43 | Endothelial Cell |
| *LOC100229347* | 0.00E+00 | 0.00E+00 | 6.92 | 0.96 | 0.02 | 43 | Endothelial Cell |
| *LOC101233446* | 1.53E-238 | 2.92E-234 | 6.90 | 0.66 | 0.01 | 43 | Endothelial Cell |
| *LOC100222941* | 0.00E+00 | 0.00E+00 | 6.87 | 0.91 | 0.02 | 43 | Endothelial Cell |
| *ACVRL1* | 9.63E-188 | 1.83E-183 | 6.81 | 0.53 | 0.01 | 43 | Endothelial Cell |
| *LOC101233598* | 2.90E-94 | 5.51E-90 | 6.80 | 0.28 | 0.00 | 43 | Endothelial Cell |
| *FN1* | 0.00E+00 | 0.00E+00 | 6.78 | 0.97 | 0.02 | 43 | Endothelial Cell |
| *P2RY8* | 2.15E-210 | 4.09E-206 | 6.78 | 0.59 | 0.01 | 43 | Endothelial Cell |
| *LOC116808812* | 2.88E-130 | 5.48E-126 | 6.77 | 0.37 | 0.00 | 43 | Endothelial Cell |
| *ALPL* | 0.00E+00 | 0.00E+00 | 6.75 | 0.88 | 0.02 | 43 | Endothelial Cell |

| Gene | P-Value | FDR | Average $log_2FC$ | Fraction 1 | Fraction 2 | Cluster | Annotation |
|---|---|---|---|---|---|---|---|
| *TEK* | 1.16E-115 | 2.20E-111 | 6.74 | 0.34 | 0.00 | 43 | Endothelial Cell |
| *RASGRP3* | 3.76E-267 | 7.14E-263 | 6.73 | 0.75 | 0.01 | 43 | Endothelial Cell |
| *LOC100224290* | 5.17E-169 | 9.84E-165 | 6.72 | 0.50 | 0.01 | 43 | Endothelial Cell |
| *EGFL7* | 2.79E-213 | 5.31E-209 | 6.71 | 0.61 | 0.01 | 43 | Endothelial Cell |
| *LOC121469413* | 8.89E-62 | 1.69E-57 | 3.81 | 0.39 | 0.04 | 44 | GABAergic Neuron |
| *ADARB2* | 7.38E-194 | 1.40E-189 | 3.74 | 0.97 | 0.24 | 44 | GABAergic Neuron |
| *EDIL3* | 1.01E-137 | 1.92E-133 | 2.84 | 0.90 | 0.31 | 44 | GABAergic Neuron |
| *EPHA5* | 2.91E-128 | 5.53E-124 | 2.60 | 0.96 | 0.34 | 44 | GABAergic Neuron |
| *ERBB4* | 4.61E-136 | 8.77E-132 | 2.56 | 0.96 | 0.45 | 44 | GABAergic Neuron |
| *XKR4* | 3.81E-88 | 7.24E-84 | 2.30 | 0.80 | 0.34 | 44 | GABAergic Neuron |
| *PDZRN3* | 1.55E-48 | 2.95E-44 | 2.29 | 0.59 | 0.19 | 44 | GABAergic Neuron |
| *TCF12* | 4.75E-152 | 9.03E-148 | 2.28 | 0.88 | 0.48 | 44 | GABAergic Neuron |
| *SH3RF3* | 5.28E-134 | 1.00E-129 | 2.16 | 0.97 | 0.57 | 44 | GABAergic Neuron |
| *OLFM3* | 4.87E-60 | 9.26E-56 | 2.11 | 0.55 | 0.25 | 44 | GABAergic Neuron |
| *DOCK11* | 1.40E-21 | 2.65E-17 | 1.99 | 0.36 | 0.11 | 44 | GABAergic Neuron |
| *EML5* | 7.20E-58 | 1.37E-53 | 1.97 | 0.54 | 0.25 | 44 | GABAergic Neuron |
| *GRIK2* | 3.32E-92 | 6.31E-88 | 1.95 | 0.97 | 0.51 | 44 | GABAergic Neuron |
| *KCNIP1* | 3.15E-90 | 5.98E-86 | 1.90 | 0.87 | 0.50 | 44 | GABAergic Neuron |
| *PROX1* | 2.36E-25 | 4.48E-21 | 1.85 | 0.44 | 0.13 | 44 | GABAergic Neuron |
| *NFIB* | 5.86E-70 | 1.11E-65 | 1.85 | 0.90 | 0.43 | 44 | GABAergic Neuron |
| *EPHA7* | 7.43E-35 | 1.41E-30 | 1.83 | 0.59 | 0.27 | 44 | GABAergic Neuron |
| *MTUS2* | 3.54E-22 | 6.74E-18 | 1.81 | 0.42 | 0.13 | 44 | GABAergic Neuron |
| *ARX* | 2.24E-30 | 4.26E-26 | 1.77 | 0.59 | 0.22 | 44 | GABAergic Neuron |
| *LOC116808643* | 7.95E-35 | 1.51E-30 | 1.75 | 0.58 | 0.27 | 44 | GABAergic Neuron |
| *NPVF* | 1.88E-297 | 3.57E-293 | 7.16 | 0.32 | 0.03 | 45 | Glutamatergic Neuron |
| *LOC100228479* | 3.65E-99 | 6.93E-95 | 6.82 | 0.29 | 0.01 | 45 | Glutamatergic Neuron |
| *ARHGEF28* | 5.45E-166 | 1.04E-161 | 5.39 | 0.67 | 0.03 | 45 | Glutamatergic Neuron |

| Gene | P-Value | FDR | Average $log_2FC$ | Fraction 1 | Fraction 2 | Cluster | Annotation |
|---|---|---|---|---|---|---|---|
| ***BMP6*** | 3.55E-68 | 6.76E-64 | 5.33 | 0.28 | 0.01 | 45 | Glutamatergic Neuron |
| ***LOC121470336*** | 4.17E-48 | 7.92E-44 | 4.49 | 0.27 | 0.01 | 45 | Glutamatergic Neuron |
| ***RFX4*** | 1.86E-229 | 3.53E-225 | 4.31 | 0.86 | 0.12 | 45 | Glutamatergic Neuron |
| ***LHX9*** | 1.52E-84 | 2.90E-80 | 4.24 | 0.49 | 0.03 | 45 | Glutamatergic Neuron |
| ***GLIS1*** | 1.03E-84 | 1.96E-80 | 3.38 | 0.63 | 0.08 | 45 | Glutamatergic Neuron |
| ***AKAP12*** | 1.41E-84 | 2.67E-80 | 3.37 | 0.57 | 0.09 | 45 | Glutamatergic Neuron |
| ***ADAMTS2*** | 1.05E-33 | 2.00E-29 | 3.21 | 0.29 | 0.04 | 45 | Glutamatergic Neuron |
| ***SYTL5*** | 9.41E-57 | 1.79E-52 | 2.97 | 0.52 | 0.08 | 45 | Glutamatergic Neuron |
| ***KLHL1*** | 8.72E-79 | 1.66E-74 | 2.86 | 0.73 | 0.16 | 45 | Glutamatergic Neuron |
| ***MTHFD1L*** | 1.90E-46 | 3.60E-42 | 2.75 | 0.47 | 0.08 | 45 | Glutamatergic Neuron |
| ***MTUS2*** | 5.70E-52 | 1.08E-47 | 2.75 | 0.56 | 0.13 | 45 | Glutamatergic Neuron |
| ***LHX2*** | 1.21E-37 | 2.31E-33 | 2.71 | 0.41 | 0.08 | 45 | Glutamatergic Neuron |
| ***CDH12*** | 8.68E-106 | 1.65E-101 | 2.61 | 0.93 | 0.27 | 45 | Glutamatergic Neuron |
| ***NPAS2*** | 2.80E-44 | 5.32E-40 | 2.49 | 0.51 | 0.11 | 45 | Glutamatergic Neuron |
| ***PCSK2*** | 3.89E-48 | 7.39E-44 | 2.36 | 0.58 | 0.16 | 45 | Glutamatergic Neuron |
| ***SCG2*** | 2.04E-69 | 3.89E-65 | 2.35 | 0.77 | 0.22 | 45 | Glutamatergic Neuron |
| ***IL1RAPL1*** | 1.53E-90 | 2.91E-86 | 2.34 | 0.89 | 0.34 | 45 | Glutamatergic Neuron |
| ***NEUROD6*** | 3.33E-57 | 6.33E-53 | 5.57 | 0.33 | 0.01 | 46 | Glutamatergic Neuron |
| ***NHLH2*** | 6.79E-107 | 1.29E-102 | 4.59 | 0.65 | 0.06 | 46 | Glutamatergic Neuron |
| ***CCK*** | 1.19E-53 | 2.26E-49 | 4.26 | 0.33 | 0.04 | 46 | Glutamatergic Neuron |
| ***NRN1*** | 8.70E-83 | 1.65E-78 | 4.07 | 0.60 | 0.06 | 46 | Glutamatergic Neuron |
| ***PTPN14*** | 6.05E-39 | 1.15E-34 | 3.43 | 0.37 | 0.06 | 46 | Glutamatergic Neuron |
| ***STXBP5L*** | 9.69E-132 | 1.84E-127 | 3.23 | 0.83 | 0.23 | 46 | Glutamatergic Neuron |
| ***LOC100218075*** | 1.25E-24 | 2.37E-20 | 2.92 | 0.33 | 0.06 | 46 | Glutamatergic Neuron |
| ***LOC100223021*** | 3.19E-22 | 6.07E-18 | 2.81 | 0.33 | 0.06 | 46 | Glutamatergic Neuron |
| ***SLC17A6*** | 6.96E-35 | 1.32E-30 | 2.74 | 0.44 | 0.10 | 46 | Glutamatergic Neuron |
| ***DPF3*** | 5.64E-63 | 1.07E-58 | 2.73 | 0.66 | 0.19 | 46 | Glutamatergic Neuron |

| Gene | P-Value | FDR | Average $log_2FC$ | Fraction 1 | Fraction 2 | Cluster | Annotation |
|---|---|---|---|---|---|---|---|
| ***ZBTB18*** | 1.58E-25 | 3.00E-21 | 2.70 | 0.36 | 0.07 | 46 | Glutamatergic Neuron |
| ***NRP2*** | 5.05E-44 | 9.61E-40 | 2.58 | 0.58 | 0.18 | 46 | Glutamatergic Neuron |
| ***NFIC*** | 9.99E-22 | 1.90E-17 | 2.44 | 0.38 | 0.08 | 46 | Glutamatergic Neuron |
| ***CRB1*** | 3.06E-28 | 5.81E-24 | 2.42 | 0.38 | 0.13 | 46 | Glutamatergic Neuron |
| ***EBF3*** | 4.91E-27 | 9.34E-23 | 2.42 | 0.36 | 0.05 | 46 | Glutamatergic Neuron |
| ***CDKN1C*** | 1.25E-19 | 2.37E-15 | 2.41 | 0.35 | 0.08 | 46 | Glutamatergic Neuron |
| ***EBF1*** | 1.78E-49 | 3.38E-45 | 2.08 | 0.62 | 0.10 | 46 | Glutamatergic Neuron |
| ***LRP8*** | 3.71E-30 | 7.06E-26 | 2.07 | 0.55 | 0.27 | 46 | Glutamatergic Neuron |
| ***PPP2R2C*** | 4.00E-56 | 7.60E-52 | 2.04 | 0.79 | 0.42 | 46 | Glutamatergic Neuron |
| ***FSTL5*** | 1.11E-39 | 2.10E-35 | 2.00 | 0.76 | 0.39 | 46 | Glutamatergic Neuron |
| ***PENK*** | 4.50E-186 | 8.55E-182 | 4.22 | 0.74 | 0.25 | 47 | Glutamatergic Neuron |
| ***LOC121469413*** | 3.70E-57 | 7.03E-53 | 3.87 | 0.52 | 0.04 | 47 | Glutamatergic Neuron |
| ***ADARB2*** | 2.02E-123 | 3.84E-119 | 3.76 | 0.97 | 0.24 | 47 | Glutamatergic Neuron |
| ***GRIP2*** | 1.55E-154 | 2.94E-150 | 3.69 | 0.93 | 0.21 | 47 | Glutamatergic Neuron |
| ***AMIGO2*** | 8.05E-39 | 1.53E-34 | 3.63 | 0.40 | 0.04 | 47 | Glutamatergic Neuron |
| ***GRIK3*** | 3.70E-167 | 7.04E-163 | 3.52 | 0.91 | 0.19 | 47 | Glutamatergic Neuron |
| ***NETO2*** | 5.02E-114 | 9.55E-110 | 3.46 | 0.77 | 0.15 | 47 | Glutamatergic Neuron |
| ***VWC2*** | 9.23E-55 | 1.76E-50 | 3.22 | 0.63 | 0.08 | 47 | Glutamatergic Neuron |
| ***GRIN3A*** | 2.06E-34 | 3.91E-30 | 3.11 | 0.43 | 0.07 | 47 | Glutamatergic Neuron |
| ***SCUBE1*** | 5.20E-54 | 9.89E-50 | 2.88 | 0.70 | 0.15 | 47 | Glutamatergic Neuron |
| ***MYO16*** | 1.54E-118 | 2.92E-114 | 2.80 | 0.91 | 0.43 | 47 | Glutamatergic Neuron |
| ***NETO1*** | 1.54E-59 | 2.92E-55 | 2.78 | 0.77 | 0.17 | 47 | Glutamatergic Neuron |
| ***LOC100227840*** | 7.85E-66 | 1.49E-61 | 2.70 | 0.85 | 0.21 | 47 | Glutamatergic Neuron |
| ***STK32B*** | 2.19E-45 | 4.16E-41 | 2.68 | 0.70 | 0.18 | 47 | Glutamatergic Neuron |
| ***SLCO3A1*** | 6.63E-33 | 1.26E-28 | 2.42 | 0.57 | 0.14 | 47 | Glutamatergic Neuron |
| ***BEND4*** | 6.05E-21 | 1.15E-16 | 2.38 | 0.37 | 0.07 | 47 | Glutamatergic Neuron |
| ***TMTC4*** | 6.95E-24 | 1.32E-19 | 2.36 | 0.47 | 0.12 | 47 | Glutamatergic Neuron |

| Gene | P-Value | FDR | Average $log_2FC$ | Fraction 1 | Fraction 2 | Cluster | Annotation |
|---|---|---|---|---|---|---|---|
| *LIN7A* | 2.95E-20 | 5.60E-16 | 2.31 | 0.38 | 0.08 | 47 | Glutamatergic Neuron |
| *LOC115494833* | 3.55E-23 | 6.76E-19 | 2.30 | 0.43 | 0.09 | 47 | Glutamatergic Neuron |
| *TMEM132D* | 1.17E-20 | 2.22E-16 | 2.29 | 0.40 | 0.09 | 47 | Glutamatergic Neuron |
| *MSX2* | 5.38E-96 | 1.02E-91 | 9.13 | 0.42 | 0.00 | 48 | Choroid Plexus Epithelial Cell |
| *SLC13A4* | 5.12E-200 | 9.72E-196 | 8.27 | 0.82 | 0.01 | 48 | Choroid Plexus Epithelial Cell |
| *TF* | 7.58E-305 | 1.44E-300 | 7.76 | 0.94 | 0.02 | 48 | Choroid Plexus Epithelial Cell |
| *SLC13A1* | 6.95E-74 | 1.32E-69 | 7.59 | 0.39 | 0.00 | 48 | Choroid Plexus Epithelial Cell |
| *TTR* | 0.00E+00 | 0.00E+00 | 7.56 | 0.98 | 0.21 | 48 | Choroid Plexus Epithelial Cell |
| *LOC115496261* | 6.76E-136 | 1.29E-131 | 7.33 | 0.67 | 0.01 | 48 | Choroid Plexus Epithelial Cell |
| *MSTN* | 1.98E-53 | 3.77E-49 | 7.31 | 0.28 | 0.00 | 48 | Choroid Plexus Epithelial Cell |
| *RARRES1* | 4.63E-94 | 8.79E-90 | 7.23 | 0.47 | 0.01 | 48 | Choroid Plexus Epithelial Cell |
| *LOC100226992* | 7.75E-57 | 1.47E-52 | 7.14 | 0.31 | 0.00 | 48 | Choroid Plexus Epithelial Cell |
| *LOC115495484* | 1.78E-51 | 3.39E-47 | 7.10 | 0.29 | 0.00 | 48 | Choroid Plexus Epithelial Cell |
| *LMX1A* | 4.52E-63 | 8.60E-59 | 6.86 | 0.36 | 0.00 | 48 | Choroid Plexus Epithelial Cell |
| *RBM47* | 5.03E-98 | 9.56E-94 | 6.79 | 0.56 | 0.01 | 48 | Choroid Plexus Epithelial Cell |
| *LOC115496260* | 2.40E-76 | 4.56E-72 | 6.73 | 0.45 | 0.01 | 48 | Choroid Plexus Epithelial Cell |
| *SLC16A12* | 1.45E-109 | 2.76E-105 | 6.55 | 0.64 | 0.01 | 48 | Choroid Plexus Epithelial Cell |
| *RSPO3* | 3.75E-114 | 7.13E-110 | 6.44 | 0.68 | 0.01 | 48 | Choroid Plexus Epithelial Cell |
| *LOC121471091* | 4.90E-122 | 9.31E-118 | 6.35 | 0.72 | 0.01 | 48 | Choroid Plexus Epithelial Cell |
| *ENPP2* | 3.31E-168 | 6.29E-164 | 6.22 | 0.82 | 0.03 | 48 | Choroid Plexus Epithelial Cell |
| *TRPM1* | 4.61E-120 | 8.77E-116 | 6.09 | 0.65 | 0.02 | 48 | Choroid Plexus Epithelial Cell |
| *TLL2* | 2.78E-110 | 5.29E-106 | 6.07 | 0.69 | 0.02 | 48 | Choroid Plexus Epithelial Cell |
| *LOC100229594* | 2.23E-84 | 4.24E-80 | 5.86 | 0.50 | 0.01 | 48 | Choroid Plexus Epithelial Cell |
| *NEK2* | 0.00E+00 | 0.00E+00 | 5.44 | 0.40 | 0.01 | 5 | Astrocyte |
| *ANLN* | 0.00E+00 | 0.00E+00 | 5.32 | 0.35 | 0.01 | 5 | Astrocyte |
| *NUSAP1* | 0.00E+00 | 0.00E+00 | 5.25 | 0.46 | 0.02 | 5 | Astrocyte |
| *TTK* | 0.00E+00 | 0.00E+00 | 5.25 | 0.37 | 0.01 | 5 | Astrocyte |

| Gene | P-Value | FDR | Average $log_2FC$ | Fraction 1 | Fraction 2 | Cluster | Annotation |
|---|---|---|---|---|---|---|---|
| *TTF2* | 0.00E+00 | 0.00E+00 | 5.22 | 0.59 | 0.03 | 5 | Astrocyte |
| *CENPF* | 0.00E+00 | 0.00E+00 | 5.10 | 0.47 | 0.02 | 5 | Astrocyte |
| *ASPM* | 0.00E+00 | 0.00E+00 | 5.09 | 0.48 | 0.02 | 5 | Astrocyte |
| *CENPE* | 0.00E+00 | 0.00E+00 | 5.08 | 0.49 | 0.02 | 5 | Astrocyte |
| *TOP2A* | 0.00E+00 | 0.00E+00 | 4.87 | 0.38 | 0.02 | 5 | Astrocyte |
| *TPX2* | 0.00E+00 | 0.00E+00 | 4.87 | 0.40 | 0.02 | 5 | Astrocyte |
| *KIF20B* | 0.00E+00 | 0.00E+00 | 4.79 | 0.33 | 0.01 | 5 | Astrocyte |
| *TACC3* | 0.00E+00 | 0.00E+00 | 4.68 | 0.29 | 0.01 | 5 | Astrocyte |
| *MKI67* | 0.00E+00 | 0.00E+00 | 4.67 | 0.57 | 0.03 | 5 | Astrocyte |
| *CDK1* | 0.00E+00 | 0.00E+00 | 4.61 | 0.31 | 0.01 | 5 | Astrocyte |
| *LOC100231522* | 0.00E+00 | 0.00E+00 | 4.60 | 0.29 | 0.02 | 5 | Astrocyte |
| *FOXM1* | 0.00E+00 | 0.00E+00 | 4.59 | 0.36 | 0.02 | 5 | Astrocyte |
| *DEPDC1B* | 0.00E+00 | 0.00E+00 | 4.58 | 0.27 | 0.01 | 5 | Astrocyte |
| *ECT2* | 0.00E+00 | 0.00E+00 | 4.52 | 0.26 | 0.01 | 5 | Astrocyte |
| *KIF18A* | 0.00E+00 | 0.00E+00 | 4.49 | 0.37 | 0.02 | 5 | Astrocyte |
| *SMC2* | 0.00E+00 | 0.00E+00 | 4.49 | 0.70 | 0.06 | 5 | Astrocyte |
| *EPHA5* | 0.00E+00 | 0.00E+00 | 3.47 | 0.98 | 0.32 | 6 | GABAergic Neuron |
| *DACH2* | 0.00E+00 | 0.00E+00 | 3.30 | 0.83 | 0.19 | 6 | GABAergic Neuron |
| *PLS1* | 0.00E+00 | 0.00E+00 | 3.21 | 0.39 | 0.06 | 6 | GABAergic Neuron |
| *UNC5C* | 0.00E+00 | 0.00E+00 | 3.13 | 0.86 | 0.29 | 6 | GABAergic Neuron |
| *SLAIN1* | 0.00E+00 | 0.00E+00 | 2.84 | 0.86 | 0.27 | 6 | GABAergic Neuron |
| *ARX* | 0.00E+00 | 0.00E+00 | 2.66 | 0.74 | 0.20 | 6 | GABAergic Neuron |
| *OPRM1* | 0.00E+00 | 0.00E+00 | 2.55 | 0.54 | 0.16 | 6 | GABAergic Neuron |
| *MPPED1* | 0.00E+00 | 0.00E+00 | 2.39 | 0.52 | 0.17 | 6 | GABAergic Neuron |
| *PDZRN3* | 0.00E+00 | 0.00E+00 | 2.36 | 0.55 | 0.18 | 6 | GABAergic Neuron |
| *ZNF536* | 0.00E+00 | 0.00E+00 | 2.27 | 0.93 | 0.46 | 6 | GABAergic Neuron |
| *NRXN3* | 0.00E+00 | 0.00E+00 | 2.25 | 1.00 | 0.69 | 6 | GABAergic Neuron |

| Gene | P-Value | FDR | Average $log_2FC$ | Fraction 1 | Fraction 2 | Cluster | Annotation |
|---|---|---|---|---|---|---|---|
| ***SOX6*** | 0.00E+00 | 0.00E+00 | 2.20 | 0.93 | 0.41 | 6 | GABAergic Neuron |
| ***GAD2*** | 0.00E+00 | 0.00E+00 | 2.13 | 0.77 | 0.33 | 6 | GABAergic Neuron |
| ***GAD1*** | 0.00E+00 | 0.00E+00 | 2.09 | 0.61 | 0.25 | 6 | GABAergic Neuron |
| ***ERBB4*** | 0.00E+00 | 0.00E+00 | 2.09 | 0.76 | 0.44 | 6 | GABAergic Neuron |
| ***PLXNA2*** | 0.00E+00 | 0.00E+00 | 1.85 | 0.56 | 0.25 | 6 | GABAergic Neuron |
| ***ZEB2*** | 0.00E+00 | 0.00E+00 | 1.83 | 0.78 | 0.38 | 6 | GABAergic Neuron |
| ***LOC121470376*** | 0.00E+00 | 0.00E+00 | 1.80 | 0.81 | 0.25 | 6 | GABAergic Neuron |
| ***SOX4*** | 0.00E+00 | 0.00E+00 | 1.80 | 0.67 | 0.32 | 6 | GABAergic Neuron |
| ***LOC121470375*** | 6.09E-309 | 1.16E-304 | 1.79 | 0.49 | 0.15 | 6 | GABAergic Neuron |
| ***SYT10*** | 0.00E+00 | 0.00E+00 | 3.80 | 0.55 | 0.09 | 7 | Glutamatergic Neuron |
| ***HMX3*** | 0.00E+00 | 0.00E+00 | 3.60 | 0.30 | 0.03 | 7 | Glutamatergic Neuron |
| ***DDC*** | 0.00E+00 | 0.00E+00 | 3.43 | 0.35 | 0.04 | 7 | Glutamatergic Neuron |
| ***EYS*** | 0.00E+00 | 0.00E+00 | 2.30 | 0.73 | 0.19 | 7 | Glutamatergic Neuron |
| ***SCG2*** | 0.00E+00 | 0.00E+00 | 2.04 | 0.62 | 0.21 | 7 | Glutamatergic Neuron |
| ***PDE7B*** | 0.00E+00 | 0.00E+00 | 1.80 | 0.55 | 0.17 | 7 | Glutamatergic Neuron |
| ***STK32B*** | 3.16E-235 | 6.02E-231 | 1.78 | 0.47 | 0.17 | 7 | Glutamatergic Neuron |
| ***LOC100221646*** | 0.00E+00 | 0.00E+00 | 1.76 | 0.59 | 0.20 | 7 | Glutamatergic Neuron |
| ***EPB41L4B*** | 5.20E-284 | 9.89E-280 | 1.71 | 0.56 | 0.21 | 7 | Glutamatergic Neuron |
| ***UNC80*** | 0.00E+00 | 0.00E+00 | 1.69 | 0.69 | 0.26 | 7 | Glutamatergic Neuron |
| ***GLRA2*** | 3.50E-196 | 6.65E-192 | 1.68 | 0.44 | 0.17 | 7 | Glutamatergic Neuron |
| ***NRP2*** | 5.30E-248 | 1.01E-243 | 1.68 | 0.49 | 0.17 | 7 | Glutamatergic Neuron |
| ***KCTD8*** | 5.91E-166 | 1.12E-161 | 1.54 | 0.42 | 0.16 | 7 | Glutamatergic Neuron |
| ***RPS6KA2*** | 3.26E-194 | 6.20E-190 | 1.47 | 0.53 | 0.23 | 7 | Glutamatergic Neuron |
| ***BRINP1*** | 6.22E-259 | 1.18E-254 | 1.45 | 0.62 | 0.27 | 7 | Glutamatergic Neuron |
| ***CELF4*** | 0.00E+00 | 0.00E+00 | 1.35 | 0.99 | 0.74 | 7 | Glutamatergic Neuron |
| ***TOX2*** | 4.46E-210 | 8.48E-206 | 1.35 | 0.59 | 0.28 | 7 | Glutamatergic Neuron |
| ***SHISA9*** | 1.12E-222 | 2.13E-218 | 1.30 | 0.62 | 0.29 | 7 | Glutamatergic Neuron |

| Gene | P-Value | FDR | Average $log_2FC$ | Fraction 1 | Fraction 2 | Cluster | Annotation |
|---|---|---|---|---|---|---|---|
| *SDK1* | 9.76E-155 | 1.86E-150 | 1.30 | 0.51 | 0.25 | 7 | Glutamatergic Neuron |
| *NTRK3* | 1.38E-223 | 2.63E-219 | 1.28 | 0.65 | 0.32 | 7 | Glutamatergic Neuron |
| *LOC115496794* | 0.00E+00 | 0.00E+00 | 4.96 | 0.51 | 0.02 | 8 | Glutamatergic Neuron |
| *SATB2* | 0.00E+00 | 0.00E+00 | 3.79 | 0.28 | 0.02 | 8 | Glutamatergic Neuron |
| *ADCYAP1* | 0.00E+00 | 0.00E+00 | 3.45 | 0.62 | 0.07 | 8 | Glutamatergic Neuron |
| *RASGRF2* | 0.00E+00 | 0.00E+00 | 3.16 | 0.41 | 0.06 | 8 | Glutamatergic Neuron |
| *LOC100226371* | 0.00E+00 | 0.00E+00 | 3.14 | 0.62 | 0.09 | 8 | Glutamatergic Neuron |
| *GAS2* | 0.00E+00 | 0.00E+00 | 3.09 | 0.56 | 0.08 | 8 | Glutamatergic Neuron |
| *LOC121470895* | 0.00E+00 | 0.00E+00 | 3.02 | 0.37 | 0.05 | 8 | Glutamatergic Neuron |
| *LOC121470712* | 0.00E+00 | 0.00E+00 | 2.88 | 0.45 | 0.07 | 8 | Glutamatergic Neuron |
| *CBLN1* | 0.00E+00 | 0.00E+00 | 2.85 | 0.37 | 0.05 | 8 | Glutamatergic Neuron |
| *LOC115494899* | 0.00E+00 | 0.00E+00 | 2.84 | 0.45 | 0.08 | 8 | Glutamatergic Neuron |
| *LOC121470713* | 0.00E+00 | 0.00E+00 | 2.72 | 0.74 | 0.14 | 8 | Glutamatergic Neuron |
| *LOC115497347* | 5.77E-267 | 1.10E-262 | 2.58 | 0.38 | 0.09 | 8 | Glutamatergic Neuron |
| *SLC17A6* | 0.00E+00 | 0.00E+00 | 2.55 | 0.49 | 0.09 | 8 | Glutamatergic Neuron |
| *RBFOX1* | 0.00E+00 | 0.00E+00 | 2.53 | 1.00 | 0.47 | 8 | Glutamatergic Neuron |
| *SLIT3* | 0.00E+00 | 0.00E+00 | 2.49 | 0.78 | 0.22 | 8 | Glutamatergic Neuron |
| *CACNA2D1* | 0.00E+00 | 0.00E+00 | 2.38 | 0.95 | 0.42 | 8 | Glutamatergic Neuron |
| *LOC100221630* | 0.00E+00 | 0.00E+00 | 2.20 | 0.66 | 0.16 | 8 | Glutamatergic Neuron |
| *LOC100221646* | 0.00E+00 | 0.00E+00 | 2.11 | 0.73 | 0.20 | 8 | Glutamatergic Neuron |
| *CDH9* | 1.82E-198 | 3.45E-194 | 2.11 | 0.37 | 0.12 | 8 | Glutamatergic Neuron |
| *TENM2* | 0.00E+00 | 0.00E+00 | 2.09 | 0.90 | 0.46 | 8 | Glutamatergic Neuron |
| *PLS1* | 0.00E+00 | 0.00E+00 | 3.37 | 0.40 | 0.06 | 9 | GABAergic Neuron |
| *LOC121470376* | 0.00E+00 | 0.00E+00 | 3.13 | 0.96 | 0.24 | 9 | GABAergic Neuron |
| *LHX6* | 0.00E+00 | 0.00E+00 | 3.08 | 0.31 | 0.06 | 9 | GABAergic Neuron |
| *LOC121470375* | 0.00E+00 | 0.00E+00 | 3.03 | 0.75 | 0.14 | 9 | GABAergic Neuron |
| *ST18* | 0.00E+00 | 0.00E+00 | 2.93 | 0.61 | 0.13 | 9 | GABAergic Neuron |

| Gene | P-Value | FDR | Average log$_2$FC | Fraction 1 | Fraction 2 | Cluster | Annotation |
|---|---|---|---|---|---|---|---|
| ***LOC121469489*** | 0.00E+00 | 0.00E+00 | 2.91 | 0.66 | 0.15 | 9 | GABAergic Neuron |
| ***MYT1*** | 0.00E+00 | 0.00E+00 | 2.64 | 0.80 | 0.26 | 9 | GABAergic Neuron |
| ***DLX1*** | 0.00E+00 | 0.00E+00 | 2.54 | 0.36 | 0.10 | 9 | GABAergic Neuron |
| ***ARX*** | 0.00E+00 | 0.00E+00 | 2.41 | 0.67 | 0.20 | 9 | GABAergic Neuron |
| ***PLXNA2*** | 0.00E+00 | 0.00E+00 | 2.33 | 0.65 | 0.25 | 9 | GABAergic Neuron |
| ***PTK2*** | 0.00E+00 | 0.00E+00 | 2.19 | 0.82 | 0.48 | 9 | GABAergic Neuron |
| ***SOX6*** | 0.00E+00 | 0.00E+00 | 2.17 | 0.91 | 0.41 | 9 | GABAergic Neuron |
| ***DACH2*** | 0.00E+00 | 0.00E+00 | 2.14 | 0.58 | 0.20 | 9 | GABAergic Neuron |
| ***MYO1B*** | 0.00E+00 | 0.00E+00 | 2.02 | 0.52 | 0.22 | 9 | GABAergic Neuron |
| ***EPHA5*** | 0.00E+00 | 0.00E+00 | 1.99 | 0.72 | 0.33 | 9 | GABAergic Neuron |
| ***BRINP3*** | 0.00E+00 | 0.00E+00 | 1.94 | 0.71 | 0.38 | 9 | GABAergic Neuron |
| ***SOX4*** | 0.00E+00 | 0.00E+00 | 1.89 | 0.70 | 0.33 | 9 | GABAergic Neuron |
| ***SLAIN1*** | 0.00E+00 | 0.00E+00 | 1.87 | 0.64 | 0.28 | 9 | GABAergic Neuron |
| ***LOC115496358*** | 0.00E+00 | 0.00E+00 | 1.80 | 0.61 | 0.24 | 9 | GABAergic Neuron |
| ***LMO1*** | 2.83E-318 | 5.38E-314 | 1.77 | 0.64 | 0.32 | 9 | GABAergic Neuron |

**Supplemental Table 5:** Sex-stratified differential gene expression in female embryonic brain clusters: Differential expression analysis comparing heat call-exposed (n=4) versus control (n=4) zebra finch embryos, where n represents pooled biological replicates (E13 medial hypothalamic punches from 5 embryos per pool), analyzed separately by sex. The analysis was performed using the MAST test (two-tailed) on snRNA-seq data with min.pct=0.25 and log2FC threshold=0.25. FDR is the FDR-corrected P-value using the Benjamini-Hochberg method. Columns show the P-Value = unadjusted p-value; average log2 fold change; Fraction 1 = the proportion of cells expressing the gene in the heat call group; Fraction 2 = proportion of cells expressing gene in the control group; Cluster = WNN cluster ID; Sex = biological sex; Gene = gene symbol.

| P-Value | FDR | Average log2FC | Fraction 1 | Fraction 2 | Cluster | Sex | Gene |
|---|---|---|---|---|---|---|---|
| **8.96E-28** | 1.70E-23 | -1.76 | 0.18 | 0.47 | 4 | Female | *TTR* |
| **9.48E-27** | 1.80E-22 | -2.09 | 0.16 | 0.47 | 7 | Female | *TTR* |
| **1.53E-25** | 2.91E-21 | -1.42 | 0.18 | 0.41 | 0 | Female | *TTR* |

| P-Value | FDR | Average log2FC | Fraction 1 | Fraction 2 | Cluster | Sex | Gene |
|---|---|---|---|---|---|---|---|
| **2.26E-20** | 4.30E-16 | -1.46 | 0.18 | 0.40 | 2 | Female | *TTR* |
| **4.57E-20** | 8.69E-16 | -1.72 | 0.15 | 0.43 | 8 | Female | *TTR* |
| **1.76E-18** | 3.35E-14 | -1.95 | 0.16 | 0.48 | 16 | Female | *TTR* |
| **1.87E-14** | 3.56E-10 | -1.72 | 0.16 | 0.44 | 14 | Female | *TTR* |
| **1.41E-12** | 2.69E-08 | -1.17 | 0.25 | 0.43 | 3 | Female | *TTR* |
| **1.75E-11** | 3.32E-07 | 1.89 | 0.32 | 0.19 | 8 | Female | *LOC121470611* |
| **1.94E-11** | 3.69E-07 | -0.72 | 0.41 | 0.57 | 3 | Female | *SPTBN4* |
| **8.34E-11** | 1.59E-06 | 0.68 | 0.40 | 0.24 | 0 | Female | *LOC116806978* |
| **8.43E-10** | 1.60E-05 | -0.26 | 0.43 | 0.62 | 6 | Female | *CSMD1* |
| **1.80E-09** | 3.41E-05 | 0.45 | 0.84 | 0.73 | 4 | Female | *HSP90AA1* |
| **2.32E-09** | 4.41E-05 | -0.27 | 0.33 | 0.53 | 11 | Female | *CELF4* |
| **6.25E-09** | 1.19E-04 | -1.47 | 0.18 | 0.33 | 10 | Female | *TTR* |
| **6.68E-09** | 1.27E-04 | -1.34 | 0.19 | 0.39 | 12 | Female | *TTR* |
| **7.68E-09** | 1.46E-04 | -1.11 | 0.19 | 0.34 | 9 | Female | *TTR* |
| **9.38E-09** | 1.78E-04 | 1.44 | 0.41 | 0.17 | 20 | Female | *LOC116806978* |
| **1.36E-08** | 2.59E-04 | -0.81 | 0.36 | 0.48 | 11 | Female | *FUS* |
| **1.93E-08** | 3.67E-04 | -0.39 | 0.52 | 0.70 | 9 | Female | *LOC100229373* |
| **2.90E-08** | 5.51E-04 | 0.75 | 0.33 | 0.19 | 2 | Female | *LOC116806978* |
| **3.78E-08** | 7.19E-04 | -1.37 | 0.20 | 0.35 | 13 | Female | *TTR* |
| **5.74E-08** | 1.09E-03 | 0.68 | 0.41 | 0.26 | 2 | Female | *IKZF2* |
| **5.75E-08** | 1.09E-03 | 1.12 | 0.28 | 0.16 | 8 | Female | *DOT1L* |
| **8.02E-08** | 1.53E-03 | -0.67 | 0.43 | 0.56 | 5 | Female | *FUS* |
| **8.44E-08** | 1.60E-03 | 3.68 | 0.60 | 0.08 | 40 | Female | *FIGN* |
| **1.16E-07** | 2.21E-03 | -1.15 | 0.19 | 0.40 | 19 | Female | *TTR* |
| **1.25E-07** | 2.37E-03 | 0.53 | 0.38 | 0.23 | 7 | Female | *LOC116806978* |
| **1.54E-07** | 2.93E-03 | 2.51 | 0.80 | 0.38 | 40 | Female | *SOX5* |
| **2.05E-07** | 3.90E-03 | 0.27 | 0.98 | 0.97 | 0 | Female | *LOC121469074* |

| P-Value | FDR | Average log2FC | Fraction 1 | Fraction 2 | Cluster | Sex | Gene |
|---|---|---|---|---|---|---|---|
| **2.51E-07** | 4.77E-03 | -0.37 | 0.30 | 0.30 | 1 | Female | *STAG1* |
| **3.49E-07** | 6.64E-03 | 3.02 | 0.68 | 0.18 | 40 | Female | *COL27A1* |
| **4.68E-07** | 8.90E-03 | -1.65 | 0.19 | 0.45 | 27 | Female | *TTR* |
| **4.76E-07** | 9.05E-03 | -2.06 | 0.15 | 0.39 | 28 | Female | *TTR* |
| **5.03E-07** | 9.57E-03 | -0.99 | 0.22 | 0.41 | 15 | Female | *TTR* |
| **5.70E-07** | 1.08E-02 | 1.26 | 0.45 | 0.22 | 21 | Female | *HS3ST5* |
| **5.77E-07** | 1.10E-02 | -1.47 | 0.25 | 0.37 | 17 | Female | *TTR* |
| **6.02E-07** | 1.15E-02 | -1.74 | 0.16 | 0.29 | 18 | Female | *TTR* |
| **6.20E-07** | 1.18E-02 | 1.54 | 0.44 | 0.37 | 8 | Female | *LOC115497347* |
| **6.27E-07** | 1.19E-02 | -0.31 | 0.29 | 0.44 | 9 | Female | *CNTNAP2* |
| **8.03E-07** | 1.53E-02 | 0.78 | 0.42 | 0.25 | 8 | Female | *LOC116806978* |
| **8.60E-07** | 1.63E-02 | 2.39 | 0.40 | 0.21 | 40 | Female | *ADK* |
| **8.74E-07** | 1.66E-02 | 0.50 | 0.62 | 0.45 | 7 | Female | *LOC116806985* |
| **8.76E-07** | 1.67E-02 | -5.36 | 0.03 | 0.51 | 48 | Female | *ROBO2* |
| **9.42E-07** | 1.79E-02 | -0.42 | 0.49 | 0.59 | 0 | Female | *SPTBN4* |
| **1.05E-06** | 1.99E-02 | 2.82 | 0.64 | 0.13 | 40 | Female | *COL9A3* |
| **1.12E-06** | 2.13E-02 | 2.81 | 0.68 | 0.16 | 40 | Female | *COL2A1* |
| **1.17E-06** | 2.22E-02 | -2.12 | 0.14 | 0.51 | 32 | Female | *TTR* |
| **1.19E-06** | 2.27E-02 | 0.82 | 0.41 | 0.23 | 12 | Female | *LOC116806978* |
| **1.34E-06** | 2.54E-02 | -0.83 | 0.18 | 0.39 | 20 | Female | *TTR* |
| **1.37E-06** | 2.60E-02 | 1.37 | 0.44 | 0.18 | 28 | Female | *LOC116806978* |
| **1.41E-06** | 2.69E-02 | 0.71 | 0.29 | 0.15 | 14 | Female | *RPS6KA3* |
| **1.99E-06** | 3.78E-02 | 2.75 | 0.60 | 0.14 | 40 | Female | *COL9A1* |
| **2.10E-06** | 3.99E-02 | 2.51 | 0.60 | 0.23 | 40 | Female | *MMP16* |
| **2.16E-06** | 4.10E-02 | -0.37 | 0.52 | 0.68 | 9 | Female | *CSMD1* |
| **2.16E-06** | 4.12E-02 | 0.57 | 0.82 | 0.61 | 20 | Female | *TSNARE1* |
| **2.18E-06** | 4.14E-02 | -0.45 | 0.35 | 0.36 | 11 | Female | *BRD1* |

| P-Value | FDR | Average log2FC | Fraction 1 | Fraction 2 | Cluster | Sex | Gene |
|---|---|---|---|---|---|---|---|
| **2.56E-06** | 4.86E-02 | -0.62 | 0.47 | 0.68 | 17 | Female | *CSMD1* |
| **2.61E-06** | 4.95E-02 | 0.77 | 0.70 | 0.66 | 8 | Female | *PDE10A* |
| **1.26E-22** | 2.40E-18 | -0.39 | 0.94 | 0.97 | 4 | Male | *LOC121469074* |
| **3.52E-18** | 6.69E-14 | 0.73 | 0.63 | 0.42 | 1 | Male | *FGF14* |
| **4.82E-15** | 9.16E-11 | -1.13 | 0.28 | 0.42 | 2 | Male | *CRIM1* |
| **6.33E-13** | 1.20E-08 | -0.85 | 0.84 | 0.72 | 26 | Male | *DSCAM* |
| **6.99E-13** | 1.33E-08 | -0.41 | 0.92 | 0.94 | 2 | Male | *FABP7* |
| **1.93E-12** | 3.68E-08 | 1.93 | 0.72 | 0.22 | 26 | Male | *PLXNA4* |
| **2.21E-11** | 4.20E-07 | -0.78 | 0.90 | 0.99 | 34 | Male | *LOC121469074* |
| **2.93E-11** | 5.56E-07 | -3.11 | 0.28 | 0.68 | 26 | Male | *MTUS2* |
| **6.69E-11** | 1.27E-06 | -0.30 | 0.94 | 0.98 | 8 | Male | *LOC121469074* |
| **6.82E-11** | 1.30E-06 | 2.47 | 0.52 | 0.11 | 26 | Male | *LOC116808653* |
| **2.18E-10** | 4.14E-06 | 2.33 | 0.42 | 0.07 | 26 | Male | *LOC105760897* |
| **2.55E-10** | 4.84E-06 | 2.26 | 0.56 | 0.14 | 26 | Male | *CACNB2* |
| **7.97E-10** | 1.52E-05 | 2.59 | 0.46 | 0.12 | 26 | Male | *PLXDC2* |
| **1.01E-09** | 1.92E-05 | 0.44 | 0.83 | 0.66 | 9 | Male | *FGF14* |
| **1.44E-09** | 2.74E-05 | 0.49 | 0.63 | 0.43 | 11 | Male | *FGF14* |
| **1.51E-09** | 2.87E-05 | 0.65 | 0.66 | 0.47 | 11 | Male | *CADM2* |
| **1.68E-09** | 3.20E-05 | -0.63 | 0.64 | 0.80 | 13 | Male | *KCNH8* |
| **2.15E-09** | 4.09E-05 | 1.50 | 0.82 | 0.41 | 26 | Male | *TENM3* |
| **2.25E-09** | 4.29E-05 | 0.63 | 0.32 | 0.24 | 0 | Male | *RPLP1* |
| **3.14E-09** | 5.97E-05 | -0.84 | 0.27 | 0.37 | 4 | Male | *THSD7A* |
| **4.08E-09** | 7.77E-05 | 0.46 | 0.58 | 0.43 | 2 | Male | *FGF14* |
| **4.36E-09** | 8.29E-05 | 2.27 | 0.58 | 0.17 | 26 | Male | *CNTN5* |
| **5.52E-09** | 1.05E-04 | -0.50 | 0.66 | 0.78 | 6 | Male | *CCSER1* |
| **6.18E-09** | 1.17E-04 | 0.56 | 0.65 | 0.53 | 5 | Male | *FGF14* |
| **7.02E-09** | 1.34E-04 | 2.25 | 0.46 | 0.10 | 26 | Male | *KCNH1* |

| P-Value | FDR | Average log2FC | Fraction 1 | Fraction 2 | Cluster | Sex | Gene |
|---|---|---|---|---|---|---|---|
| **1.42E-08** | 2.69E-04 | 2.39 | 0.44 | 0.10 | 26 | Male | *NR2F2* |
| **2.19E-08** | 4.17E-04 | 1.32 | 0.86 | 0.54 | 26 | Male | *TENM4* |
| **2.59E-08** | 4.92E-04 | -1.81 | 0.50 | 0.74 | 26 | Male | *DAB1* |
| **3.04E-08** | 5.79E-04 | -0.76 | 0.60 | 0.73 | 17 | Male | *CCSER1* |
| **3.80E-08** | 7.22E-04 | 1.73 | 0.56 | 0.17 | 26 | Male | *KIF26B* |
| **4.22E-08** | 8.03E-04 | 3.19 | 0.30 | 0.04 | 26 | Male | *PLXNA2* |
| **4.74E-08** | 9.01E-04 | 3.11 | 0.30 | 0.04 | 26 | Male | *ANO6* |
| **6.17E-08** | 1.17E-03 | 0.60 | 0.64 | 0.48 | 11 | Male | *LOC116808807* |
| **1.13E-07** | 2.15E-03 | 2.19 | 0.32 | 0.05 | 26 | Male | *LOC121469249* |
| **1.31E-07** | 2.48E-03 | 1.66 | 0.44 | 0.11 | 26 | Male | *RASGEF1A* |
| **1.51E-07** | 2.87E-03 | -0.81 | 0.38 | 0.51 | 11 | Male | *NRP1* |
| **1.78E-07** | 3.39E-03 | 1.19 | 0.86 | 0.52 | 26 | Male | *ARPP21* |
| **2.04E-07** | 3.88E-03 | 0.90 | 0.34 | 0.21 | 8 | Male | *LOC116807842* |
| **2.44E-07** | 4.64E-03 | 0.51 | 0.88 | 0.50 | 26 | Male | *GRIK2* |
| **2.49E-07** | 4.73E-03 | 2.13 | 0.26 | 0.04 | 26 | Male | *SV2C* |
| **2.85E-07** | 5.42E-03 | -5.06 | 0.00 | 0.25 | 26 | Male | *TFAP2B* |
| **3.28E-07** | 6.24E-03 | -2.23 | 0.22 | 0.58 | 26 | Male | *THSD7A* |
| **3.35E-07** | 6.36E-03 | -0.72 | 0.82 | 0.98 | 35 | Male | *FIGN* |
| **3.46E-07** | 6.58E-03 | -0.51 | 0.16 | 0.26 | 1 | Male | *IARS1* |
| **3.77E-07** | 7.17E-03 | 1.43 | 0.44 | 0.13 | 26 | Male | *MPDZ* |
| **3.80E-07** | 7.22E-03 | -2.76 | 0.22 | 0.54 | 26 | Male | *ZEB2* |
| **5.65E-07** | 1.07E-02 | 3.44 | 0.31 | 0.07 | 34 | Male | *CCK* |
| **7.26E-07** | 1.38E-02 | -0.61 | 0.38 | 0.53 | 11 | Male | *ROBO2* |
| **7.54E-07** | 1.43E-02 | 2.09 | 0.42 | 0.11 | 26 | Male | *LOC116807842* |
| **7.57E-07** | 1.44E-02 | -0.63 | 0.41 | 0.56 | 11 | Male | *PDE3A* |
| **9.06E-07** | 1.72E-02 | -0.82 | 0.37 | 0.54 | 13 | Male | *NRP1* |
| **9.11E-07** | 1.73E-02 | -0.60 | 0.76 | 0.87 | 21 | Male | *TMEFF1* |

| P-Value | FDR | Average log2FC | Fraction 1 | Fraction 2 | Cluster | Sex | Gene |
|---|---|---|---|---|---|---|---|
| **1.01E-06** | 1.91E-02 | 0.47 | 0.47 | 0.34 | 2 | Male | *GRIA2* |
| **1.21E-06** | 2.31E-02 | 0.52 | 0.48 | 0.36 | 3 | Male | *SPTBN4* |
| **1.33E-06** | 2.53E-02 | 1.09 | 0.26 | 0.11 | 16 | Male | *TTR* |
| **1.52E-06** | 2.90E-02 | 0.59 | 0.48 | 0.37 | 2 | Male | *FGFRL1* |
| **1.53E-06** | 2.91E-02 | 2.34 | 0.34 | 0.07 | 26 | Male | *DPYD* |
| **1.73E-06** | 3.29E-02 | 2.27 | 0.26 | 0.04 | 26 | Male | *RIT2* |
| **1.75E-06** | 3.32E-02 | 1.70 | 0.42 | 0.12 | 26 | Male | *KIF1A* |
| **2.15E-06** | 4.09E-02 | 0.78 | 0.40 | 0.25 | 11 | Male | *EPHA4* |
| **2.18E-06** | 4.14E-02 | 1.81 | 0.48 | 0.18 | 26 | Male | *LOC115491411* |
| **2.28E-06** | 4.34E-02 | 0.49 | 0.74 | 0.42 | 26 | Male | *LOC100217927* |
| **2.29E-06** | 4.35E-02 | 1.60 | 0.56 | 0.23 | 26 | Male | *HSPA8* |

**Supplemental Table 6: Differential chromatin accessibility by cell type.** Number of tested peaks and significantly differentially accessible (DA) peaks between heat call and Control for each annotated cell type, with DA peaks partitioned into higher vs lower accessibility in heat Call (FDR < 0.05). Percentages are calculated relative to the total number of DA peaks across all cell types.

| Cell Type | Total Peaks Tested | Significant DA Peaks | % of All DA Peaks | Higher Accessibility (n) | Higher Accessibility (%) | Lower Accessibility (n) | Lower Accessibility (%) |
|---|---|---|---|---|---|---|---|
| Astrocyte | 95755 | 204 | 86.4 | 197 | 96.6 | 7 | 3.4 |
| GABAergic Neuron | 34737 | 25 | 10.6 | 19 | 76 | 6 | 24 |
| Glutamatergic Neuron | 87264 | 4 | 1.7 | 3 | 75 | 1 | 25 |
| Glutamatergic/GABA | 12772 | 2 | 0.8 | 2 | 100 | 0 | 0 |
| Endothelial Cell | 24883 | 1 | 0.4 | 1 | 100 | 0 | 0 |
| Capillary Endothelial | 8899 | 0 | 0 | 0 | NA | 0 | NA |
| Choroid Plexus Epithelial Cell | 13158 | 0 | 0 | 0 | NA | 0 | NA |
| Ependymal Cell | 9529 | 0 | 0 | 0 | NA | 0 | NA |
| Fibroblast | 5468 | 0 | 0 | 0 | NA | 0 | NA |
| Macrophage | 6364 | 0 | 0 | 0 | NA | 0 | NA |
| Microglia | 7283 | 0 | 0 | 0 | NA | 0 | NA |
| Oligodendrocyte | 7271 | 0 | 0 | 0 | NA | 0 | NA |
| Vascular Associated Smooth Muscle Cell | 10203 | 0 | 0 | 0 | NA | 0 | NA |

**Supplemental Table 7: Top differentially active TF motifs (chromVAR).** Top 20 transcription factor motifs with increased chromVAR deviation in heat Call versus control embryos, ranked by adjusted P value from differential chromVAR activity testing (Wilcoxon rank-sum test). For each motif (JASPAR ID), the table reports the mapped TF name and family, the mean deviation difference (Mean Deviation), the raw P-value, the FDR-adjusted P-value (FDR), and $-\log_{10}$(FDR).

| JASPAR ID | TF Name | TF Family | Mean Deviation | Raw P-value | FDR | -log10(FDR) |
|---|---|---|---|---|---|---|
| MA1527.1 | *NFIC* | *Nuclear factor 1* | 0.14 | 1.97E-12 | 1.47E-09 | 8.83 |
| MA1142.1 | *FOSL1::JUND* | *Fos-related::Jun-related* | 0.12 | 5.26E-12 | 3.92E-09 | 8.41 |
| MA0462.2 | *BATF::JUN* | *B-ATF-related factors::Jun-related* | 0.12 | 5.88E-12 | 4.39E-09 | 8.36 |
| MA0835.2 | *BATF3* | *B-ATF-related factors* | 0.12 | 2.03E-11 | 1.51E-08 | 7.82 |
| MA1634.1 | *BATF* | *B-ATF-related factors* | 0.12 | 2.58E-11 | 1.92E-08 | 7.72 |
| MA0491.2 | *JUND* | *Jun-related* | 0.12 | 6.95E-11 | 5.19E-08 | 7.28 |
| MA1528.1 | *NFIX* | *Nuclear factor 1* | 0.12 | 2.20E-10 | 1.64E-07 | 6.78 |
| MA1643.1 | *NFIB* | *Nuclear factor 1* | 0.13 | 2.31E-10 | 1.72E-07 | 6.76 |
| MA0119.1 | *NFIC::TLX1* | *Nuclear factor 1::NK* | 0.11 | 5.79E-10 | 4.32E-07 | 6.36 |
| MA0655.1 | *JDP2* | *Fos-related* | 0.10 | 1.59E-09 | 1.19E-06 | 5.92 |
| MA0489.1 | *JUN* | *Jun-related* | 0.12 | 9.82E-09 | 7.32E-06 | 5.14 |
| MA0161.2 | *NFIC* | *Nuclear factor 1* | 0.13 | 1.14E-08 | 8.51E-06 | 5.07 |
| MA1141.1 | *FOS::JUND* | *Fos-related::Jun-related* | 0.12 | 1.24E-08 | 9.23E-06 | 5.03 |
| MA0490.2 | *JUNB* | *Jun-related* | 0.11 | 2.23E-08 | 1.66E-05 | 4.78 |
| MA1132.1 | *JUN::JUNB* | *Jun-related* | 0.10 | 3.68E-08 | 2.74E-05 | 4.56 |
| MA1622.1 | *Smad2::Smad3* | *SMAD factors::SMAD factors* | 0.12 | 4.37E-08 | 3.26E-05 | 4.49 |
| MA0476.1 | *FOS* | *Fos-related* | 0.11 | 5.22E-08 | 3.89E-05 | 4.41 |
| MA0478.1 | *FOSL2* | *Fos-related* | 0.11 | 7.54E-08 | 5.62E-05 | 4.25 |
| MA0477.2 | *FOSL1* | *Fos-related* | 0.12 | 1.11E-07 | 8.30E-05 | 4.08 |
| MA1138.1 | *FOSL2::JUNB* | *Fos-related::Jun-related* | 0.11 | 1.15E-07 | 8.58E-05 | 4.07 |

**Supplemental Table 8: Astrocyte subcluster marker genes.** The top 10 differentially expressed marker genes defining each of 4 WNN astrocyte subclusters in the E13 zebra finch hypothalamus, identified using FindAllMarkers (Wilcoxon rank-sum test; adjusted $P < 0.05$, log2FC $\geq 0.10$, pct.1 $\geq 0.10$) from the snRNA-seq dataset and ranked by log2 fold-change. Subclusters were identified at resolution 0.06 using weighted nearest neighbor (WNN) integration of RNA and ATAC modalities. Columns: gene (marker gene symbol), cluster (WNN astrocyte subcluster ID), P-value (raw), average log2FC (average log2 fold-change for the subcluster versus all other subclusters), Fraction 1 (fraction of nuclei in the subcluster expressing the gene), Fraction 2 (fraction of nuclei in all other subclusters expressing the gene), and FDR (FDR-adjusted P-value).

| Gene | Cluster | P-Value | FDR | Average log2FC | Fraction 1 | Fraction 2 |
|---|---|---|---|---|---|---|
| ***LOC105758645*** | 0 | 0.00E+00 | 0.00E+00 | 2.61 | 0.92 | 0.31 |
| ***KCNH8*** | 0 | 3.12E-188 | 6.10E-184 | 2.46 | 0.25 | 0.09 |
| ***CPA6*** | 0 | 7.08E-50 | 1.39E-45 | 2.09 | 0.10 | 0.04 |
| ***LOC121470970*** | 0 | 2.56E-80 | 5.01E-76 | 2.06 | 0.12 | 0.04 |
| ***PDE9A*** | 0 | 4.01E-58 | 7.84E-54 | 1.94 | 0.12 | 0.05 |
| ***KCNQ5*** | 0 | 0.00E+00 | 0.00E+00 | 1.92 | 0.63 | 0.26 |
| ***LOC115497609*** | 0 | 7.55E-78 | 1.48E-73 | 1.84 | 0.13 | 0.05 |
| ***FOXG1*** | 0 | 0.00E+00 | 0.00E+00 | 1.79 | 0.87 | 0.41 |
| ***FAM53A*** | 0 | 0.00E+00 | 0.00E+00 | 1.69 | 0.61 | 0.36 |
| ***CD24*** | 0 | 6.10E-105 | 1.20E-100 | 1.68 | 0.27 | 0.15 |
| ***NDNF*** | 1 | 2.26E-299 | 4.42E-295 | 4.28 | 0.17 | 0.01 |
| ***THRB*** | 1 | 0.00E+00 | 0.00E+00 | 3.80 | 0.44 | 0.05 |
| ***LMX1B*** | 1 | 9.92E-144 | 1.94E-139 | 3.67 | 0.11 | 0.02 |
| ***IRX1*** | 1 | 0.00E+00 | 0.00E+00 | 3.60 | 0.19 | 0.02 |
| ***LOC115496633*** | 1 | 0.00E+00 | 0.00E+00 | 3.57 | 0.22 | 0.02 |
| ***VIT*** | 1 | 0.00E+00 | 0.00E+00 | 3.56 | 0.30 | 0.03 |
| ***STARD9*** | 1 | 0.00E+00 | 0.00E+00 | 3.48 | 0.44 | 0.08 |
| ***SYNPO2*** | 1 | 1.09E-230 | 2.13E-226 | 3.38 | 0.15 | 0.02 |
| ***FGFBP3*** | 1 | 4.43E-294 | 8.67E-290 | 3.26 | 0.19 | 0.02 |
| ***ADAM12*** | 1 | 0.00E+00 | 0.00E+00 | 3.19 | 0.32 | 0.04 |
| ***TTF2*** | 2 | 0.00E+00 | 0.00E+00 | 4.76 | 0.59 | 0.04 |

| Gene | Cluster | P-Value | FDR | Average log2FC | Fraction 1 | Fraction 2 |
|---|---|---|---|---|---|---|
| ***MELK*** | 2 | 0.00E+00 | 0.00E+00 | 4.64 | 0.17 | 0.01 |
| ***AURKA*** | 2 | 2.09E-271 | 4.09E-267 | 4.61 | 0.14 | 0.01 |
| ***MXD3*** | 2 | 1.86E-268 | 3.64E-264 | 4.61 | 0.14 | 0.01 |
| ***CDC20*** | 2 | 2.05E-234 | 4.01E-230 | 4.60 | 0.12 | 0.01 |
| ***KPNA2*** | 2 | 0.00E+00 | 0.00E+00 | 4.53 | 0.19 | 0.01 |
| ***ANLN*** | 2 | 0.00E+00 | 0.00E+00 | 4.49 | 0.36 | 0.02 |
| ***KIF23*** | 2 | 0.00E+00 | 0.00E+00 | 4.45 | 0.21 | 0.01 |
| ***CENPE*** | 2 | 0.00E+00 | 0.00E+00 | 4.44 | 0.49 | 0.04 |
| ***NEK2*** | 2 | 0.00E+00 | 0.00E+00 | 4.43 | 0.40 | 0.03 |
| ***SHH*** | 3 | 0.00E+00 | 0.00E+00 | 4.90 | 0.56 | 0.04 |
| ***CRACR2A*** | 3 | 0.00E+00 | 0.00E+00 | 4.32 | 0.24 | 0.02 |
| ***TSHR*** | 3 | 0.00E+00 | 0.00E+00 | 3.93 | 0.21 | 0.02 |
| ***LOC115494438*** | 3 | 5.27E-245 | 1.03E-240 | 3.81 | 0.15 | 0.01 |
| ***MOXD1*** | 3 | 0.00E+00 | 0.00E+00 | 3.79 | 0.22 | 0.02 |
| ***SLIT2*** | 3 | 0.00E+00 | 0.00E+00 | 3.69 | 0.97 | 0.37 |
| ***ITGA8*** | 3 | 4.02E-244 | 7.88E-240 | 3.68 | 0.17 | 0.02 |
| ***NKX2-4*** | 3 | 2.03E-221 | 3.98E-217 | 3.40 | 0.15 | 0.02 |
| ***LOC100227924*** | 3 | 1.44E-298 | 2.83E-294 | 3.24 | 0.20 | 0.02 |
| ***LOC115494899*** | 3 | 4.48E-136 | 8.77E-132 | 3.22 | 0.12 | 0.02 |

**Supplemental Table 9: Astro-M7 astrocyte hdWGCNA module genes.** Genes assigned to astrocyte co-expression module Astro-M7 (n = 50) identified by hdWGCNA on metacell-aggregated astrocyte expression. Astro-M7 exhibited the strongest positive module–trait correlation with prenatal heat call exposure. Eigengene-based connectivity (kME) was computed as the Pearson correlation between each gene's single-cell expression and each module eigengene.

| Gene | Module | Color | kME grey | kME Astro- M1 | kME Astro- M2 | kME Astro- M3 | kME Astro-M4 | kME Astro-M5 | kME Astro-M6 | kME Astro-M7 | kME Astro-M8 | kME Astro-M9 | kME Astro-M10 | kME Astro-M11 |
|---|---|---|---|---|---|---|---|---|---|---|---|---|---|---|
| ***POU2F1*** | Astro-M7 | magenta | -0.00 | 0.02 | 0.17 | -0.11 | -0.15 | 0.09 | 0.04 | 0.27 | 0.01 | 0.08 | 0.04 | 0.05 |
| ***EPHA3*** | Astro-M7 | magenta | -0.11 | -0.13 | 0.17 | -0.25 | -0.31 | 0.20 | 0.01 | 0.43 | 0.02 | 0.16 | -0.01 | -0.05 |
| ***ZBTB20*** | Astro-M7 | magenta | -0.20 | 0.01 | 0.34 | -0.10 | -0.41 | 0.16 | -0.01 | 0.46 | -0.16 | 0.07 | -0.01 | -0.03 |
| ***TMEM131*** | Astro-M7 | magenta | -0.16 | -0.31 | 0.04 | -0.30 | -0.43 | 0.44 | -0.06 | 0.52 | -0.02 | 0.29 | 0.00 | -0.05 |
| ***LOC105758645*** | Astro-M7 | magenta | -0.30 | -0.02 | 0.60 | -0.46 | -0.67 | 0.15 | 0.04 | 0.79 | -0.13 | 0.19 | -0.02 | -0.08 |
| ***LOC115497609*** | Astro-M7 | magenta | -0.04 | 0.02 | 0.20 | -0.11 | -0.17 | 0.04 | 0.04 | 0.26 | -0.01 | 0.08 | 0.01 | 0.03 |
| ***LOC105758648*** | Astro-M7 | magenta | -0.21 | -0.20 | 0.24 | -0.40 | -0.51 | 0.33 | -0.05 | 0.64 | -0.04 | 0.29 | 0.01 | -0.07 |
| ***POU3F3*** | Astro-M7 | magenta | -0.05 | -0.06 | 0.11 | -0.13 | -0.18 | 0.12 | 0.00 | 0.26 | -0.00 | 0.12 | 0.03 | -0.01 |
| ***SH3RF3*** | Astro-M7 | magenta | -0.06 | -0.31 | 0.05 | -0.25 | -0.40 | 0.43 | 0.03 | 0.47 | 0.11 | 0.24 | 0.02 | 0.00 |
| ***DCUN1D2*** | Astro-M7 | magenta | -0.06 | -0.10 | 0.04 | -0.13 | -0.17 | 0.16 | -0.03 | 0.24 | -0.02 | 0.10 | 0.02 | -0.02 |
| ***BCOR*** | Astro-M7 | magenta | -0.01 | -0.05 | 0.07 | -0.05 | -0.14 | 0.12 | -0.00 | 0.19 | 0.01 | 0.13 | 0.03 | 0.03 |
| ***PLXNA4*** | Astro-M7 | magenta | -0.01 | -0.35 | 0.06 | -0.29 | -0.50 | 0.48 | 0.09 | 0.54 | 0.22 | 0.30 | 0.03 | 0.03 |
| ***CELF2*** | Astro-M7 | magenta | -0.11 | -0.00 | 0.31 | -0.25 | -0.30 | 0.09 | 0.03 | 0.48 | 0.00 | 0.18 | 0.04 | -0.01 |
| ***CACNA2D1*** | Astro-M7 | magenta | 0.09 | -0.11 | 0.14 | -0.13 | -0.19 | 0.12 | 0.15 | 0.24 | 0.17 | 0.16 | 0.04 | 0.08 |
| ***PLXNB2*** | Astro-M7 | magenta | -0.15 | -0.15 | 0.23 | -0.21 | -0.39 | 0.25 | 0.01 | 0.49 | -0.07 | 0.12 | -0.00 | -0.03 |
| ***ST7*** | Astro-M7 | magenta | -0.02 | -0.21 | 0.03 | -0.22 | -0.25 | 0.32 | 0.02 | 0.35 | 0.09 | 0.19 | 0.02 | 0.03 |
| ***SLC38A1*** | Astro-M7 | magenta | 0.10 | -0.13 | 0.03 | -0.09 | -0.15 | 0.18 | 0.11 | 0.17 | 0.16 | 0.12 | 0.03 | 0.09 |
| ***SLC6A15*** | Astro-M7 | magenta | 0.06 | -0.06 | 0.14 | -0.10 | -0.16 | 0.11 | 0.11 | 0.21 | 0.11 | 0.08 | 0.04 | 0.07 |
| ***PTPRN2*** | Astro-M7 | magenta | 0.01 | -0.16 | 0.35 | -0.19 | -0.52 | 0.21 | 0.21 | 0.56 | 0.17 | 0.22 | 0.03 | 0.07 |
| ***LOC121469345*** | Astro-M7 | magenta | 0.01 | -0.04 | 0.14 | -0.07 | -0.17 | 0.08 | 0.05 | 0.21 | 0.04 | 0.08 | 0.03 | 0.04 |
| ***CDK6*** | Astro-M7 | magenta | -0.05 | -0.02 | 0.19 | -0.12 | -0.19 | 0.11 | 0.01 | 0.33 | -0.05 | 0.12 | -0.00 | 0.03 |
| ***GLCCI1*** | Astro-M7 | magenta | -0.15 | -0.17 | 0.10 | -0.27 | -0.34 | 0.29 | -0.06 | 0.45 | -0.05 | 0.10 | 0.00 | -0.04 |
| ***ICA1*** | Astro-M7 | magenta | 0.01 | -0.13 | 0.02 | -0.12 | -0.16 | 0.22 | 0.03 | 0.22 | 0.06 | 0.12 | 0.03 | 0.03 |

| Gene | Module | Color | kME grey | kME Astro- M1 | kME Astro- M2 | kME Astro- M3 | kME Astro-M4 | kME Astro-M5 | kME Astro-M6 | kME Astro-M7 | kME Astro-M8 | kME Astro-M9 | kME Astro-M10 | kME Astro-M11 |
|---|---|---|---|---|---|---|---|---|---|---|---|---|---|---|
| ***NXPH1*** | Astro-M7 | magenta | -0.05 | -0.21 | 0.05 | -0.26 | -0.29 | 0.32 | -0.01 | 0.37 | 0.07 | 0.19 | 0.01 | 0.00 |
| ***DGKB*** | Astro-M7 | magenta | -0.01 | 0.10 | 0.26 | -0.10 | -0.18 | 0.05 | 0.06 | 0.30 | 0.01 | 0.07 | 0.04 | 0.06 |
| ***TOX*** | Astro-M7 | magenta | -0.02 | -0.13 | 0.14 | -0.11 | -0.31 | 0.25 | 0.05 | 0.37 | 0.05 | 0.17 | 0.01 | 0.06 |
| ***KCNQ5*** | Astro-M7 | magenta | -0.18 | -0.01 | 0.46 | -0.32 | -0.47 | 0.09 | 0.04 | 0.61 | -0.07 | 0.20 | -0.01 | -0.02 |
| ***CD24*** | Astro-M7 | magenta | -0.08 | -0.06 | 0.18 | -0.12 | -0.23 | 0.08 | 0.02 | 0.30 | -0.03 | 0.07 | 0.01 | -0.02 |
| ***FAM53A*** | Astro-M7 | magenta | -0.21 | -0.33 | 0.08 | -0.30 | -0.50 | 0.48 | -0.10 | 0.58 | -0.07 | 0.20 | -0.01 | -0.09 |
| ***TMEM131L*** | Astro-M7 | magenta | 0.01 | -0.19 | 0.05 | -0.15 | -0.24 | 0.24 | 0.07 | 0.27 | 0.10 | 0.09 | 0.02 | 0.03 |
| ***FBXW7*** | Astro-M7 | magenta | -0.05 | -0.09 | 0.13 | -0.15 | -0.24 | 0.20 | 0.01 | 0.33 | -0.01 | 0.12 | 0.02 | 0.02 |
| ***MAML3*** | Astro-M7 | magenta | -0.09 | -0.25 | -0.01 | -0.21 | -0.34 | 0.42 | -0.03 | 0.40 | 0.03 | 0.16 | 0.03 | -0.01 |
| ***KCNC1*** | Astro-M7 | magenta | 0.09 | -0.14 | 0.14 | -0.13 | -0.26 | 0.24 | 0.17 | 0.32 | 0.21 | 0.19 | 0.04 | 0.08 |
| ***MEIS2*** | Astro-M7 | magenta | -0.06 | -0.09 | 0.30 | -0.27 | -0.34 | 0.14 | 0.12 | 0.45 | 0.05 | 0.21 | -0.01 | 0.03 |
| ***LOC101234101*** | Astro-M7 | magenta | -0.03 | -0.01 | 0.18 | -0.10 | -0.19 | 0.07 | 0.04 | 0.26 | 0.01 | 0.07 | 0.02 | 0.03 |
| ***FOXG1*** | Astro-M7 | magenta | -0.25 | -0.08 | 0.45 | -0.34 | -0.61 | 0.19 | 0.04 | 0.67 | -0.09 | 0.18 | -0.01 | -0.06 |
| ***BCL11B*** | Astro-M7 | magenta | 0.01 | -0.34 | -0.02 | -0.24 | -0.35 | 0.43 | 0.07 | 0.38 | 0.20 | 0.32 | 0.03 | 0.04 |
| ***SYNE2*** | Astro-M7 | magenta | 0.03 | -0.15 | 0.02 | -0.14 | -0.19 | 0.27 | 0.02 | 0.24 | 0.08 | 0.14 | 0.06 | 0.06 |
| ***KCNT2*** | Astro-M7 | magenta | 0.03 | -0.17 | 0.03 | -0.13 | -0.23 | 0.28 | 0.04 | 0.27 | 0.11 | 0.17 | 0.04 | 0.06 |
| ***SOX2*** | Astro-M7 | magenta | -0.20 | 0.05 | 0.24 | -0.25 | -0.21 | 0.07 | -0.09 | 0.37 | -0.17 | 0.05 | 0.03 | -0.07 |
| ***IGDCC3*** | Astro-M7 | magenta | -0.11 | -0.14 | 0.06 | -0.21 | -0.27 | 0.26 | -0.06 | 0.37 | -0.02 | 0.15 | 0.02 | -0.05 |
| ***MAD1L1*** | Astro-M7 | magenta | -0.02 | -0.04 | 0.20 | -0.15 | -0.22 | 0.12 | 0.04 | 0.33 | 0.01 | 0.24 | 0.03 | 0.06 |
| ***MN1*** | Astro-M7 | magenta | -0.07 | -0.05 | 0.19 | -0.19 | -0.23 | 0.10 | 0.00 | 0.36 | 0.00 | 0.18 | 0.01 | 0.00 |
| ***PRDM16*** | Astro-M7 | magenta | -0.13 | 0.08 | 0.36 | -0.21 | -0.25 | 0.01 | 0.02 | 0.41 | -0.10 | 0.14 | 0.00 | 0.00 |
| ***MAP1B*** | Astro-M7 | magenta | 0.02 | -0.02 | 0.22 | -0.08 | -0.23 | 0.10 | 0.10 | 0.30 | 0.07 | 0.07 | -0.11 | 0.13 |
| ***ASCL1*** | Astro-M7 | magenta | -0.09 | -0.23 | -0.03 | -0.18 | -0.25 | 0.33 | -0.07 | 0.32 | -0.02 | 0.22 | 0.02 | -0.03 |
| ***NOTCH1*** | Astro-M7 | magenta | -0.13 | -0.14 | 0.04 | -0.23 | -0.20 | 0.24 | -0.10 | 0.34 | -0.09 | 0.16 | 0.03 | -0.05 |
| ***CCND2*** | Astro-M7 | magenta | -0.09 | -0.06 | 0.10 | -0.16 | -0.16 | 0.12 | -0.05 | 0.27 | -0.06 | 0.05 | 0.01 | -0.03 |
| ***FANCC*** | Astro-M7 | magenta | -0.03 | -0.07 | 0.10 | -0.15 | -0.17 | 0.12 | -0.00 | 0.25 | -0.03 | 0.19 | -0.07 | 0.06 |
| ***HMGA2*** | Astro-M7 | magenta | -0.08 | 0.08 | 0.27 | -0.13 | -0.19 | 0.02 | 0.01 | 0.33 | -0.05 | 0.13 | 0.01 | -0.00 |

**Supplemental Table 10: ShinyGO enrichment for Astro-M7 (human annotation).** Gene Ontology (GO) biological process enrichment of human orthologs corresponding to Astro-M7 module genes, performed using ShinyGO. Columns: Pathway (GO term ID and name), Enrichment FDR (Benjamini–Hochberg–adjusted hypergeometric test q-value), nGenes (number of input genes overlapping the term), Pathway Genes (total genes annotated to the term), Fold Enrichment (overrepresentation of input genes versus background), and Genes (overlapping input genes).

| Pathway | Enrichment FDR | nGenes | Pathway Genes | Fold Enrichment | Genes |
|---|---|---|---|---|---|
| **GO:0097150 Neuronal stem cell population maintenance** | 1.14E-02 | 3 | 27 | 34.27 | *FANCC, NOTCH1, SOX2* |
| **GO:2000179 Pos. reg. of neural precursor cell proliferation** | 1.02E-02 | 4 | 66 | 18.28 | *ASCL1, NOTCH1, FOXG1, TOX* |
| **GO:0033077 T cell differentiation in thymus** | 1.2E-02 | 4 | 92 | 16.22 | *CDK6, BCL11B, TOX, TMEM131L* |
| **GO:0045665 Neg. reg. of neuron differentiation** | 1.2E-02 | 4 | 76 | 15.96 | *ASCL1, NOTCH1, FOXG1, SOX2* |
| **GO:0061351 Neural precursor cell proliferation** | 3.95E-03 | 6 | 176 | 10.62 | *ASCL1, NOTCH1, FOXG1, PLXNB2, TOX, POU3F3* |
| **GO:0045664 Reg. of neuron differentiation** | 6.25E-03 | 6 | 198 | 9.48 | *ASCL1, BCL11B, MAP1B, NOTCH1, FOXG1, SOX2* |
| **GO:0021537 Telencephalon development** | 1.03E-03 | 8 | 301 | 8.90 | *SYNE2, BCL11B, KCNC1, ASCL1, FOXG1, POU3F3, PLXNA4, CDK6* |
| **GO:0030900 Forebrain development** | 4.79E-05 | 11 | 442 | 8.25 | *SOX2, SYNE2, BCL11B, KCNC1, ASCL1, NOTCH1, FOXG1, TOX, POU3F3, PLXNA4, CDK6* |
| **GO:0007420 Brain development** | 4.79E-05 | 14 | 795 | 6.04 | *MEIS2, SOX2, SYNE2, BCL11B, KCNC1, ASCL1, NOTCH1, FANCC, FOXG1, PLXNB2, TOX, POU3F3, PLXNA4, CDK6* |
| **GO:0060322 Head development** | 4.79E-05 | 14 | 848 | 5.65 | *MEIS2, SOX2, SYNE2, BCL11B, KCNC1, ASCL1, NOTCH1, FANCC, FOXG1, PLXNB2, TOX, POU3F3, PLXNA4, CDK6* |
| **GO:0060284 Reg. of cell development** | 3.1E-03 | 11 | 915 | 4.71 | *ASCL1, PLXNB2, PLXNA4, CDK6, FBXW7, MAP1B, MEIS2, NOTCH1, FOXG1, TOX, TMEM131L* |
| **GO:0007417 Central nervous system development** | 9.38E-04 | 14 | 1121 | 4.32 | *MEIS2, SOX2, SYNE2, BCL11B, KCNC1, ASCL1, NOTCH1, FANCC, FOXG1, PLXNB2, TOX, POU3F3, PLXNA4, CDK6* |
| **GO:0008284 Pos. reg. of cell population proliferation** | 1.2E-02 | 10 | 1063 | 4.01 | *MEIS2, NOTCH1, HMGA2, CD24, ASCL1, FOXG1, TOX, POU3F3, CDK6, CCND2* |

| Pathway | Enrichment FDR | nGenes | Pathway Genes | Fold Enrichment | Genes |
|---|---|---|---|---|---|
| **GO:0045595 Reg. of cell differentiation** | 1.03E-03 | 16 | 1703 | 3.60 | *ASCL1, PLXNB2, PLXNA4, CDK6, NOTCH1, FBXW7, BCL11B, MAP1B, MEIS2, FOXG1, SOX2, TOX, TMEM131L, CD24, EPHA3, ST7* |
| **GO:0042127 Reg. of cell population proliferation** | 1.08E-03 | 16 | 1851 | 3.54 | *MEIS2, NOTCH1, HMGA2, MN1, SOX2, CD24, MAD1L1, CDK6, BCL11B, ASCL1, FOXG1, TOX, POU3F3, CCND2, TMEM131L, FBXW7* |
| **GO:2000026 Reg. of multicellular organismal development** | 4.28E-03 | 14 | 1531 | 3.48 | *ASCL1, PLXNB2, PLXNA4, CDK6, FBXW7, MAP1B, MEIS2, NOTCH1, FOXG1, TOX, TMEM131L, BCOR, CD24, HMGA2* |
| **GO:0045892 Neg. reg. of dna-templated transcription** | 1.2E-02 | 12 | 1401 | 3.39 | *SOX2, ZBTB20, BCOR, ASCL1, PRDM16, POU2F1, HMGA2, FOXG1, CDK6, NOTCH1, POU3F3, MEIS2* |
| **GO:0008283 Cell population proliferation** | 2.25E-03 | 17 | 2185 | 3.13 | *MEIS2, NOTCH1, HMGA2, MN1, SOX2, CD24, MAD1L1, CDK6, BCL11B, ASCL1, FOXG1, PLXNB2, TOX, POU3F3, CCND2, TMEM131L, FBXW7* |
| **GO:0050793 Reg. of developmental proc.** | 1.04E-02 | 18 | 2666 | 2.56 | *ASCL1, PLXNB2, PLXNA4, CDK6, NOTCH1, FBXW7, BCL11B, MAP1B, MEIS2, FOXG1, SOX2, TOX, TMEM131L, HMGA2, BCOR, CD24, EPHA3, ST7* |
| **GO:0051239 Reg. of multicellular organismal proc.** | 1.14E-02 | 19 | 3355 | 2.41 | *ASCL1, ZBTB20, PLXNB2, PLXNA4, CDK6, CD24, MAD1L1, FBXW7, MAP1B, MEIS2, NOTCH1, FOXG1, TOX, TMEM131L, HMGA2, CACNA2D1, BCOR, PRDM16, CELF2* |

**Supplemental Table 11: ShinyGO enrichment for Astro-M7 (zebra finch annotation).** Gene Ontology (GO) biological process enrichment of human orthologs corresponding to Astro-M7 module genes, performed using ShinyGO. Columns: Pathway (GO term ID and name), Enrichment FDR (Benjamini–Hochberg–adjusted hypergeometric test q-value), nGenes (number of input genes overlapping the term), Pathway Genes (total genes annotated to the term), Fold Enrichment (overrepresentation of input genes versus background), and Genes (overlapping input genes).

| Pathway | Enrichment FDR | nGenes | Pathway Genes | Fold Enrichment | Genes |
|---|---|---|---|---|---|
| **GO:0007221 Pos. reg. of transcription of notch receptor target** | 1.13E-02 | 2 | 4 | 146.66 | *MAML3, NOTCH1* |
| **GO:0021902 Commitment of neuronal cell to specific neuron type in forebrain** | 1.56E-02 | 2 | 6 | 97.77 | *BCL11B, ASCL1* |
| **GO:0021879 Forebrain neuron differentiation** | 1.38E-02 | 3 | 31 | 31.43 | *FOXG1, BCL11B, ASCL1* |
| **GO:0060563 Neuroepithelial cell differentiation** | 1.38E-02 | 3 | 31 | 29.33 | *MAP1B, NOTCH1, ASCL1* |
| **GO:0048663 Neuron fate commitment** | 8.42E-03 | 4 | 54 | 22.14 | *ASCL1, FOXG1, NOTCH1, BCL11B* |
| **GO:0021953 Central nervous system neuron differentiation** | 1.38E-02 | 5 | 141 | 10.78 | *FOXG1, PLXNA4, TOX, BCL11B, ASCL1* |
| **GO:0021537 Telencephalon development** | 1.56E-02 | 5 | 161 | 9.40 | *FOXG1, PLXNA4, BCL11B, SYNE2, ASCL1* |
| **GO:0030900 Forebrain development** | 5.62E-03 | 7 | 241 | 8.85 | *FOXG1, PLXNA4, NOTCH1, TOX, BCL11B, SYNE2, ASCL1* |
| **GO:0007409 Axonogenesis** | 5.62E-03 | 7 | 267 | 8.66 | *PLXNA4, MAP1B, NOTCH1, BCL11B, EPHA3, PLXNB2, FOXG1* |
| **GO:0061564 Axon development** | 7.98E-03 | 7 | 292 | 7.93 | *PLXNA4, MAP1B, NOTCH1, BCL11B, EPHA3, PLXNB2, FOXG1* |
| **GO:0007420 Brain development** | 5.62E-03 | 9 | 404 | 6.82 | *FOXG1, FANCC, PLXNA4, NOTCH1, TOX, BCL11B, SYNE2, ASCL1, PLXNB2* |
| **GO:0048667 Cell morphogenesis involved in neuron differentiation** | 1.13E-02 | 7 | 341 | 6.73 | *PLXNA4, MAP1B, NOTCH1, BCL11B, EPHA3, PLXNB2, FOXG1* |
| **GO:0060322 Head development** | 5.62E-03 | 9 | 438 | 6.33 | *FOXG1, FANCC, PLXNA4, NOTCH1, TOX, BCL11B, SYNE2, ASCL1, PLXNB2* |
| **GO:0048812 Neuron projection morphogenesis** | 1.38E-02 | 7 | 364 | 6.28 | *PLXNA4, MAP1B, NOTCH1, BCL11B, EPHA3, PLXNB2, FOXG1* |
| **GO:0060284 Reg. of cell development** | 8.45E-03 | 8 | 396 | 6.18 | *MEIS2, TMEM131L, NOTCH1, TOX, PLXNB2, FBXW7, ASCL1, FOXG1* |
| **GO:0120039 Plasma membrane bounded cell projection morphogenesis** | 1.38E-02 | 7 | 373 | 6.11 | *PLXNA4, MAP1B, NOTCH1, BCL11B, EPHA3, PLXNB2, FOXG1* |
| **GO:0048858 Cell projection morphogenesis** | 1.38E-02 | 7 | 375 | 6.07 | *PLXNA4, MAP1B, NOTCH1, BCL11B, EPHA3, PLXNB2, FOXG1* |
| **GO:0007417 Central nervous system development** | 1.13E-02 | 9 | 549 | 5.02 | *FOXG1, FANCC, PLXNA4, NOTCH1, TOX, BCL11B, SYNE2, ASCL1, PLXNB2* |

| Pathway | Enrichment FDR | nGenes | Pathway Genes | Fold Enrichment | Genes |
|---|---|---|---|---|---|
| **GO:0048666 Neuron development** | 1.38E-02 | 9 | 663 | 4.44 | *PLXNB2, ASCL1, FOXG1, PLXNA4, MAP1B, NOTCH1, TOX, BCL11B, EPHA3* |
| **GO:0042127 Reg. of cell population proliferation** | 1.57E-02 | 10 | 809 | 3.81 | *MN1, ASCL1, MEIS2, TMEM131L, NOTCH1, MAD1L1, TOX, BCL11B, FOXG1, CCND2* |

**Supplemental Table 12: Astrocyte differential accessibility peak-to-gene annotations**. Differentially accessible (DA) ATAC peaks in astrocytes comparing heat call versus control playback (MAST; FDR < 0.05 and |log2FC| ≥ 0.25). Peaks were annotated to the nearest gene using Signac ClosestFeature, and the table reports DA statistics together with gene proximity information for each peak. Columns: peak (genomic coordinates), gene_name (nearest gene), distance (bp from peak to the nearest gene transcription start site; 0 indicates promoter overlap), Gene Biotype, average log2FC (Heat Call/Control), FDR (FDR-adjusted p-value), Fraction 1 (fraction of heat call astrocytes with accessible chromatin at this peak), and Fraction 2 (fraction of control astrocytes with accessible chromatin at this peak).

| Peak | Gene | Distance | Gene Biotype | Average log2FC | FDR | Fraction 1 | Fraction 2 |
|---|---|---|---|---|---|---|---|
| **NW-024545321.1-45497-46517** | *LOC121468926* | 0 | NA | 0.98 | 2.26E-48 | 0.15 | 0.07 |
| **NW-024545320.1-20536-21507** | *LOC121468922* | 1716 | Protein coding | 0.96 | 1.18E-43 | 0.15 | 0.07 |
| **NW-024545308.1-66979-67836** | *LOC121468895* | 0 | NA | 0.59 | 4.21E-42 | 0.23 | 0.15 |
| **NC-044225.2-17400862-17401778** | *YIPF5* | 0 | NA | 0.46 | 2.02E-17 | 0.20 | 0.14 |
| **NC-044212.2-28422-29337** | *LOC116809227* | 0 | NA | 0.29 | 2.78E-17 | 0.26 | 0.21 |
| **NC-044224.2-13659761-13660705** | *DCP1A* | 0 | NA | 0.30 | 8.64E-14 | 0.12 | 0.09 |
| **NC-044211.2-76822446-76823394** | *POGLUT2* | 0 | NA | 0.39 | 8.73E-14 | 0.11 | 0.08 |
| **NC-044211.2-69373996-69374901** | *FDX1* | 0 | NA | 0.30 | 1.24E-13 | 0.15 | 0.12 |
| **NC-054767.1-987391-988188** | *RING1* | 0 | NA | 0.25 | 5.56E-13 | 0.23 | 0.18 |
| **NC-044243.2-411703-412610** | *NFIX* | 0 | NA | 0.33 | 2.47E-12 | 0.15 | 0.11 |
| **NC-044223.2-10968813-10969745** | *CTU2* | 0 | NA | 0.25 | 4.31E-12 | 0.27 | 0.22 |
| **NC-044232.2-12335224-12336167** | *BMP7* | 104497 | NA | 0.31 | 6.15E-12 | 0.19 | 0.15 |
| **NC-044225.2-7428128-7429749** | *PDLIM4* | 0 | NA | 0.33 | 1.79E-11 | 0.17 | 0.13 |
| **NC-044224.2-14070501-14071510** | *LOC100219242* | 0 | NA | 0.36 | 1.89E-11 | 0.21 | 0.16 |
| **NC-054767.1-2077204-2078128** | *LOC116807445* | 0 | NA | 0.25 | 2.03E-11 | 0.18 | 0.15 |
| **NC-044214.2-31148384-31149284** | *MYO6* | 0 | NA | 0.30 | 5.07E-11 | 0.18 | 0.14 |
| **NC-044217.2-60043656-60044566** | *KLHDC1* | 0 | NA | 0.25 | 5.80E-11 | 0.20 | 0.16 |
| **NC-044240.2-753642-754515** | *NMRK2* | 0 | NA | 0.26 | 9.14E-11 | 0.18 | 0.14 |
| **NC-044223.2-6113164-6114105** | *CSNK2A2* | 0 | NA | 0.27 | 1.63E-10 | 0.13 | 0.10 |
| **NC-044221.2-2356666-2357648** | *KLHL6* | 0 | NA | 0.41 | 2.07E-10 | 0.11 | 0.08 |

| Peak | Gene | Distance | Gene Biotype | Average log2FC | FDR | Fraction 1 | Fraction 2 |
|---|---|---|---|---|---|---|---|
| **NC-044218.2-29866478-29867381** | *LOC121470143* | 0 | NA | 0.27 | 4.87E-10 | 0.20 | 0.16 |
| **NC-044213.2-4296865-4297771** | *LOC121469361* | 1536 | NA | 0.29 | 6.67E-10 | 0.21 | 0.17 |
| **NC-044211.2-32894826-32895759** | *ANKRD42* | 0 | NA | 0.31 | 9.85E-10 | 0.19 | 0.15 |
| **NC-044213.2-14401521-14402425** | *ITGB1* | 0 | NA | 0.29 | 2.74E-09 | 0.15 | 0.12 |
| **NC-044218.2-21823949-21824886** | *MMS19* | 0 | NA | 0.27 | 3.08E-09 | 0.18 | 0.14 |
| **NC-044213.2-52510853-52511824** | *GLI3* | 0 | NA | 0.32 | 3.90E-09 | 0.11 | 0.08 |
| **NC-044222.2-7647270-7648233** | *CCPG1* | 0 | NA | 0.28 | 3.98E-09 | 0.12 | 0.10 |
| **NC-044212.2-887423-888289** | *TSPAN33* | 0 | NA | 0.52 | 4.10E-09 | 0.10 | 0.07 |
| **NC-044217.2-57031944-57032899** | *DACT1* | 64411 | NA | 0.46 | 4.65E-09 | 0.12 | 0.08 |
| **NC-044236.2-7474186-7475117** | *RPS25* | 0 | NA | 0.25 | 5.42E-09 | 0.13 | 0.11 |
| **NC-054768.1-15964-16770** | *OSGEP* | 0 | NA | 0.29 | 5.74E-09 | 0.12 | 0.10 |
| **NC-044213.2-21070852-21071780** | *MINDY3* | 0 | NA | 0.34 | 6.12E-09 | 0.14 | 0.11 |
| **NC-044218.2-35581656-35582554** | *LOC121470206* | 0 | NA | 0.43 | 6.82E-09 | 0.15 | 0.11 |
| **NC-044242.2-558840-559580** | *SP1* | 1365 | Protein coding | 0.26 | 6.84E-09 | 0.13 | 0.10 |
| **NC-044213.2-65273066-65274024** | *LOC100190710* | 0 | NA | 0.40 | 8.23E-09 | 0.14 | 0.10 |
| **NC-044213.2-71534722-71535626** | *ATP9B* | 0 | NA | 0.27 | 1.09E-08 | 0.19 | 0.15 |
| **NC-044241.2-6521867-6522814** | *CDC42SE2* | 4757 | Protein coding | 0.52 | 2.84E-08 | 0.10 | 0.07 |
| **NC-044226.2-3633487-3634411** | *MAD1L1* | 0 | NA | 0.29 | 3.06E-08 | 0.16 | 0.13 |
| **NC-044216.2-7284408-7285268** | *MBNL3* | 0 | NA | 0.29 | 4.43E-08 | 0.15 | 0.12 |
| **NC-044212.2-53563924-53564841** | *LOC115493394* | 58233 | NA | 0.28 | 8.43E-08 | 0.15 | 0.12 |
| **NC-044217.2-6527953-6528913** | *TPCN2* | 0 | NA | 0.29 | 9.41E-08 | 0.10 | 0.08 |
| **NC-044211.2-56143572-56144485** | *RGCC* | 0 | NA | 0.26 | 1.00E-07 | 0.12 | 0.10 |
| **NC-044219.2-19333210-19334116** | *LOC100232284* | 53423 | NA | 0.29 | 1.15E-07 | 0.16 | 0.13 |
| **NC-044214.2-48930368-48931338** | *MCM9* | 0 | NA | 0.34 | 1.21E-07 | 0.14 | 0.11 |
| **NC-044220.2-22317561-22318496** | *ZBTB41* | 0 | NA | 0.26 | 1.27E-07 | 0.15 | 0.12 |
| **NC-044213.2-122726419-122727375** | *PAG1* | 0 | NA | 0.29 | 1.31E-07 | 0.10 | 0.08 |

| Peak | Gene | Distance | Gene Biotype | Average log2FC | FDR | Fraction 1 | Fraction 2 |
|---|---|---|---|---|---|---|---|
| **NC-044226.2-3834330-3835237** | *LOC115497219* | 689 | NA | 0.26 | 1.35E-07 | 0.11 | 0.09 |
| **NC-044219.2-36304568-36305484** | *RND3* | 3727 | NA | 0.27 | 1.51E-07 | 0.21 | 0.17 |
| **NC-044211.2-63720419-63721359** | *RFC3* | 25621 | NA | 0.41 | 1.76E-07 | 0.12 | 0.08 |
| **NC-044214.2-69141268-69142236** | *IRF2BP2* | 16497 | NA | 0.28 | 1.98E-07 | 0.12 | 0.10 |
| **NC-044216.2-6978207-6979155** | *GPC4* | 0 | NA | 0.26 | 2.06E-07 | 0.15 | 0.12 |
| **NC-044214.2-105427333-105428264** | *PLB1* | 4687 | NA | 0.29 | 2.06E-07 | 0.12 | 0.10 |
| **NC-044239.2-2335986-2336950** | *KAT2A* | 0 | NA | 0.26 | 2.18E-07 | 0.16 | 0.13 |
| **NC-044215.2-45826249-45827231** | *CFI* | 0 | NA | 0.31 | 2.43E-07 | 0.14 | 0.11 |
| **NC-044217.2-59992239-59993042** | *NEMF* | 0 | NA | 0.25 | 2.93E-07 | 0.12 | 0.10 |
| **NC-044220.2-19014681-19015597** | *LOC100218707* | 0 | NA | 0.28 | 3.73E-07 | 0.13 | 0.10 |
| **NC-044216.2-18984961-18985855** | *GDPD2* | 0 | NA | 0.25 | 4.42E-07 | 0.15 | 0.12 |
| **NC-044213.2-85790596-85791457** | *TMEM245* | 0 | NA | 0.29 | 4.46E-07 | 0.15 | 0.12 |
| **NC-044214.2-57697851-57698778** | *SYNE1* | 0 | NA | 0.31 | 4.59E-07 | 0.12 | 0.09 |
| **NC-044231.2-938966-939720** | *MSI2* | 0 | NA | 0.26 | 4.82E-07 | 0.12 | 0.09 |
| **NC-044229.2-5752152-5753110** | *PKN3* | 0 | NA | 0.29 | 5.12E-07 | 0.12 | 0.09 |
| **NC-044215.2-7182123-7183064** | *LOC115494888* | 50105 | NA | 0.34 | 5.23E-07 | 0.12 | 0.09 |
| **NC-044211.2-97717633-97718559** | *NHS* | 0 | NA | 0.34 | 5.40E-07 | 0.11 | 0.08 |
| **NC-044217.2-12713913-12714903** | *SOX6* | 0 | NA | 0.26 | 5.49E-07 | 0.20 | 0.16 |
| **NC-044213.2-25564872-25565718** | *ASNS* | 0 | NA | 0.31 | 5.81E-07 | 0.12 | 0.09 |
| **NC-044221.2-15988019-15988944** | *LOC115496426* | 0 | NA | 0.25 | 6.17E-07 | 0.16 | 0.13 |
| **NC-044224.2-19530203-19531185** | *WNT7A* | 15666 | NA | 0.35 | 6.24E-07 | 0.11 | 0.09 |
| **NC-044217.2-54096483-54097517** | *LOC116808502* | 49052 | NA | 0.30 | 6.34E-07 | 0.16 | 0.12 |
| **NC-044219.2-8993902-8994883** | *RPL37A* | 4792 | NA | 0.25 | 6.42E-07 | 0.19 | 0.16 |
| **NC-044212.2-15149965-15150987** | *CELSR1* | 0 | NA | 0.27 | 6.49E-07 | 0.11 | 0.08 |
| **NC-044214.2-80049774-80050726** | *TFB2M* | 0 | NA | 0.25 | 6.64E-07 | 0.11 | 0.09 |
| **NC-044225.2-12839238-12840139** | *MRPL22* | 0 | NA | 0.30 | 6.94E-07 | 0.10 | 0.08 |
| **NC-045028.1-17183846-17184780** | *LOC116806894* | 13796 | NA | 0.25 | 7.49E-07 | 0.11 | 0.09 |

| Peak | Gene | Distance | Gene Biotype | Average log2FC | FDR | Fraction 1 | Fraction 2 |
|---|---|---|---|---|---|---|---|
| **NC-044215.2-13207917-13208809** | *WDR1* | 0 | NA | 0.32 | 7.77E-07 | 0.17 | 0.13 |
| **NC-044214.2-112584940-112585861** | *MAP4K3* | 11192 | Protein coding | 0.27 | 8.15E-07 | 0.12 | 0.10 |
| **NC-044219.2-10524394-10525352** | *SLC4A3* | 0 | NA | 0.27 | 8.28E-07 | 0.12 | 0.09 |
| **NC-044214.2-93087974-93088895** | *LCLAT1* | 0 | NA | 0.31 | 8.51E-07 | 0.15 | 0.11 |
| **NC-044217.2-45867682-45868583** | *TRIP11* | 0 | NA | 0.26 | 9.22E-07 | 0.16 | 0.13 |
| **NC-044211.2-44450558-44451442** | *LOC116808177* | 22929 | NA | 0.29 | 9.52E-07 | 0.12 | 0.09 |
| **NC-044240.2-5276956-5277842** | *RPL36* | 0 | NA | 0.28 | 1.09E-06 | 0.15 | 0.12 |
| **NC-044214.2-60352717-60353694** | *TULP4* | 0 | NA | 0.29 | 1.26E-06 | 0.13 | 0.10 |
| **NC-044221.2-1449494-1450520** | *COPS9* | 0 | NA | 0.30 | 1.41E-06 | 0.15 | 0.12 |
| **NC-044214.2-55417955-55418882** | *SLC2A12* | 6424 | Protein coding | 0.26 | 1.47E-06 | 0.14 | 0.11 |
| **NC-044212.2-7068023-7069007** | *FAM107B* | 0 | NA | 0.26 | 1.50E-06 | 0.11 | 0.09 |
| **NC-044223.2-1586178-1587151** | *FTO* | 0 | NA | 0.27 | 1.66E-06 | 0.14 | 0.11 |
| **NC-044215.2-38520222-38521164** | *KLHL2* | 0 | NA | 0.26 | 1.67E-06 | 0.15 | 0.12 |
| **NC-044221.2-6824192-6825165** | *LIMS2* | 0 | NA | 0.27 | 1.70E-06 | 0.11 | 0.09 |
| **NC-044213.2-34507852-34508779** | *CREB5* | 0 | NA | 0.30 | 1.81E-06 | 0.10 | 0.08 |
| **NC-044224.2-514937-515878** | *SRGAP3* | 0 | NA | 0.25 | 1.81E-06 | 0.11 | 0.09 |
| **NC-044232.2-8385490-8386425** | *RGS19* | 0 | NA | 0.29 | 1.93E-06 | 0.10 | 0.08 |
| **NC-044217.2-51953918-51954853** | *TRMT61A* | 0 | NA | 0.26 | 2.10E-06 | 0.14 | 0.11 |
| **NC-044219.2-13605665-13606592** | *G6PC2* | 0 | NA | 0.25 | 2.11E-06 | 0.21 | 0.17 |
| **NC-044227.2-3475241-3476145** | *LOC121470755* | 0 | NA | 0.35 | 2.29E-06 | 0.13 | 0.10 |
| **NC-044213.2-137797025-137797965** | *TAF2* | 51 | Protein coding | 0.27 | 2.39E-06 | 0.11 | 0.09 |
| **NC-044216.2-5098322-5099223** | *LOC100223640* | 0 | NA | 0.26 | 2.61E-06 | 0.15 | 0.12 |
| **NC-044215.2-64617545-64618452** | *LOC100229648* | 29085 | Protein coding | 0.35 | 2.80E-06 | 0.11 | 0.08 |
| **NC-044212.2-2459332-2460211** | *PLXNA4* | 0 | NA | 0.33 | 2.97E-06 | 0.14 | 0.11 |
| **NC-054767.1-225388-226323** | *MGAT1* | 206 | NA | 0.27 | 3.17E-06 | 0.11 | 0.09 |

| Peak | Gene | Distance | Gene Biotype | Average log2FC | FDR | Fraction 1 | Fraction 2 |
|---|---|---|---|---|---|---|---|
| **NC-044225.2-14492818-14493723** | *KIAA1191* | 0 | NA | 0.26 | 3.34E-06 | 0.14 | 0.11 |
| **NC-044213.2-2635471-2636386** | *CCDC13* | 0 | NA | 0.27 | 3.60E-06 | 0.12 | 0.09 |
| **NC-044224.2-10000341-10001331** | *SLC41A3* | 0 | NA | 0.30 | 3.98E-06 | 0.11 | 0.09 |
| **NC-044213.2-42266174-42267108** | *POMGNT2* | 0 | NA | 0.26 | 4.01E-06 | 0.14 | 0.11 |
| **NC-044239.2-1551684-1552566** | *SRCIN1* | 0 | NA | 0.30 | 4.27E-06 | 0.14 | 0.11 |
| **NC-044211.2-24329324-24330265** | *LRRC58* | 0 | NA | 0.25 | 4.54E-06 | 0.14 | 0.11 |
| **NC-044230.2-8683652-8684606** | *CASKIN2* | 0 | NA | 0.25 | 4.56E-06 | 0.12 | 0.09 |
| **NC-044225.2-16849037-16849958** | *PWWP2A* | 0 | NA | 0.31 | 4.70E-06 | 0.15 | 0.12 |
| **NC-044232.2-3099637-3100576** | *SOGA1* | 3211 | Protein coding | 0.26 | 4.87E-06 | 0.13 | 0.10 |
| **NC-044226.2-15205040-15206015** | *RAB40C* | 0 | NA | 0.30 | 5.46E-06 | 0.12 | 0.09 |
| **NC-044218.2-29891305-29892202** | *LOC121470143* | 12305 | NA | 0.45 | 5.54E-06 | 0.10 | 0.07 |
| **NC-044225.2-1255609-1256312** | *CTNNA1* | 1700 | NA | 0.28 | 5.89E-06 | 0.11 | 0.09 |
| **NC-044222.2-5472197-5473143** | *MTMR10* | 0 | NA | 0.28 | 6.03E-06 | 0.16 | 0.13 |
| **NW-024545360.1-31611-32507** | *UXT* | 0 | NA | 0.28 | 7.05E-06 | 0.14 | 0.11 |
| **NC-044213.2-73360905-73361647** | *TNS3* | 0 | NA | 0.29 | 7.21E-06 | 0.14 | 0.11 |
| **NC-044215.2-64534876-64535764** | *LOC115495184* | 0 | NA | 0.27 | 7.57E-06 | 0.13 | 0.11 |
| **NC-044235.2-2973106-2974057** | *GRHL3* | 0 | NA | 0.29 | 7.96E-06 | 0.11 | 0.09 |
| **NC-044212.2-1944777-1945725** | *EXOC4* | 0 | NA | 0.38 | 8.61E-06 | 0.10 | 0.08 |
| **NC-044220.2-30420320-30421245** | *LOC100223828* | 0 | NA | 0.28 | 8.92E-06 | 0.11 | 0.09 |
| **NC-044211.2-88931459-88932409** | *SOX1* | 14105 | NA | 0.28 | 9.63E-06 | 0.12 | 0.09 |
| **NC-044222.2-10467202-10468134** | *SPPL2A* | 0 | NA | 0.33 | 1.00E-05 | 0.11 | 0.08 |
| **NC-044219.2-11149653-11150553** | *PSMD14* | 0 | NA | 0.27 | 1.02E-05 | 0.13 | 0.10 |
| **NC-044215.2-29303027-29303988** | *MTUS1* | 0 | NA | 0.28 | 1.11E-05 | 0.12 | 0.09 |
| **NC-044214.2-35889085-35890075** | *GABRR1* | 0 | NA | 0.29 | 1.16E-05 | 0.12 | 0.10 |
| **NC-044213.2-128473693-128474567** | *SDC2* | 0 | NA | 0.29 | 1.44E-05 | 0.13 | 0.10 |
| **NC-044230.2-323222-324134** | *KCNJ2* | 26104 | NA | 0.32 | 1.60E-05 | 0.10 | 0.08 |

| Peak | Gene | Distance | Gene Biotype | Average log2FC | FDR | Fraction 1 | Fraction 2 |
|---|---|---|---|---|---|---|---|
| **NC-044218.2-29655267-29656200** | *LOC121470203* | 0 | NA | 0.28 | 2.20E-05 | 0.13 | 0.10 |
| **NC-044218.2-19752390-19753291** | *LOC116808551* | 0 | NA | 0.29 | 2.21E-05 | 0.10 | 0.08 |
| **NC-044223.2-3950062-3951008** | *LOC121470587* | 14597 | NA | 0.27 | 2.40E-05 | 0.11 | 0.09 |
| **NC-044211.2-8450854-8451728** | *NRIP1* | 0 | NA | 0.26 | 2.84E-05 | 0.15 | 0.12 |
| **NC-044211.2-105604141-105605058** | *OTC* | 10923 | NA | 0.32 | 2.84E-05 | 0.12 | 0.09 |
| **NC-044225.2-14725483-14726405** | *PRELID1* | 0 | NA | 0.26 | 2.97E-05 | 0.14 | 0.12 |
| **NC-044221.2-4115584-4116444** | *UBE2G2* | 0 | NA | 0.33 | 3.14E-05 | 0.12 | 0.09 |
| **NC-044215.2-59152453-59153438** | *LOC105759261* | 12086 | NA | 0.29 | 3.22E-05 | 0.12 | 0.09 |
| **NC-044220.2-18966786-18967740** | *SLC30A7* | 0 | NA | 0.26 | 3.72E-05 | 0.14 | 0.12 |
| **NC-044220.2-933091-933984** | *CTH* | 0 | NA | 0.29 | 3.80E-05 | 0.13 | 0.11 |
| **NC-044220.2-15481669-15482605** | *LOC115496205* | 0 | NA | 0.34 | 4.00E-05 | 0.13 | 0.10 |
| **NC-044214.2-3255574-3256458** | *LOC101234062* | 0 | NA | 0.25 | 4.12E-05 | 0.11 | 0.09 |
| **NC-044223.2-5422492-5423449** | *TMEM208* | 0 | NA | 0.29 | 4.68E-05 | 0.11 | 0.09 |
| **NC-044233.2-2821132-2822039** | *PRDM16* | 0 | NA | 0.25 | 5.46E-05 | 0.13 | 0.11 |
| **NC-044214.2-84900252-84901216** | *ACYP2* | 0 | NA | 0.31 | 5.50E-05 | 0.11 | 0.08 |
| **NC-044211.2-36650647-36651597** | *LOC121469757* | 0 | NA | 0.26 | 5.71E-05 | 0.10 | 0.08 |
| **NC-044217.2-28105-29020** | *LUZP2* | 0 | NA | 0.31 | 5.76E-05 | 0.12 | 0.09 |
| **NC-044230.2-8002407-8003660** | *RHBDF2* | 0 | NA | 0.28 | 5.94E-05 | 0.12 | 0.09 |
| **NC-044219.2-10128308-10129169** | *CFAP65* | 0 | NA | 0.31 | 6.46E-05 | 0.10 | 0.08 |
| **NC-044211.2-70030075-70031029** | *LOC115495774* | 35476 | NA | 0.28 | 6.49E-05 | 0.11 | 0.09 |
| **NC-044220.2-22411576-22412495** | *KCNT2* | 0 | NA | 0.28 | 6.83E-05 | 0.12 | 0.10 |
| **NC-044211.2-96773600-96774561** | *ASB9* | 26194 | NA | 0.31 | 7.26E-05 | 0.10 | 0.08 |
| **NC-044220.2-24164649-24165607** | *LOC100220051* | 0 | NA | 0.26 | 7.47E-05 | 0.11 | 0.09 |
| **NC-044226.2-3984989-3986058** | *IQCE* | 0 | NA | 0.25 | 7.76E-05 | 0.10 | 0.08 |
| **NC-044219.2-37542649-37543666** | *ACVR1* | 0 | NA | 0.27 | 9.30E-05 | 0.15 | 0.12 |
| **NC-044232.2-7216853-7217711** | *LOC101233889* | 546 | NA | 0.26 | 9.33E-05 | 0.12 | 0.10 |
| **NC-044215.2-66066597-66067516** | *AP1AR* | 0 | NA | 0.25 | 9.36E-05 | 0.11 | 0.09 |

| Peak | Gene | Distance | Gene Biotype | Average log2FC | FDR | Fraction 1 | Fraction 2 |
|---|---|---|---|---|---|---|---|
| **NC-044214.2-39916754-39917657** | *POU3F2* | 1857 | Protein coding | 0.29 | 9.77E-05 | 0.15 | 0.12 |
| **NC-044229.2-9338214-9339100** | *LOC100225943* | 0 | NA | 0.29 | 9.88E-05 | 0.10 | 0.08 |
| **NC-044214.2-68300033-68300931** | *MTR* | 0 | NA | 0.28 | 9.89E-05 | 0.13 | 0.11 |
| **NC-044213.2-103638668-103639638** | *THOC1* | 0 | NA | 0.28 | 1.03E-04 | 0.10 | 0.08 |
| **NC-044221.2-24320352-24321308** | *SHOX2* | 36158 | NA | 0.26 | 1.05E-04 | 0.12 | 0.10 |
| **NC-044219.2-5126225-5127158** | *SEC22A* | 0 | NA | 0.28 | 1.06E-04 | 0.14 | 0.11 |
| **NC-044225.2-16421781-16422576** | *PCYOX1L* | 737 | Protein coding | 0.25 | 1.13E-04 | 0.15 | 0.12 |
| **NC-044214.2-29720166-29721081** | *RIMS1* | 5797 | NA | 0.26 | 1.16E-04 | 0.15 | 0.12 |
| **NC-044231.2-105894-106838** | *TXNDC17* | 0 | NA | 0.25 | 1.29E-04 | 0.13 | 0.11 |
| **NC-044218.2-23076933-23077880** | *TRIM8* | 3653 | NA | 0.28 | 1.31E-04 | 0.10 | 0.08 |
| **NC-044225.2-9342895-9343832** | *SMAD5* | 24567 | Protein coding | 0.31 | 1.33E-04 | 0.12 | 0.09 |
| **NC-044218.2-3684722-3685533** | *RNLS* | 0 | NA | 0.36 | 1.41E-04 | 0.10 | 0.08 |
| **NC-044217.2-34253400-34254335** | *FOXG1* | 0 | NA | 0.32 | 1.54E-04 | 0.13 | 0.10 |
| **NC-054768.1-466870-467783** | *ACIN1* | 0 | NA | 0.26 | 1.85E-04 | 0.14 | 0.11 |
| **NC-044218.2-31337036-31337945** | *HTRA1* | 0 | NA | 0.29 | 2.23E-04 | 0.10 | 0.08 |
| **NC-044213.2-66432675-66433555** | *GFOD1* | 0 | NA | 0.34 | 2.25E-04 | 0.11 | 0.08 |
| **NC-044211.2-97726002-97727600** | *NHS* | 0 | NA | 0.28 | 2.43E-04 | 0.12 | 0.09 |
| **NC-044214.2-99684811-99685702** | *HHAT* | 0 | NA | 0.26 | 2.60E-04 | 0.13 | 0.11 |
| **NC-044217.2-54934399-54935336** | *SYNE2* | 0 | NA | 0.26 | 2.73E-04 | 0.11 | 0.09 |
| **NC-044240.2-1178933-1179566** | *MIDN* | 0 | NA | 0.30 | 2.80E-04 | 0.11 | 0.08 |
| **NC-044212.2-64603372-64604286** | *ARFGAP3* | 0 | NA | 0.30 | 2.89E-04 | 0.10 | 0.08 |
| **NC-044211.2-75104859-75105803** | *TM9SF2* | 0 | NA | 0.27 | 3.29E-04 | 0.14 | 0.11 |
| **NC-044217.2-22751705-22752578** | *LOC100220107* | 0 | NA | 0.25 | 4.44E-04 | 0.11 | 0.09 |
| **NC-044221.2-11187499-11188338** | *LOC100228212* | 0 | NA | 0.26 | 4.68E-04 | 0.11 | 0.09 |
| **NC-044213.2-47076225-47077131** | *KIAA0895* | 0 | NA | 0.31 | 5.30E-04 | 0.12 | 0.09 |

| Peak | Gene | Distance | Gene Biotype | Average log2FC | FDR | Fraction 1 | Fraction 2 |
|---|---|---|---|---|---|---|---|
| **NC-044223.2-17516038-17516968** | *LOC121470584* | 0 | NA | 0.29 | 5.74E-04 | 0.11 | 0.08 |
| **NC-044233.2-6824123-6825043** | *PINK1* | 0 | NA | 0.31 | 6.26E-04 | 0.11 | 0.08 |
| **NC-044226.2-4953416-4954318** | *FBXL18* | 0 | NA | 0.29 | 6.60E-04 | 0.12 | 0.09 |
| **NC-044212.2-53435509-53436415** | *STAB2* | 6030 | Protein coding | 0.26 | 7.88E-04 | 0.10 | 0.08 |
| **NC-044217.2-51960526-51961360** | *TRMT61A* | 0 | NA | 0.27 | 8.07E-04 | 0.10 | 0.08 |
| **NC-044212.2-24016133-24017069** | *ST7* | 0 | NA | 0.26 | 8.26E-04 | 0.13 | 0.10 |
| **NC-044238.2-3257008-3257981** | *LOC121471158* | 0 | NA | 0.27 | 8.42E-04 | 0.11 | 0.09 |
| **NC-044217.2-56884038-56884920** | *DAAM1* | 0 | NA | 0.33 | 9.45E-04 | 0.11 | 0.09 |
| **NC-044230.2-10807732-10808604** | *DGKE* | 0 | NA | 0.27 | 9.83E-04 | 0.11 | 0.09 |
| **NC-044214.2-44228655-44229614** | *NR2E1* | 0 | NA | 0.29 | 1.17E-03 | 0.11 | 0.09 |
| **NC-044215.2-49298431-49299310** | *TSPAN5* | 0 | NA | 0.27 | 1.32E-03 | 0.12 | 0.09 |
| **NC-044216.2-1761611-1762566** | *LOC115495205* | 6390 | lncRNA | 0.27 | 1.38E-03 | 0.11 | 0.09 |
| **NC-044223.2-10895968-10896882** | *ZC3H18* | 0 | NA | 0.28 | 1.39E-03 | 0.10 | 0.08 |
| **NC-044222.2-17311187-17312123** | *ADAMTS17* | 0 | NA | 0.27 | 1.40E-03 | 0.10 | 0.08 |
| **NC-044219.2-3088353-3089280** | *USP40* | 0 | NA | 0.26 | 1.42E-03 | 0.10 | 0.08 |
| **NC-044217.2-54706949-54707911** | *HSPA2* | 86 | NA | 0.34 | 1.48E-03 | 0.12 | 0.09 |
| **NC-044215.2-66585344-66586234** | *SHROOM3* | 0 | NA | 0.26 | 1.53E-03 | 0.10 | 0.08 |
| **NC-044211.2-85600961-85601901** | *LOC105758645* | 0 | NA | 0.27 | 1.81E-03 | 0.13 | 0.10 |
| **NC-044212.2-49656143-49657045** | *C1AH22orf23* | 0 | NA | 0.33 | 1.84E-03 | 0.10 | 0.08 |
| **NC-044220.2-10241771-10242733** | *ARTN* | 994 | Protein coding | 0.29 | 2.04E-03 | 0.12 | 0.10 |
| **NC-044212.2-12109758-12110661** | *ARMC10* | 0 | NA | 0.28 | 2.05E-03 | 0.11 | 0.09 |
| **NC-044232.2-4106173-4107104** | *MYBL2* | 0 | NA | 0.30 | 2.16E-03 | 0.11 | 0.08 |
| **NC-044215.2-6286489-6287435** | *FGFRL1* | 17832 | NA | 0.30 | 2.21E-03 | 0.11 | 0.09 |
| **NC-044215.2-68375467-68376326** | *PCDH18* | 0 | NA | 0.25 | 2.29E-03 | 0.11 | 0.09 |
| **NC-044213.2-113506499-113507377** | *FAM110B* | 0 | NA | 0.25 | 2.31E-03 | 0.10 | 0.08 |
| **NC-044223.2-11250871-11251497** | *ANKRD11* | 0 | NA | 0.30 | 2.50E-03 | 0.13 | 0.11 |

| Peak | Gene | Distance | Gene Biotype | Average log2FC | FDR | Fraction 1 | Fraction 2 |
|---|---|---|---|---|---|---|---|
| **NC-044220.2-17183411-17184328** | *SLC44A3* | 5611 | Protein coding | 0.27 | 2.61E-03 | 0.10 | 0.08 |
| **NC-044241.2-14826600-14827439** | *FAM172A* | 0 | NA | 0.30 | 2.84E-03 | 0.11 | 0.09 |
| **NC-044224.2-3422171-3423157** | *LOC100219828* | 17432 | Protein coding | 0.36 | 3.05E-03 | 0.10 | 0.08 |
| **NC-044219.2-21589998-21590852** | *LOC121470299* | 0 | NA | 0.25 | 5.06E-03 | 0.14 | 0.11 |
| **NC-044224.2-5552177-5553167** | *LRIG1* | 24683 | Protein coding | 0.30 | 5.26E-03 | 0.11 | 0.08 |
| **NC-044240.2-2640845-2641845** | *LOC121471221* | 0 | NA | 0.30 | 5.39E-03 | 0.11 | 0.09 |
| **NC-044221.2-4612828-4613714** | *SRPRB* | 0 | NA | 0.25 | 7.33E-03 | 0.11 | 0.09 |
| **NC-044219.2-14653472-14654357** | *SLC25A12* | 0 | NA | 0.26 | 7.82E-03 | 0.12 | 0.09 |
| **NC-044222.2-2841633-2842385** | *LOC115496649* | 0 | NA | 0.25 | 9.11E-03 | 0.10 | 0.08 |
| **NC-044211.2-26745812-26746753** | *EPHA1* | 0 | NA | 0.26 | 9.46E-03 | 0.10 | 0.08 |
| **NC-044238.2-4204120-4205020** | *TSPAN2* | 0 | NA | 0.25 | 9.55E-03 | 0.10 | 0.08 |
| **NC-044213.2-139990241-139991154** | *TRIB1* | 0 | NA | 0.26 | 9.73E-03 | 0.10 | 0.08 |
| **NC-044217.2-38416467-38417365** | *LOC121470075* | 0 | NA | 0.28 | 1.00E-02 | 0.10 | 0.08 |
| **NC-054769.1-740051-740966** | *TSEN34* | 0 | NA | 0.34 | 1.21E-02 | 0.11 | 0.09 |
| **NC-044213.2-46982841-46983735** | *EEPD1* | 0 | NA | 0.25 | 1.25E-02 | 0.11 | 0.09 |
| **NC-044221.2-23882192-23883159** | *TRIM59* | 0 | NA | 0.25 | 1.31E-02 | 0.11 | 0.09 |
| **NC-044213.2-27770014-27770950** | *LOC121469368* | 23616 | NA | 0.30 | 1.56E-02 | 0.10 | 0.08 |
| **NC-044220.2-24137861-24138830** | *SWT1* | 0 | NA | 0.26 | 2.01E-02 | 0.12 | 0.10 |
| **NC-044230.2-2107848-2108846** | *LOC115497647* | 0 | NA | 0.30 | 2.21E-02 | 0.12 | 0.10 |
| **NC-044222.2-4314444-4315405** | *VPS13C* | 0 | NA | 0.27 | 2.69E-02 | 0.12 | 0.10 |
| **NC-044215.2-49393082-49394046** | *RAP1GDS1* | 0 | NA | 0.35 | 3.59E-02 | 0.10 | 0.08 |
| **NC-044214.2-29239506-29240513** | *OGFRL1* | 0 | NA | 0.26 | 4.10E-02 | 0.11 | 0.09 |
| **NC-044214.2-65807388-65808323** | *SASH1* | 0 | NA | 0.28 | 4.15E-02 | 0.10 | 0.08 |
| **NC-044214.2-105850456-105851400** | *LOC100218075* | 14235 | Protein coding | 0.28 | 4.26E-02 | 0.10 | 0.08 |

**Supplemental Table 13: Rewired and differentially accessible chromatin astrocyte peaks in heat call embryos.** List of 57 "dual-hit" regulatory peaks in astrocytes that (i) were differentially accessible in Heat Call versus Control playback (FindMarkers on ATAC peaks; MAST; two-sided; Benjamini–Hochberg FDR) and (ii) also served as endpoints of Heat Call-specific rewired Cicero co-accessibility connections (co-accessibility ≥0.15; active in Heat Call with both endpoints accessible in ≥5% of astrocytes and inactive in Control with ≥1 endpoint <5% accessible). Peaks are reported as genomic coordinates (peak) with associated differential accessibility statistics: P-Value (unadjusted P value), Average log2FC (log2 fold-change in peak accessibility, Heat Call vs Control), Fraction 1 and Fraction 2 (fractions of astrocytes with the peak accessible in Heat Call and Control, respectively), and FDR (BH-adjusted FDR). Sample size: Control n = 4 pooled biological replicates and Heat Call n = 4 pooled biological replicates (five hypothalamic tissue punches per pool; embryos collected at E13), comprising 55,950 total nuclei and 14,949 astrocyte nuclei used for the astrocyte-specific analyses.

| Peak | P-Value | FDR | Average log2FC | Fraction 1 | Fraction 2 |
|---|---|---|---|---|---|
| **NC-044232.2-12335224-12336167** | **7.82E-17** | 6.15E-12 | 0.31 | 0.19 | 0.15 |
| **NC-044221.2-2356666-2357648** | **2.64E-15** | 2.07E-10 | 0.41 | 0.11 | 0.08 |
| **NC-044213.2-4296865-4297771** | **8.48E-15** | 6.67E-10 | 0.29 | 0.21 | 0.17 |
| **NC-044212.2-53563924-53564841** | **1.07E-12** | 8.43E-08 | 0.28 | 0.15 | 0.12 |
| **NC-044217.2-6527953-6528913** | **1.20E-12** | 9.41E-08 | 0.29 | 0.10 | 0.08 |
| **NC-044211.2-56143572-56144485** | **1.28E-12** | 1.00E-07 | 0.26 | 0.12 | 0.10 |
| **NC-044219.2-19333210-19334116** | **1.46E-12** | 1.15E-07 | 0.29 | 0.16 | 0.13 |
| **NC-044219.2-36304568-36305484** | **1.91E-12** | 1.51E-07 | 0.27 | 0.21 | 0.17 |
| **NC-044211.2-63720419-63721359** | **2.24E-12** | 1.76E-07 | 0.41 | 0.12 | 0.08 |
| **NC-044216.2-6978207-6979155** | **2.61E-12** | 2.06E-07 | 0.26 | 0.15 | 0.12 |
| **NC-044214.2-105427333-105428264** | **2.62E-12** | 2.06E-07 | 0.29 | 0.12 | 0.10 |
| **NC-044231.2-938966-939720** | **6.14E-12** | 4.82E-07 | 0.26 | 0.12 | 0.09 |
| **NC-044215.2-7182123-7183064** | **6.66E-12** | 5.23E-07 | 0.34 | 0.12 | 0.09 |
| **NC-044213.2-25564872-25565718** | **7.39E-12** | 5.81E-07 | 0.31 | 0.12 | 0.09 |
| **NC-044224.2-19530203-19531185** | **7.93E-12** | 6.24E-07 | 0.35 | 0.11 | 0.09 |
| **NC-044212.2-15149965-15150987** | **8.26E-12** | 6.49E-07 | 0.27 | 0.11 | 0.08 |
| **NC-044211.2-44450558-44451442** | **1.21E-11** | 9.52E-07 | 0.29 | 0.12 | 0.09 |
| **NC-044214.2-55417955-55418882** | **1.87E-11** | 1.47E-06 | 0.26 | 0.14 | 0.11 |

| Peak | P-Value | FDR | Average log2FC | Fraction 1 | Fraction 2 |
|---|---|---|---|---|---|
| **NC-044212.2-7068023-7069007** | **1.90E-11** | 1.50E-06 | 0.26 | 0.11 | 0.09 |
| **NC-044213.2-34507852-34508779** | **2.30E-11** | 1.81E-06 | 0.30 | 0.10 | 0.08 |
| **NC-044224.2-514937-515878** | **2.31E-11** | 1.81E-06 | 0.25 | 0.11 | 0.09 |
| **NC-044217.2-51953918-51954853** | **2.67E-11** | 2.10E-06 | 0.26 | 0.14 | 0.11 |
| **NC-044227.2-3475241-3476145** | **2.92E-11** | 2.29E-06 | 0.35 | 0.13 | 0.10 |
| **NC-044215.2-64617545-64618452** | **3.56E-11** | 2.80E-06 | 0.35 | 0.11 | 0.08 |
| **NC-044212.2-2459332-2460211** | **3.78E-11** | 2.97E-06 | 0.33 | 0.14 | 0.11 |
| **NC-044211.2-24329324-24330265** | **5.77E-11** | 4.54E-06 | 0.25 | 0.14 | 0.11 |
| **NC-044232.2-3099637-3100576** | **6.19E-11** | 4.87E-06 | 0.26 | 0.13 | 0.10 |
| **NC-044226.2-15205040-15206015** | **6.95E-11** | 5.46E-06 | 0.30 | 0.12 | 0.09 |
| **NC-044235.2-2973106-2974057** | **1.01E-10** | 7.96E-06 | 0.29 | 0.11 | 0.09 |
| **NC-044212.2-1944777-1945725** | **1.09E-10** | 8.61E-06 | 0.38 | 0.10 | 0.08 |
| **NC-044211.2-88931459-88932409** | **1.23E-10** | 9.63E-06 | 0.28 | 0.12 | 0.09 |
| **NC-044230.2-323222-324134** | **2.04E-10** | 1.60E-05 | 0.32 | 0.10 | 0.08 |
| **NC-044223.2-3950062-3951008** | **3.05E-10** | 2.40E-05 | 0.27 | 0.11 | 0.09 |
| **NC-044211.2-8450854-8451728** | **3.61E-10** | 2.84E-05 | 0.26 | 0.15 | 0.12 |
| **NC-044211.2-105604141-105605058** | **3.61E-10** | 2.84E-05 | 0.32 | 0.12 | 0.09 |
| **NC-044233.2-2821132-2822039** | **6.95E-10** | 5.46E-05 | 0.25 | 0.13 | 0.11 |
| **NC-044219.2-37542649-37543666** | **1.18E-09** | 9.30E-05 | 0.27 | 0.15 | 0.12 |
| **NC-044232.2-7216853-7217711** | **1.19E-09** | 9.33E-05 | 0.26 | 0.12 | 0.10 |
| **NC-044214.2-39916754-39917657** | **1.24E-09** | 9.77E-05 | 0.29 | 0.15 | 0.12 |
| **NC-044229.2-9338214-9339100** | **1.26E-09** | 9.88E-05 | 0.29 | 0.10 | 0.08 |
| **NC-044221.2-24320352-24321308** | **1.34E-09** | 1.05E-04 | 0.26 | 0.12 | 0.10 |
| **NC-044225.2-16421781-16422576** | **1.43E-09** | 1.13E-04 | 0.25 | 0.15 | 0.12 |
| **NC-044214.2-29720166-29721081** | **1.48E-09** | 1.16E-04 | 0.26 | 0.15 | 0.12 |
| **NC-044218.2-23076933-23077880** | **1.66E-09** | 1.31E-04 | 0.28 | 0.10 | 0.08 |
| **NC-044223.2-17516038-17516968** | **7.30E-09** | 5.74E-04 | 0.29 | 0.11 | 0.08 |

| Peak | P-Value | FDR | Average log2FC | Fraction 1 | Fraction 2 |
|---|---|---|---|---|---|
| **NC-044212.2-53435509-53436415** | **1.00E-08** | 7.88E-04 | 0.26 | 0.10 | 0.08 |
| **NC-044214.2-44228655-44229614** | **1.49E-08** | 1.17E-03 | 0.29 | 0.11 | 0.09 |
| **NC-044222.2-17311187-17312123** | **1.78E-08** | 1.40E-03 | 0.27 | 0.10 | 0.08 |
| **NC-044217.2-54706949-54707911** | **1.88E-08** | 1.48E-03 | 0.34 | 0.12 | 0.09 |
| **NC-044211.2-85600961-85601901** | **2.31E-08** | 1.81E-03 | 0.27 | 0.13 | 0.10 |
| **NC-044220.2-10241771-10242733** | **2.60E-08** | 2.04E-03 | 0.29 | 0.12 | 0.10 |
| **NC-044215.2-6286489-6287435** | **2.81E-08** | 2.21E-03 | 0.30 | 0.11 | 0.09 |
| **NC-044213.2-113506499-113507377** | **2.94E-08** | 2.31E-03 | 0.25 | 0.10 | 0.08 |
| **NC-044223.2-11250871-11251497** | **3.18E-08** | 2.50E-03 | 0.30 | 0.13 | 0.11 |
| **NC-044224.2-5552177-5553167** | **6.69E-08** | 5.26E-03 | 0.30 | 0.11 | 0.08 |
| **NC-044219.2-14653472-14654357** | **9.95E-08** | 7.82E-03 | 0.26 | 0.12 | 0.09 |
| **NC-044214.2-105850456-105851400** | **5.42E-07** | 4.26E-02 | 0.28 | 0.10 | 0.08 |

**Supplemental Table 14: Peak-to-gene links for dual-hit peaks (all tested pairs).** Table lists all peak–gene pairs tested by Signac LinkPeaks for the set of 57 “dual-hit” astrocyte peaks (DA-gained and participating in heat call-specific rewired connections), restricted to genes within 500 kb. For each pair, the Score is the Pearson correlation between peak accessibility and SCT-normalized gene expression across astrocyte nuclei, P-Value is the corresponding correlation P-value; Peak is reported as genomic coordinates.

| Peak | Gene | Score | P-Value |
|---|---|---|---|
| **NC-044212.2-2459332-2460211** | *PLXNA4* | 0.27 | 9.08E-11 |
| **NC-044215.2-7182123-7183064** | *FAM53A* | 0.24 | 8.45E-06 |
| **NC-044212.2-1944777-1945725** | *PLXNA4* | 0.19 | 1.21E-03 |
| **NC-044211.2-85600961-85601901** | *LOC105758648* | 0.16 | 1.38E-03 |
| **NC-044212.2-7068023-7069007** | *FAM107B* | 0.15 | 3.30E-10 |
| **NC-044212.2-53563924-53564841** | *ASCL1* | 0.14 | 3.56E-06 |
| **NC-044215.2-6286489-6287435** | *FGFRL1* | 0.13 | 7.32E-04 |
| **NC-044214.2-29720166-29721081** | *KCNQ5* | 0.13 | 1.12E-04 |
| **NC-044233.2-2821132-2822039** | *PRDM16* | 0.11 | 4.24E-04 |
| **NC-044215.2-7182123-7183064** | *TACC3* | 0.10 | 3.82E-05 |
| **NC-044215.2-7182123-7183064** | *LOC100227664* | 0.09 | 4.00E-06 |
| **NC-044212.2-1944777-1945725** | *NET1* | 0.09 | 2.38E-03 |
| **NC-044221.2-24320352-24321308** | *SMC4* | 0.09 | 1.75E-02 |
| **NC-044219.2-14653472-14654357** | *DLX2* | 0.08 | 3.79E-04 |
| **NC-044224.2-19530203-19531185** | *WNT7A* | 0.08 | 1.91E-04 |
| **NC-044212.2-53563924-53564841** | *LOC100227265* | 0.08 | 1.09E-03 |
| **NC-044213.2-4296865-4297771** | *CTDSPL* | 0.08 | 2.99E-04 |
| **NC-044231.2-938966-939720** | *MSI2* | 0.07 | 6.47E-03 |
| **NC-044212.2-7068023-7069007** | *GRM3* | -0.07 | 3.82E-02 |
| **NC-044222.2-17311187-17312123** | *ADAMTS17* | 0.07 | 6.53E-03 |
| **NC-044211.2-24329324-24330265** | *GPR156* | 0.07 | 1.17E-06 |
| **NC-044219.2-14653472-14654357** | *DLX1* | 0.07 | 1.31E-02 |
| **NC-044217.2-6527953-6528913** | *CCND1* | 0.07 | 2.94E-04 |
| **NC-044212.2-1944777-1945725** | *EXOC4* | 0.07 | 1.49E-05 |

| Peak | Gene | Score | P-Value |
|---|---|---|---|
| **NC-044211.2-24329324-24330265** | *POLQ* | 0.06 | 2.06E-03 |
| **NC-044219.2-14653472-14654357** | *SLC25A12* | 0.06 | 2.02E-03 |
| **NC-044212.2-53563924-53564841** | *LOC121469119* | 0.06 | 2.45E-04 |
| **NC-044219.2-36304568-36305484** | *RND3* | 0.06 | 3.17E-07 |
| **NC-044214.2-39916754-39917657** | *POU3F2* | 0.06 | 2.70E-05 |
| **NC-044212.2-2459332-2460211** | *LOC121471101* | 0.06 | 4.93E-07 |
| **NC-044213.2-34507852-34508779** | *CREB5* | 0.06 | 4.29E-03 |
| **NC-044211.2-85600961-85601901** | *LOC115497607* | 0.06 | 3.10E-04 |
| **NC-044232.2-7216853-7217711** | *TPX2* | 0.05 | 2.62E-02 |
| **NC-044219.2-14653472-14654357** | *METAP1D* | 0.05 | 3.70E-02 |
| **NC-044233.2-2821132-2822039** | *LOC116809204* | 0.05 | 1.36E-03 |
| **NC-044216.2-6978207-6979155** | *GPC4* | 0.05 | 5.13E-04 |
| **NC-044224.2-19530203-19531185** | *IQSEC1* | 0.05 | 1.17E-02 |
| **NC-044212.2-53435509-53436415** | *ASCL1* | 0.05 | 3.16E-02 |
| **NC-044211.2-85600961-85601901** | *POU3F3* | 0.05 | 1.00E-02 |
| **NC-044233.2-2821132-2822039** | *LOC121470970* | 0.05 | 7.46E-04 |
| **NC-044212.2-53435509-53436415** | *LOC121469119* | 0.05 | 1.09E-03 |
| **NC-044211.2-56143572-56144485** | *RGCC* | 0.05 | 1.87E-07 |
| **NC-044214.2-44228655-44229614** | *NR2E1* | 0.05 | 1.71E-03 |

**Supplemental Table 15: Significant pseudotime shifts by cluster.** Clusters showing significant differences in pseudotime between control and heat Call conditions (FDR < 0.05; Wilcoxon rank-sum test) from Slingshot-based trajectory analysis of all hypothalamic cell types. For each WNN cluster, the table reports the cluster ID, cell type annotation, number of cells per condition (N Control, N Heat Call), mean pseudotime in each condition (Mean PT Control, Mean PT Heat Call), the difference in mean pseudotime (Delta PT (Heat Call − Control), nominal P value (Wilcoxon P), and FDR.

| Cluster | Annotation | N Control | N Heat Call | Mean PT Control | Mean PT Heat Call | Delta PT | Wilcoxon P | FDR |
|---|---|---|---|---|---|---|---|---|
| **1** | Astrocyte | 1474 | 49 | 3.3 | 21.67 | 18.37 | 2.05E-32 | 4.72E-32 |
| **11** | Astrocyte | 758 | 3 | 8.82 | 24.62 | 15.8 | 2.78E-03 | 4E-03 |
| **6** | GABAergic Neuron | 8 | 5 | 9.51 | 18.31 | 8.8 | 1.36E-02 | 1.85E-02 |
| **10** | GABAergic Neuron | 720 | 758 | 7.56 | 15.65 | 8.09 | 1.43E-241 | 1.1E-240 |
| **23** | Astrocyte | 386 | 4 | 17.38 | 23.89 | 6.51 | 5.82E-04 | 8.93E-04 |
| **13** | GABAergic Neuron | 698 | 666 | 7.14 | 13.57 | 6.43 | 6.49E-257 | 7.46E-256 |
| **20** | Astrocyte | 388 | 12 | 18.79 | 25.19 | 6.4 | 3.62E-09 | 7.56E-09 |
| **18** | GABAergic Neuron | 9 | 6 | 7.12 | 12.95 | 5.83 | 4.2E-04 | 6.9E-04 |
| **24** | GABAergic Neuron | 383 | 19 | 7.12 | 12.68 | 5.56 | 4.11E-89 | 1.58E-88 |
| **3** | Glutamatergic/GABA | 677 | 114 | 7.12 | 11.62 | 4.49 | 1.9E-148 | 8.73E-148 |
| **5** | Astrocyte | 1046 | 2 | 3.69 | 8.12 | 4.43 | 2.23E-02 | 2.56E-02 |
| **41** | Astrocyte | 109 | 155 | 20.28 | 24.31 | 4.03 | 1.7E-43 | 4.9E-43 |
| **22** | Oligodendrocyte | 104 | 117 | 18.77 | 22.6 | 3.84 | 1.23E-37 | 3.15E-37 |
| **31** | Glutamatergic Neuron | 7 | 28 | 7.12 | 10.96 | 3.84 | 5.41E-05 | 1.04E-04 |
| **2** | Astrocyte | 1428 | 1519 | 21.21 | 24.75 | 3.54 | 0E+00 | 0E+00 |
| **21** | Microglia | 532 | 503 | 26.26 | 29.63 | 3.38 | 1.56E-170 | 8.95E-170 |
| **30** | Astrocyte | 270 | 257 | 23.56 | 26.77 | 3.21 | 9.83E-88 | 3.23E-87 |
| **8** | Glutamatergic Neuron | 2 | 185 | 14.26 | 3.32 | -10.94 | 1.71E-02 | 2.07E-02 |
| **12** | Glutamatergic Neuron | 5 | 387 | 21.4 | 9.44 | -11.96 | 1.32E-04 | 2.34E-04 |
| **7** | Glutamatergic Neuron | 2 | 735 | 19.21 | 5.18 | -14.03 | 1.48E-02 | 1.9E-02 |

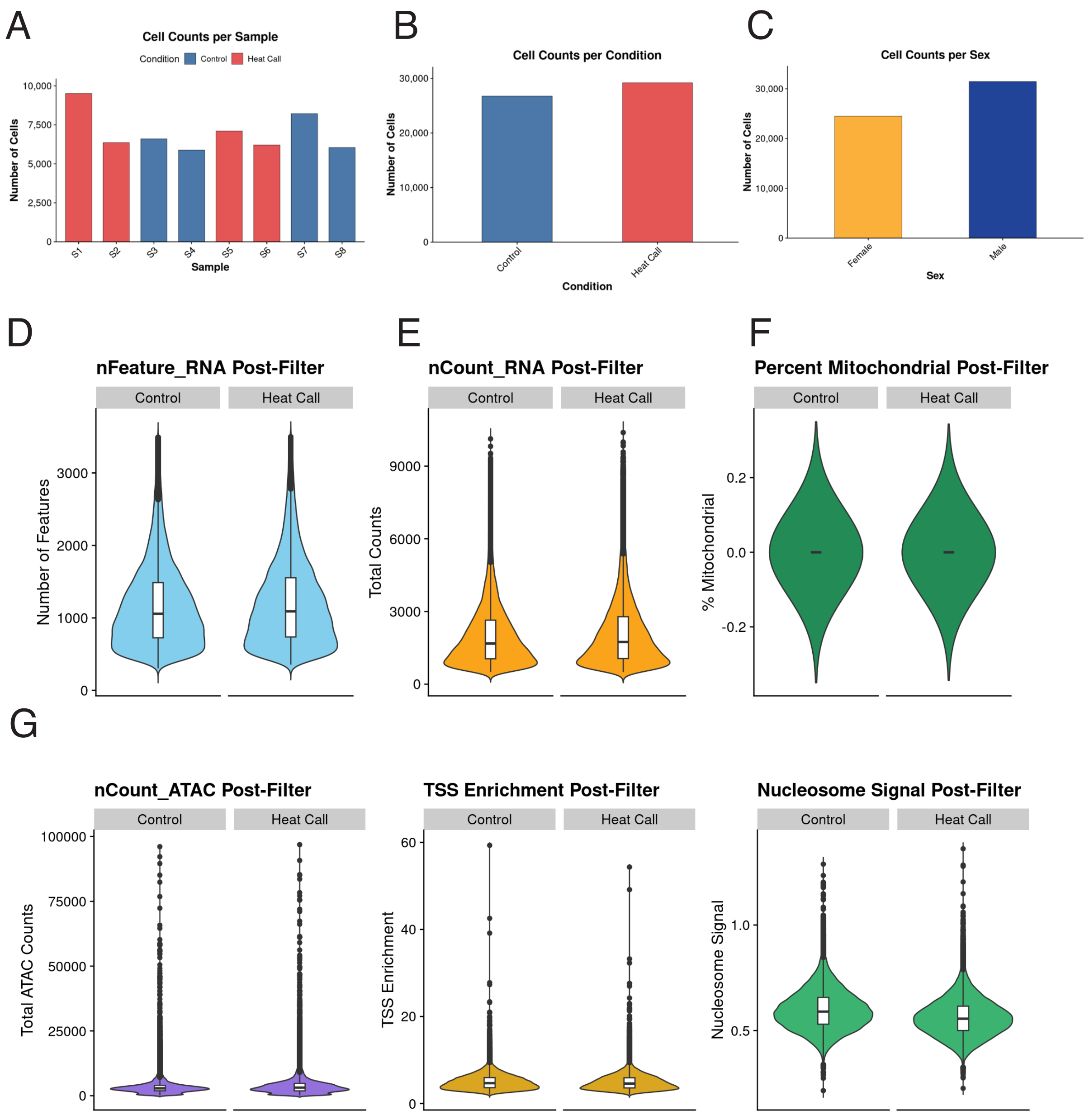
A
Cell Counts per Sample
Condition Control Heat Call
Number of Cells
10,000
7,500
5,000
2,500
0
S1 S2 S3 S4 S5 S6 S7 S8
Sample
B
Cell Counts per Condition
Number of Cells
30,000
20,000
10,000
0
Control
Heat Call
Condition
C
Cell Counts per Sex
Number of Cells
30,000
20,000
10,000
0
Female
Male
Sex
D
nFeature_RNA Post-Filter
Control
Heat Call
Number of Features
3000
2000
1000
0
E
nCount_RNA Post-Filter
Control
Heat Call
Total Counts
9000
6000
3000
0
F
Percent Mitochondrial Post-Filter
Control
Heat Call
% Mitochondrial
0.2
0.0
-0.2
G
nCount_ATAC Post-Filter
Control
Heat Call
Total ATAC Counts
100000
75000
50000
25000
0
TSS Enrichment Post-Filter
Control
Heat Call
TSS Enrichment
60
40
20
0
Nucleosome Signal Post-Filter
Control
Heat Call
Nucleosome Signal
1.0
0.5

**Supplemental Figure 1: Single-Nucleus RNA/ATAC Sequencing Quality Control.** Post-QC distributions of nuclei counts (A–C), RNA quality metrics (D–F), and ATAC quality metrics (G) from single-nucleus multiome sequencing (10x Chromium ATAC + Gene Expression) of E13 zebra finch hypothalamus. (A) Nuclei counts per sample (n = 8 pools). (B) Nuclei per condition (Control n = 4 pools; Heat Call n = 4 pools). (C) Nuclei per sex (Female n = 4 pools; Male n = 4 pools). (D–F) Post-filter distributions of the number of unique genes detected (nFeature_RNA), total number of RNA molecules (nCount_RNA), and fraction of RNA counts from mitochondrial genes (percent mitochondrial) per nucleus. (G) Post-filter distributions of ATAC quality metrics: total number of ATAC fragments per nucleus (nCount_ATAC), nucleosome signal, and TSS enrichment score. Each pool contained five hypothalamic tissue punches. Final dataset: 55,950 nuclei passed QC (57,388 RNA; 70,720 ATAC). Violin plots show median (center line) and interquartile range (box).

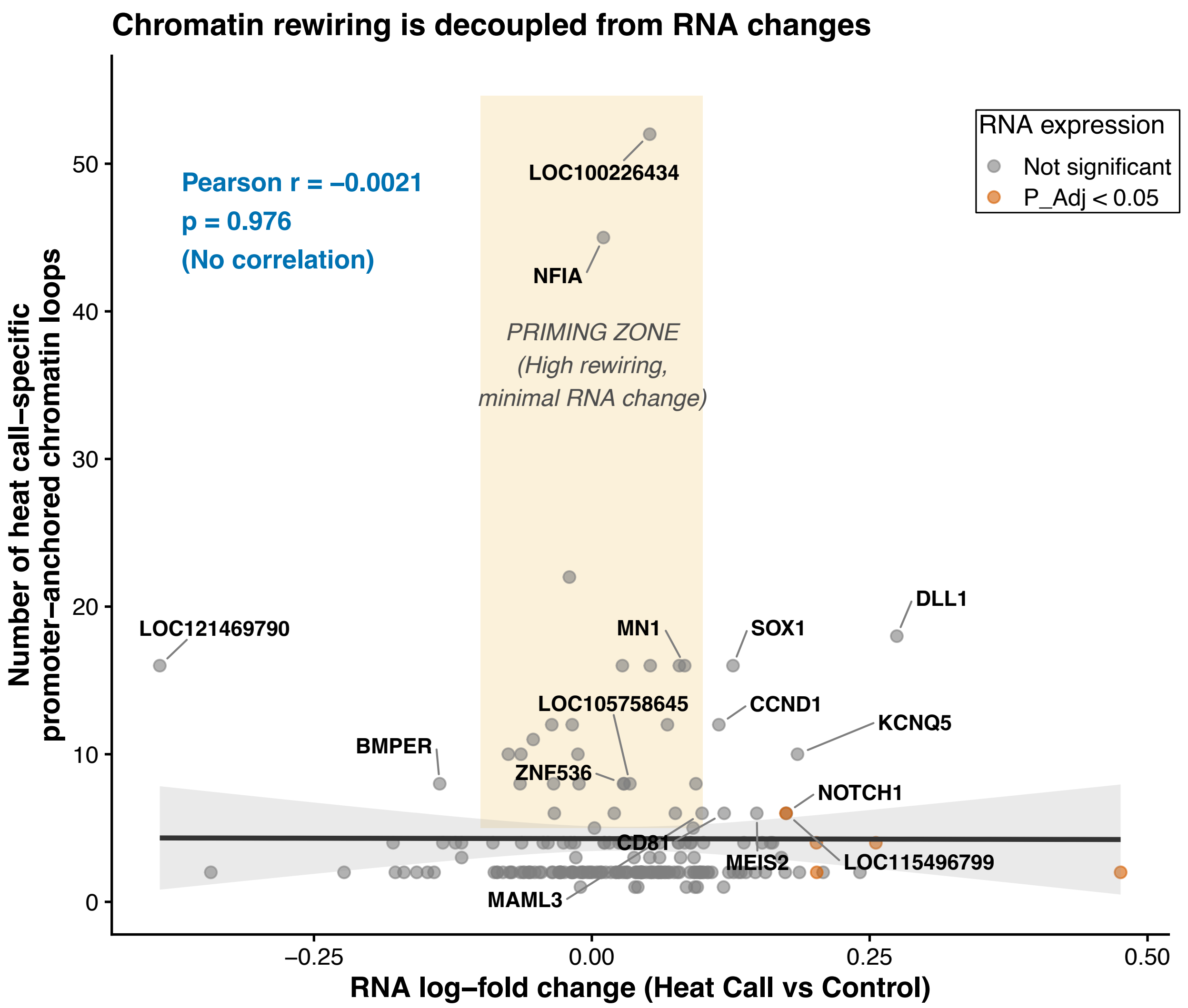


**Supplemental Figure 2: Chromatin rewiring is decoupled from RNA changes in astrocytes.** Scatter plot of RNA log fold-change (Heat Call vs Control; x-axis) versus the number of heat call-specific promoter-anchored chromatin loops per gene (y-axis; promoters defined as ±2 kb from TSS; Cicero co-accessibility ≥0.15, loops active in heat call when both endpoints accessible in ≥5% of astrocytes and inactive in Control when at least one endpoint accessible in <5% of astrocytes). Point color indicates RNA differential expression significance (MAST, two-sided; Benjamini–Hochberg FDR; $P_{adj} < 0.05$). Pearson correlation is shown ($r = -0.0021$, $p = 0.976$). Sample size: Control n = 4 pooled biological replicates and heat call n = 4 pooled biological replicates (five hypothalamic punches per pool; E13 embryos), comprising 55,950 total nuclei and 14,949 astrocyte nuclei.

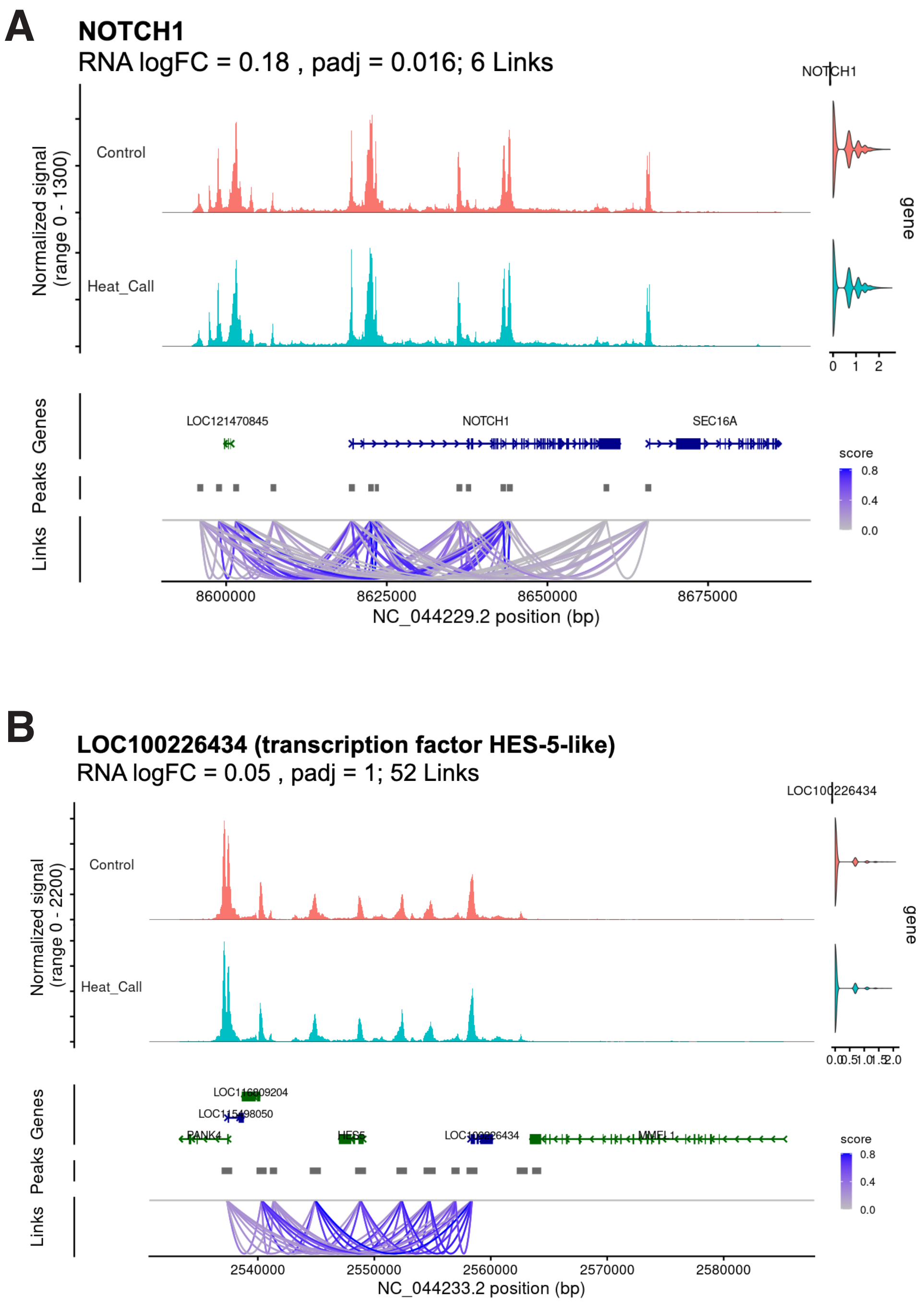
A
NOTCH1
RNA logFC = 0.18 , padj = 0.016; 6 Links
NOTCH1
Normalized signal
(range 0 - 1300)
Control
Heat_Call
gene
0
1
2
Genes
Peaks
Links
LOC121470845
NOTCH1
SEC16A
score
0.8
0.4
0.0
8600000
8625000
8650000
8675000
NC_044229.2 position (bp)
B
LOC100226434 (transcription factor HES-5-like)
RNA logFC = 0.05 , padj = 1; 52 Links
LOC100226434
Normalized signal
(range 0 - 2200)
Control
Heat_Call
gene
Genes
Peaks
Links
score
0.8
0.4
0.0
2540000
2550000
2560000
2570000
2580000
NC_044233.2 position (bp)

**Supplemental Figure 1: Heat call–specific chromatin rewiring at Notch signaling genes in astrocytes.** Chromatin accessibility (snATAC-seq coverage tracks) and co-accessibility links (arcs) at (A) *NOTCH1* and (B) *LOC100226434* (HES-5-like) loci in embryonic day 13 zebra finch hypothalamic astrocytes. Arcs represent heat call–specific rewired connections (Cicero co-accessibility ≥0.15; both peaks accessible in ≥5% of Heat Call astrocytes; at least one peak <5% accessible in Control astrocytes). Arc color intensity indicates co-accessibility score. Right: violin plots show per-nucleus gene expression distributions by condition. RNA log fold-changes (Heat Call vs Control) and adjusted P values are from MAST differential expression testing with Benjamini–Hochberg correction. *NOTCH1*: RNA logFC = 0.18, padj = 0.016, 6 heat call–specific links. *LOC100226434*: RNA logFC = 0.05, padj = 1.0, 52 heat call–specific links. Sample size: n = 4 biological replicates per condition (pooled samples, 5 tissue punches each), n = 14,949 total astrocyte nuclei, n = 55,950 total nuclei across all cell types.

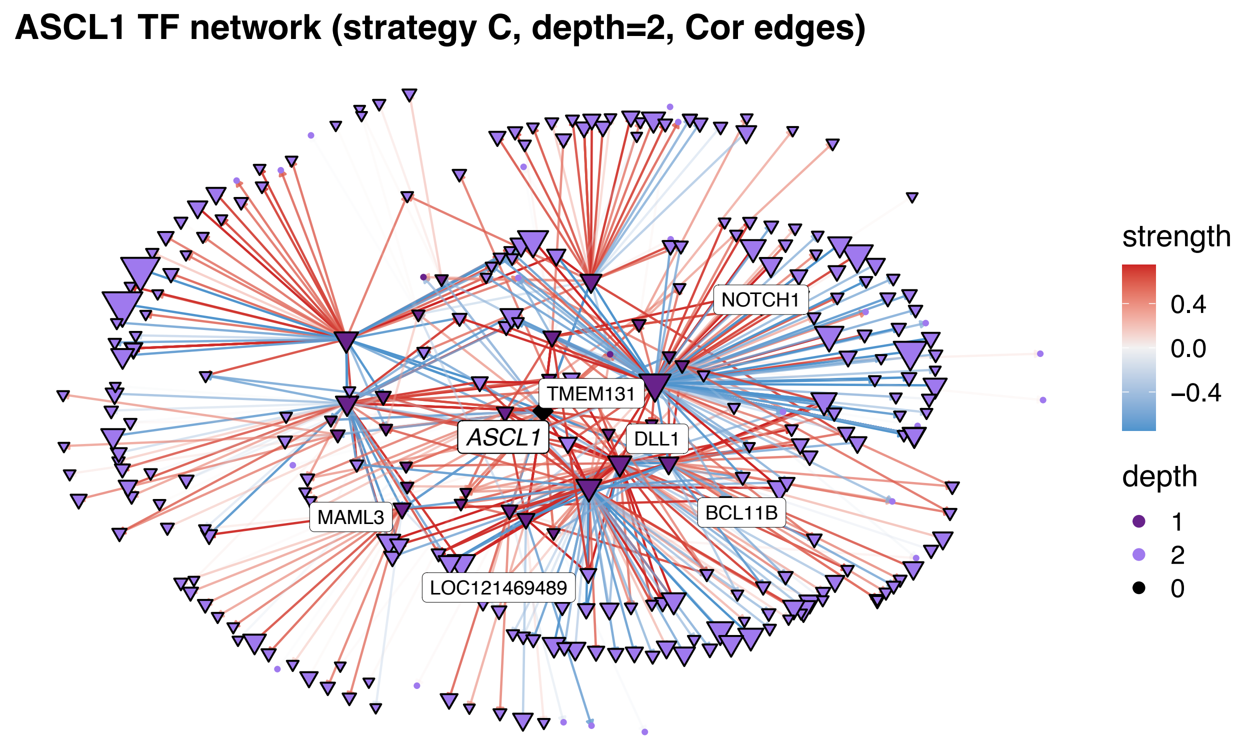


**Supplemental Figure 4: *ASCL1* transcription factor regulatory network in astrocytes:** Regulatory network showing *ASCL1* and its target genes in heat call-exposed astrocytes (n=14,949 nuclei from n=4 pooled samples, where each pool contains 5 E13 medial hypothalamic punches). Network inferred using XGBoost-based framework with motif-informed regulon assignment (strategy C, regulatory gain threshold=0.005, depth=2). Nodes represent transcription factors (labeled) and target genes (triangles), with node size indicating network depth (0=*ASCL1*, 1=direct targets, 2=secondary targets). Edge color represents correlation strength between TF and target expression: red=positive correlation, blue=negative correlation, gray=weak correlation. Edge thickness represents regulatory gain (XGBoost importance score). Key Notch pathway components (*DLL1, NOTCH1, MAML3*) emerge as *ASCL1* network neighbors, consistent with coordinated regulation of gliogenic programs.
